# Sinoatrial node impulses emerge from incessant, unique, kinetic transitions in disordered local calcium dynamics

**DOI:** 10.1101/2024.12.18.629224

**Authors:** Syevda Tagirova, Alexander V. Maltsev, Georgiana L. Baca, Victor A. Maltsev, Rostislav Bychkov, Edward G. Lakatta

## Abstract

We used both linear and nonlinear analyses to determine how information processing within and among incessant heterogeneous local Ca^2+^ oscillation (LCO) results in formation of rhythmic impulses in mouse SAN *ex vivo*.

Phase analysis delineated a network of functional pacemaker cell clusters, distinguished by their LCO amplitudes, kinetics, and phases. Cross-talk of LCO dynamics within the network culminated in rhythmic SAN global Ca^2+^ transients (CaTs), having a mean rate and rhythm identical to that recorded by the reference sharp electrode in the right atria, indicating that CaTs are induced by global SAN electrical impulses. Initial conditions of each impulse and subsequent LCO ensemble evolution during an impulse differed from each other, associated with an apparent stochastic process (carrying a degree of uncertainty) within network. A small pacemaker cluster located near the crista terminalis (CT) exhibited the highest degrees of intrinsic power, earliest rotor-like energy transfer, most frequent point-to-point instability, earliest acrophase, greatest impulse-to-impulse variability within the SAN functional cluster network. Unique, variable small-world network Ca^2+^ information sharing within and among *all* clusters during initial and terminal impulse phases*, created a unique solution* for each impulse, while preserving the identity of each cluster (a highly efficient form of information processing at low wiring and energy costs). Cross-recurrence analysis verified that LCO dynamics within the small cluster near CT were more stochastic and less deterministic than those of the other clusters, indicating that this small cluster took the lead in the initiation of the SAN impulse and that the others followed.

## 1. Introduction

Incessant, stochastic, oscillatory calcium (Ca^2+^) signals underlie important biologic processes throughout nature, and it has been argued that oscillatory frequency-dependent amplitude control provides a precise, noise-resistant control strategy that allows functional Ca^2+^ biologic signals that are of graded amplitude to be stable[1, 2]. For example, self-organization of local Ca^2+^ dynamics in the brainstem are implicated in initiation of signals that regulate rhythmic automatic breathing,[3] in the pancreas with respect to the regulation of insulin secretion[4], and in melanocyte initiation of skin protective signals[5].

Rhythmic Ca^2+^ dynamics initiate the first heartbeat[6] and the emergence of spiking regimes in the early heartbeat are driven by a noisy phase transition in which gradual and largely asynchronous development of Ca^2+-^dependent single-cell bioelectrical properties produce a stereotyped and robust tissue-scale transition from quiescence to coordinated beating[7]. In cells isolated from the adult heart’s pacemaker, the sinoatrial node (SAN), frequency-dependent amplitude control of spontaneous ensemble Ca^2+^ signals formed by spontaneous local Ca^2+^ releases (oscillations) allows for rapid, graded control of the action potential (AP) firing rate[8–15].

Pacemaker cells embedded in SAN tissue, incessantly generate spontaneous, partially stochastic,[16] local Ca^2+^ oscillations (LCO) that are heterogeneous in phase, frequency and amplitude throughout the SAN (see Video S1)[17].

The quintessential research frontier in SAN impulse formation is to understand how information encoded within local Ca^2+^ oscillations (LCOs) that are heterogeneous within and among pacemaker cells across SAN tissue is translated into formation of rhythmic SAN impulses that initiate rhythmic impulses that emerge within the SAN to initiate heartbeats[8, 17, 18]. Studies using confocal 3D tile imaging coupled to intracellular Ca^2+^ measurements have detailed novel cytoarchitecture of the network of nerves, glial and interstitial cells and pacemaker cells within the SAN[17, 18]. A novel approach (chronopix analysis) to detect phase heterogeneity of Ca^2+^ signaling within image pixels demonstrated that, in antithesis to a concentric, continuous spread of impulse formation,[19–21] the spread of Ca^2+^ signals is discontinuous throughout the SAN.

It has been suggested that SAN cytoarchitecture harboring Ca^2+^ oscillations resembles neuronal tissue [8, 17, 18]. In neuronal tissue the LCO information sharing underpins neural communication and plasticity, from modulation of neurotransmitter release to the activation of gene transcription essential for long-term potentiation and memory formation[1]. It is known that the role of microdomains within neurons, where Ca^2+^ concentrations can rapidly oscillate, underscore the precision with which Ca^2+^ signaling must be regulated to affect neuronal function accurately[1, 22–24].

The discovery of small-world network topology in the brain provided insights into neural organization. These networks are characterized by high local clustering coefficients (similar to regular lattices) while maintaining short average path lengths (similar to random networks) through the presence of select long-range connections. This architecture allows for both segregated and integrated information processing[25–27].

Inspired by previous studies of how spontaneous local Ca^2+^ dynamics participate in the formation of APs in single SAN cells[8–15], and the idea that synchronization of incessant heterogeneous Ca^2+^ oscillations is a universalsignaling mechanism at multiple scales [2, 28], we applied, for the first time in ex vivo SAN tissue, a wide range of analytic strategies adapted from dynamical systems and neuroscience to SAN Ca^2+^ dynamics. Our goal was to define how rhythmic global SAN tissue Ca^2+^ impulses emerge from incessant stochastic, heterogenous LCOs in pacemaker cells.

## 2. Materials and Methods

### 2.1. Sinoatrial node tissue (SAN) preparation

Our experiments conformed to the Guide for the Care and Use of Laboratory Animals, published by the US National Institutes of Health. The experimental protocols were approved by the Animal Care and Use Committee of the National Institutes of Health (protocol # 457-LCS-2027). See details in supplementary materials.

### 2.2. Data acquisition of local Ca^2+^ signals/oscillations (LCO) during impulse formation in SAN ex vivo

The SAN preparations (n=4) were incubated with Calbryte 520 AM (25 μM) a fluorescent calcium indicator, for 1.5 hours at room temperature. See detailed methods in supplementary materials.

### 2.3. Atrial cell action potential recordings

Because action potential (AP) characteristics differ in the SAN pacemaker cells among regions of the SAN [17], we placed a sharp electrode in the atria near the crista terminalis (CT) to deduce the timing of SAN action potential (AP) with reference to atrial excitation. See detailed methods in supplementary materials.

### 2.4. Immunolabeling

The SAN preparations were washed three times in phosphate-buffered saline (PBS) and permeabilized overnight in PBS containing 0.2% Triton X-100 and 20% DMSO. After blocking the nonspecific binding sites by incubation for 8 hours with 0.2% Tween-20 in PBS containing 3% normal donkey serum, SAN whole-mount preparations were incubated for 3 days with the primary antibodies diluted in PBS containing 0.2% Tween-20 and 3% normal donkey serum, and then washed three times with PBS containing 0.2% Tween-20, incubated overnight with appropriate secondary antibodies then washed three times with PBS containing 0.2% Tween-20. Immunolabeling of whole-mount SANs was imaged with a ZEISS-LSM800 confocal microscope. Antibodies: HCN4+ cells were identified by rabbit polyclonal antibodies for hyperpolarization-activated, cyclic nucleotide-gate cation channels HCN4 (1:300; Alomone Labs). Mouse monoclonal antibody to connexin 43 (1:300; Invitrogen) was used to label the gap junctions of striated cells.

### 2.5. Data analysis

In the present study, we used multiple software programs to apply a diverse set of linear and nonlinear analytical analyses of local Ca^2+^ (LCO) dynamics. For additional details see the supplementary materials.

## 3. Glossary of Terms

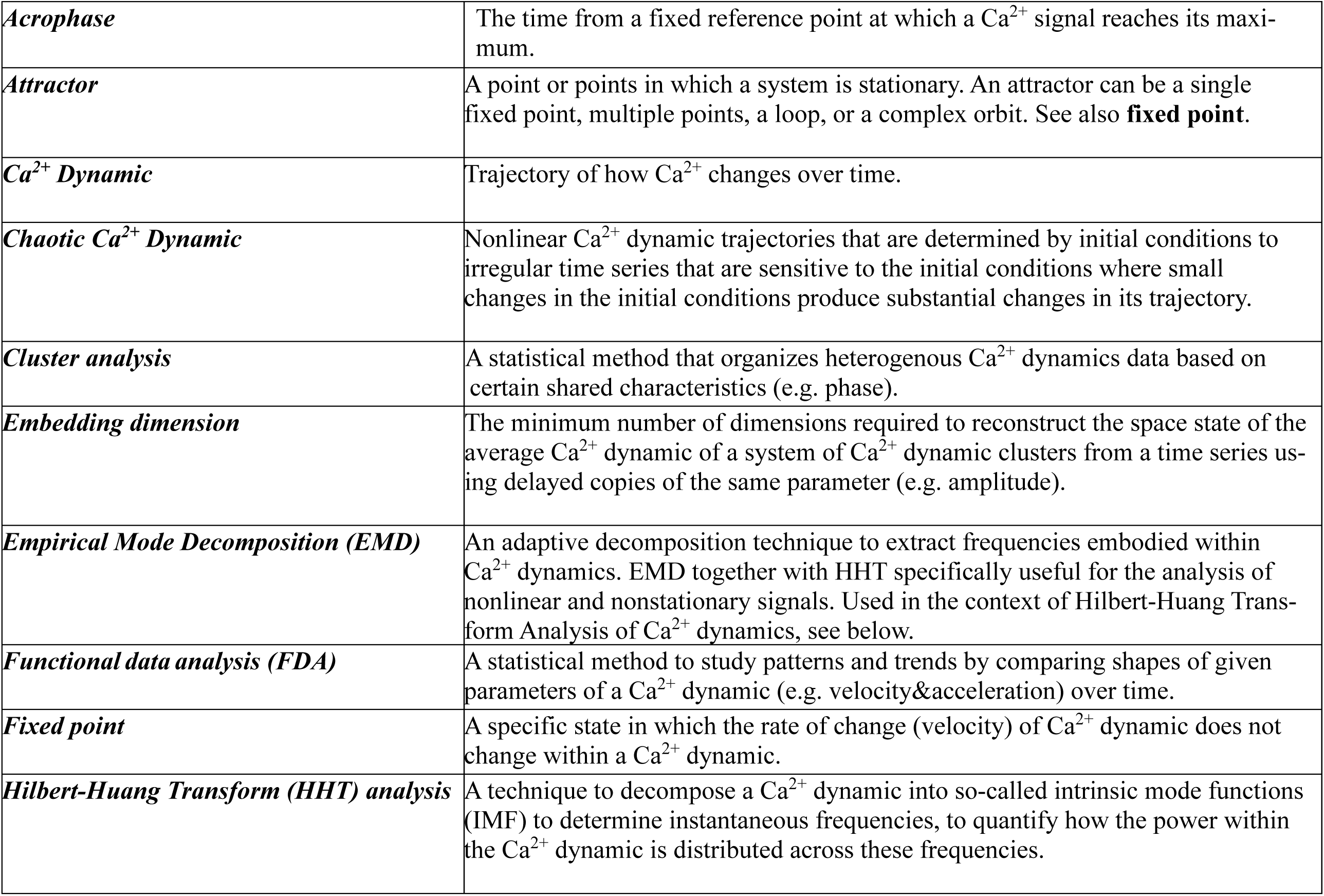
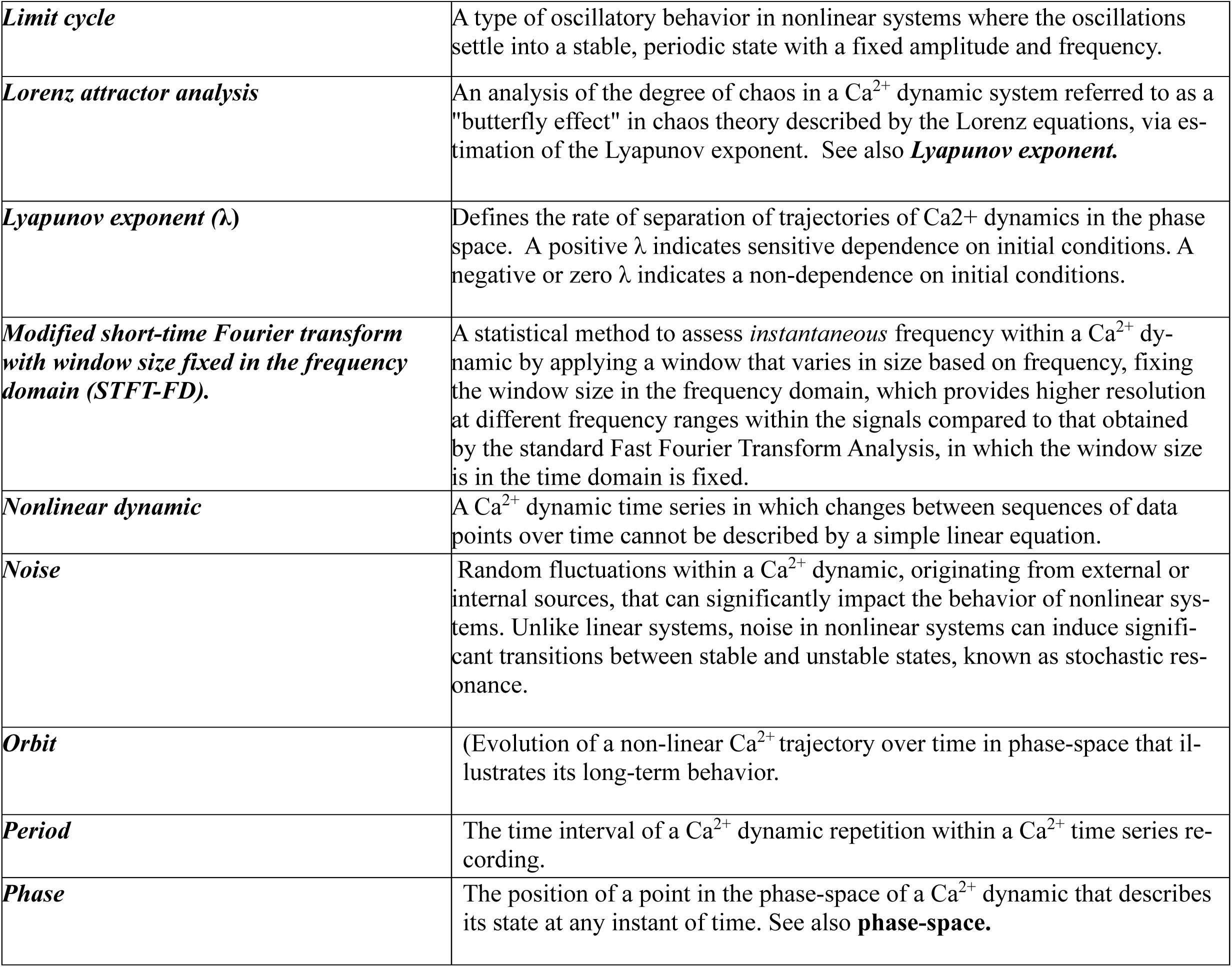
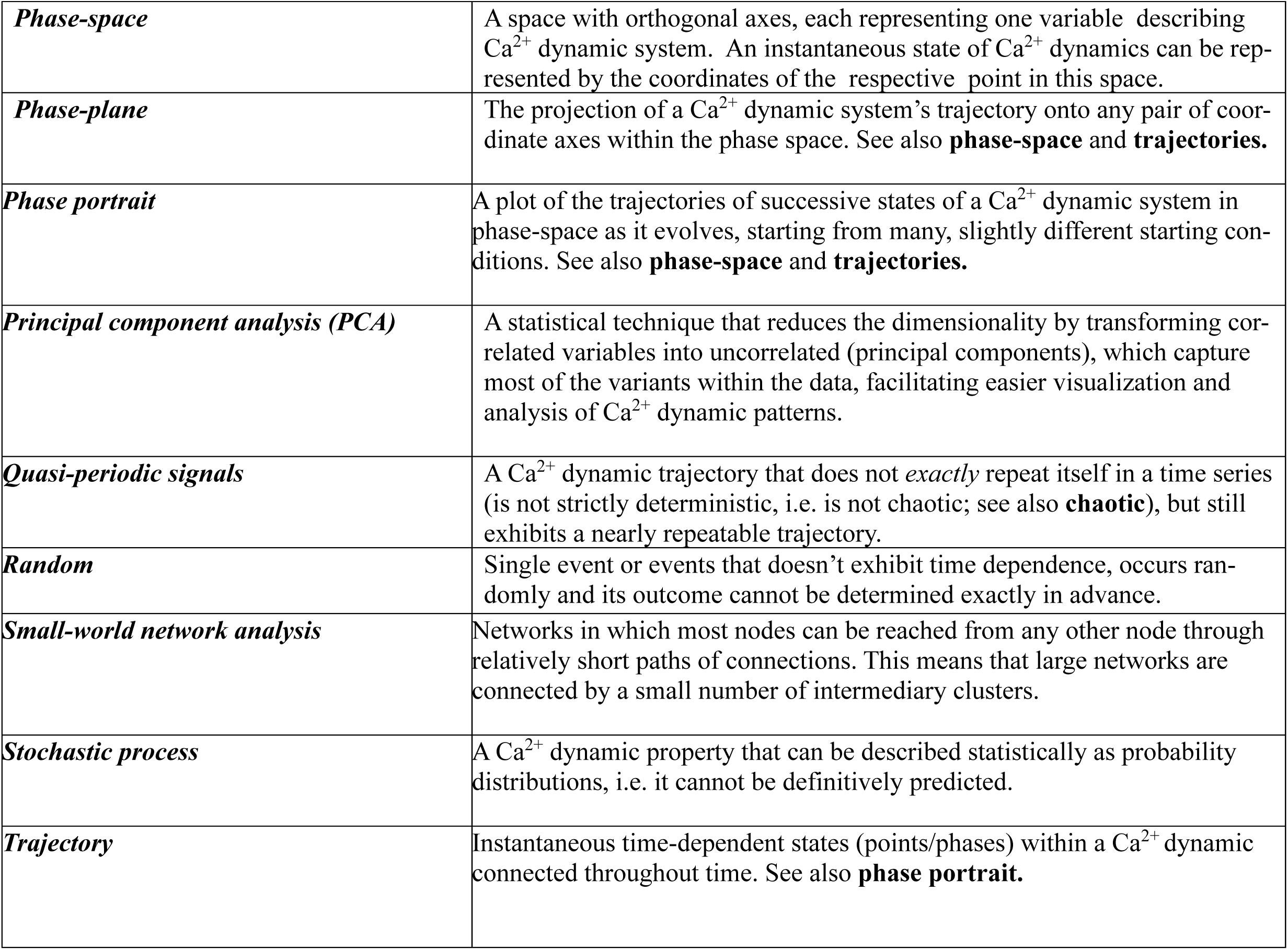

|  |  |
| --- | --- |
| *Acrophase* | The time from a fixed reference point at which a $\text{Ca}^{2+}$ signal reaches its maximum. |
| *Attractor* | A point or points in which a system is stationary. An attractor can be a single fixed point, multiple points, a loop, or a complex orbit. See also **fixed point**. |
| *$\text{Ca}^{2+}$ Dynamic* | Trajectory of how $\text{Ca}^{2+}$ changes over time. |
| *Chaotic $\text{Ca}^{2+}$ Dynamic* | Nonlinear $\text{Ca}^{2+}$ dynamic trajectories that are determined by initial conditions to irregular time series that are sensitive to the initial conditions where small changes in the initial conditions produce substantial changes in its trajectory. |
| *Cluster analysis* | A statistical method that organizes heterogenous $\text{Ca}^{2+}$ dynamics data based on certain shared characteristics (e.g. phase). |
| *Embedding dimension* | The minimum number of dimensions required to reconstruct the space state of the average $\text{Ca}^{2+}$ dynamic of a system of $\text{Ca}^{2+}$ dynamic clusters from a time series using delayed copies of the same parameter (e.g. amplitude). |
| *Empirical Mode Decomposition (EMD)* | An adaptive decomposition technique to extract frequencies embodied within $\text{Ca}^{2+}$ dynamics. EMD together with HHT specifically useful for the analysis of nonlinear and nonstationary signals. Used in the context of Hilbert-Huang Transform Analysis of $\text{Ca}^{2+}$ dynamics, see below. |
| *Functional data analysis (FDA)* | A statistical method to study patterns and trends by comparing shapes of given parameters of a $\text{Ca}^{2+}$ dynamic (e.g. velocity&acceleration) over time. |
| *Fixed point* | A specific state in which the rate of change (velocity) of $\text{Ca}^{2+}$ dynamic does not change within a $\text{Ca}^{2+}$ dynamic. |
| *Hilbert-Huang Transform (HHT) analysis* | A technique to decompose a $\text{Ca}^{2+}$ dynamic into so-called intrinsic mode functions (IMF) to determine instantaneous frequencies, to quantify how the power within the $\text{Ca}^{2+}$ dynamic is distributed across these frequencies. |
| ***Limit cycle*** | A type of oscillatory behavior in nonlinear systems where the oscillations settle into a stable, periodic state with a fixed amplitude and frequency. |
| ***Lorenz attractor analysis*** | An analysis of the degree of chaos in a $\text{Ca}^{2+}$ dynamic system referred to as a "butterfly effect" in chaos theory described by the Lorenz equations, via estimation of the Lyapunov exponent. See also ***Lyapunov exponent***. |
| ***Lyapunov exponent ($\lambda$)*** | Defines the rate of separation of trajectories of $\text{Ca}^{2+}$ dynamics in the phase space. A positive $\lambda$ indicates sensitive dependence on initial conditions. A negative or zero $\lambda$ indicates a non-dependence on initial conditions. |
| ***Modified short-time Fourier transform with window size fixed in the frequency domain (STFT-FD).*** | A statistical method to assess *instantaneous* frequency within a $\text{Ca}^{2+}$ dynamic by applying a window that varies in size based on frequency, fixing the window size in the frequency domain, which provides higher resolution at different frequency ranges within the signals compared to that obtained by the standard Fast Fourier Transform Analysis, in which the window size is in the time domain is fixed. |
| ***Nonlinear dynamic*** | A $\text{Ca}^{2+}$ dynamic time series in which changes between sequences of data points over time cannot be described by a simple linear equation. |
| ***Noise*** | Random fluctuations within a $\text{Ca}^{2+}$ dynamic, originating from external or internal sources, that can significantly impact the behavior of nonlinear systems. Unlike linear systems, noise in nonlinear systems can induce significant transitions between stable and unstable states, known as stochastic resonance. |
| ***Orbit*** | (Evolution of a non-linear $\text{Ca}^{2+}$ trajectory over time in phase-space that illustrates its long-term behavior. |
| ***Period*** | The time interval of a $\text{Ca}^{2+}$ dynamic repetition within a $\text{Ca}^{2+}$ time series recording. |
| ***Phase*** | The position of a point in the phase-space of a $\text{Ca}^{2+}$ dynamic that describes its state at any instant of time. See also **phase-space**. |
| ***Phase-space*** | A space with orthogonal axes, each representing one variable describing $\text{Ca}^{2+}$ dynamic system. An instantaneous state of $\text{Ca}^{2+}$ dynamics can be represented by the coordinates of the respective point in this space. |
| ***Phase-plane*** | The projection of a $\text{Ca}^{2+}$ dynamic system's trajectory onto any pair of coordinate axes within the phase space. See also **phase-space** and **trajectories**. |
| ***Phase portrait*** | A plot of the trajectories of successive states of a $\text{Ca}^{2+}$ dynamic system in phase-space as it evolves, starting from many, slightly different starting conditions. See also **phase-space** and **trajectories**. |
| ***Principal component analysis (PCA)*** | A statistical technique that reduces the dimensionality by transforming correlated variables into uncorrelated (principal components), which capture most of the variants within the data, facilitating easier visualization and analysis of $\text{Ca}^{2+}$ dynamic patterns. |
| ***Quasi-periodic signals*** | A $\text{Ca}^{2+}$ dynamic trajectory that does not *exactly* repeat itself in a time series (is not strictly deterministic, i.e. is not chaotic; see also **chaotic**), but still exhibits a nearly repeatable trajectory. |
| ***Random*** | Single event or events that doesn't exhibit time dependence, occurs randomly and its outcome cannot be determined exactly in advance. |
| ***Small-world network analysis*** | Networks in which most nodes can be reached from any other node through relatively short paths of connections. This means that large networks are connected by a small number of intermediary clusters. |
| ***Stochastic process*** | A $\text{Ca}^{2+}$ dynamic property that can be described statistically as probability distributions, i.e. it cannot be definitively predicted. |
| ***Trajectory*** | Instantaneous time-dependent states (points/phases) within a $\text{Ca}^{2+}$ dynamic connected throughout time. See also **phase portrait**. |

## 4. Results

Previous high-spatial resolution SAN Ca²⁺ imaging studies have demonstrated the occurrence of incessant, stochastic, and heterogeneous local Ca²⁺ oscillations (LCOs) within SAN pacemaker cells (**see Video S1**).

### 4.1. Global SAN Ca^2+^ transients (CaT) emerged during time series Ca^2+^ recordings

We performed high-speed time-series imaging of Ca²⁺ dynamics in ex vivo mouse SAN tissue (n=4) to investigate how information encoded in heterogeneous LCOs integrates into global SAN Ca²⁺ impulses (Ca²⁺ transients, CaTs). Repetitive, spontaneous, rhythmic global SAN CaTs emerged during 2.7 sec Ca^2+^ recordings (**Fig. 1A**). Mean peak to peak interval of the global CaT between different SANs (200.15±43.42-ms, n=4) was similar to the mean peak to peak AP interval (199.37±43.25 ms, n=4) (**Table S1**), recorded by a sharp intracellular microelectrode in the right atrium, indicating that the frequency of the global CaTs was the same as frequency of electrical impulses that emerged from the SAN to activate the atrium. The general sine wave-like shape of the CaTs suggested that there more than one mechanism determines each CaT.

**Fig. 1.**
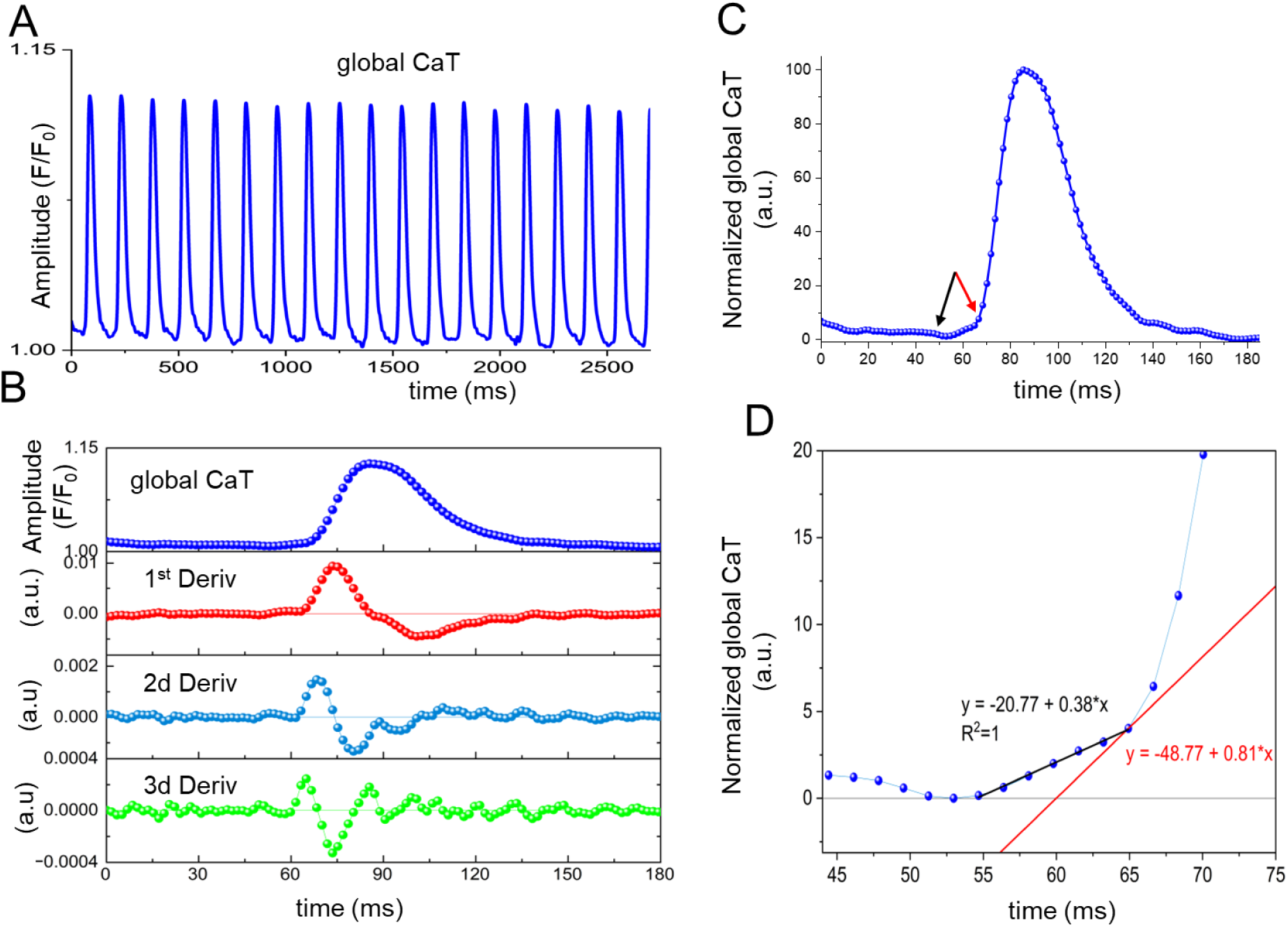
Kinetic analysis of the dynamics of the global Ca^2+^ impulses, i.e. Ca^2+^ transients (CaTs) in SAN tissue. **(A)** SAN CaTs (F/F_0_) waveforms during 2.7 sec Ca^2+^ recordings. **(B)** Kinetic analysis of the first CaT (upper Panel) during first impulse and its first (dCaT/dt), second (d2CaT/dt2), and third derivatives (d3CaT/dt3) in lower Panels. **(C)** Normalized waveform of the first CaT in Panel A. Peak 1^st^, 2^nd^ and 3^rd^ derivatives occurred at 73.5 ms and 68.3 ms and 64.9 ms, respectively, at times at which the amplitude of the CaT achieved 45%, 11.8% and 4% of its maximum amplitude, respectively. Arrows show onset (**black**) and end (**red**) of the foot (ignition). **(D)** Zoomed **Panel C** between ∼45-75 ms of the first impulse. The slow phase of normalized CaT in **Panel B** (occurring between ∼50-65 ms during impulse formation) was modeled using a tangent (**red line**) and a linear fit (**black line).** The earliest inflection point occurred at 64.9 ms, coinciding with the peak of the 3^rd^ derivative. At the inflection point time, the tangent slope was 0.81, and the integrated global Ca^2+^ signal amplitude had achieved 4.02% of its maximum. A linear model fitted to the Ca^2+^ signal during the interval from 50-65 ms showed strong local linearity (approximately 12 ms) with a slope of 0.38 (a.u./ms, r = 1.0). The peak-to-peak intervals of the global CaT during the 2.7second time-series in Fig. 1A are shown in **Fig. S1, Panel A**.

Figure 1B illustrates a detailed kinetic analysis of the first integrated (global) SAN CaT, the Ca^2+^ signal of which began to *steadily increase* from baseline levels (background noise) with onset of the Ca^2+^ signal 3^rd^ derivative at ∼45-50 ms (**Panel B**), and continued to increase slowly until the time at which 3^rd^ derivative of CaT reached its peak (at 64.9 ms), marked by an inflection point in the global CaT trace in Panel C and by a tangent line in **Panel D**. Beyond this inflection point, the global CaT rapidly accelerated, reaching its peak about 20 ms later. Thus, in SAN tissue, the global CaT exhibits a clear, slow foot that lasts about 15ms and contributes approximately 4.02% to the CaT peak amplitude.

### 4.2. Global SAN CaTs are formed by integration of variable Ca^2+^ dynamics within regional functional clusters of pacemaker cells

Next, we related temporal characteristics of the global CaT in **Fig. 1** to Ca^2+^ information flow during and between the formation of SAN global CaTs within the time-series recordings, by extracting characteristics of the local Ca^2+^ dynamics (LCO), such as amplitude and acrophase, every 1.7 ms in 50 regions of interest (ROIs) that encompassed the entire visualized area in **Figure 2A, Video 1**. LCO amplitudes (F/F_0_) differed within and among ROIs (**Figure 2B**). **Video 2** illustrates the 3D relationships between LCO amplitudes (normalized) and the time within each of the ROIs in each cluster at every 1.7 ms during the initial 180 ms of the time-series recording.

**Fig. 2.**
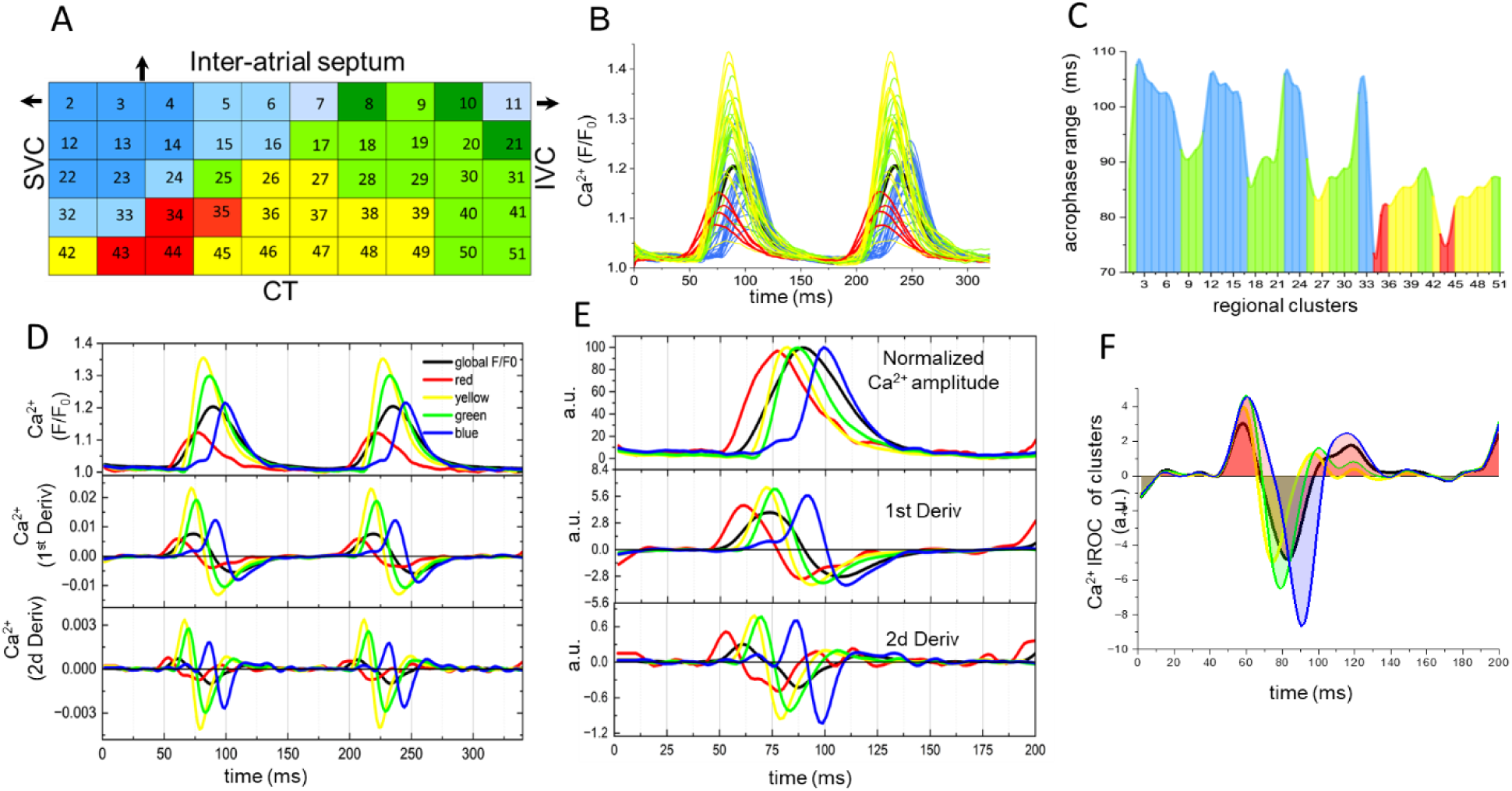
Analyses of the dynamics of local Ca^2+^ oscillations and global Ca^2+^ transients (CaT) in SAN tissue. **(A)** Virtual map of the spatial SAN area from which 50 regions of interest (ROIs) LCO characteristics were extracted from every 1.7 ms . **(B)** Variability of LCO amplitudes (F/F_0_) of 50 ROIs during time series recordings. Superimposed black trace represent a global CaT**. (C)** Acrophases (the times when the maximum amplitudes of the Ca^2+^ dynamics within each ROI were achieved during impulses) of *individual* ROI LCOs mapped on color coded SAN virtual space**. See also <u>Video 1</u> and Video 2. (D)** Average LCO amplitudes of four-color coded clusters (upper Panel) during ∼320 ms time series recordings and its first (dCaT/dt) and second (d2CaT/dt2) derivatives (lower Panels). **(E)** Normalized LCO waveform during and its derivatives during the first global CaT (overlaid blac trace). **(F)** The instantaneous rate of change (IROC) between clusters and the global CaT presented (black trace). The relative IROC of the red color-coded clusters to that of other clusters and the global CaT, determined by subtracting the 1^st^ derivatives of the yellow, green, and blue clusters and global CaT, from the 1^st^ derivative of the red cluster at each time point. Note that the increase in the IROC for the red color-coded cluster during the initiation of the impulse (pre-foot and foot stages, initial ∼15-65 milliseconds of impulse formation) occurred earlier and exhibited a greater magnitude than that of the other clusters, which demonstrated minimal variation during this timeframe.

Acrophases (the times when the maximum amplitudes of the Ca^2+^ dynamics within each ROI were achieved during impulses) of *individual* ROI LCOs (**Fig. 2C**), as color-coded in **Fig.2A**, differed by up to ∼ 34 ms. Note that distributions of acrophases within the SAN ROIs formed distinct spatial clusters having similar, yet not identical phases (**Fig. 2C**).

We next calculated the average Ca^2+^ dynamics within each color-coded cluster of ROIs visualized in **Fig. 2A**. The shortest average Ca^2+^ dynamic acrophases (76.5 ± 2.9 ms) were recorded within the red functional cluster, whereas the longest acrophases (103.1 ± 2.9 ms) occurred in the blue cluster. The mean acrophases of the yellow and green clusters were 83.9 ± 1.2 ms and 89.2 ± 1.9 ms, respectively (**Panel D**).

Figure 2D shows an example of the global CaT and its derivatives (a black trace in both the upper and lower Panels) overlaid on the average LCO Ca^2+^ dynamics from the four distinct color-coded clusters, including their respective derivatives.

Cross-talk between local Ca^2+^ dynamics within the network of clusters formed rhythmic SAN global Ca^2+^ transients (CaTs), during which the phase and kinetic uniqueness of the Ca^2+^ dynamic within each cluster was preserved. The *mean* peak-to-peak Ca^2+^ dynamic interval and mean standard deviations (SDs) across the time-series recording (**Fig. S1B**) in the red cluster differed from that of the other clusters, with the red cluster showing the highest SD (∼30%).

The ensemble Ca^2+^ signal of the red cluster was the first to increase not only in this impulse but across all 19 impulses that emerged within the time-series. The time at which the Ca^2+^ signal of the red cluster began to accelerate (as indicated by the initiation of its second derivatives in the lower Panel) at ∼45 ms following the onset of the recording and corresponding approximately to the time of initiation of the foot of the global CaT **(Fig. 1C-D)**. The maximum velocity of this signal occurs at about 61 ms, a time that was close to the occurrence of the foot of the global CaT, as depicted in **Fig. 1**. Although the contribution of the red cluster dominated the foot of the global SAN CaT, it is important to note that the Ca^2+^ dynamic in other clusters also began to increase during the foot, but at a later time than the Ca^2+^ dynamic of the red cluster **(Panels D-F).** Even *prior* to the initiation of the foot, the amplitude of the integrated Ca^2+^ signal of the red cluster dominated that of all other clusters (on average up to 3.6 fold, **Fig. S2**). Therefore, the Ca^2+^ dynamics within each of four functional clusters contributed to the pre-foot and foot formation of the global CaT, although starting at different times. This diversity in timing of regional Ca^2+^ dynamics is crucial to the shape formation, and to the overall development of the foot of the global CaT. The maximum velocity of the normalized amplitude of the Ca^2+^ signal of the red (4.6 a.u. at 61ms) cluster was slower compared to that of the yellow (6.6 a.u. at 72ms), green (6.5 a.u. at 77ms). The blue cluster differed from other clusters in having a biphasic rate of rise: the velocity of the initial slow component occurring at 73ms and was 0.9 a.u. followed by rapid increase in velocity to 5.7 a.u. which peaked at 91ms **(Panel E)**.

The growth of the instantaneous rate of change (IROC) (**Panel F**) in the red color-coded cluster occurred earlier and was greater compared to the three other clusters and the global CaT during the impulse initiation phase (from approximately 15 to 61 ms, covering the pre-foot and foot stages). During this period, the other clusters either exhibited minimal changes or remained unchanged (see **Panel F**, highlighted red areas above zero). At the time of maximum velocity in the red cluster, the Ca^2+^ dynamic and its IROC in the yellow cluster began to rise rapidly, closely followed by a similar increase in the Ca^2+^ dynamic in the green cluster. Simultaneously, there was a gradual increase in the Ca^2+^ dynamics within the blue clustered ROIs (between ∼61-71 ms of impulse formation) followed by a plateau of about 7 milliseconds, following which the blue cluster accelerated rapidly. The average peak amplitude of the Ca^2+^ signal in the blue color-coded cluster was intermediate between those of the red, yellow, and green color-coded clusters (**Panel D**).

### 4.3. Frequency components analysis of the SAN Ca^2+^ dynamic

Frequency domain analysis indicated that a wide range of physiological frequencies (2-12Hz) underpinned each global Ca^2+^ impulse (**Fig. S3**). Because Ca^2+^ dynamic characteristics within SAN regional clusters are ultimately derived from the frequency characteristics of biologic noise [29, 30] underlying LCOs, the frequency components of the total power (energy) expended in each cluster must differ. We quantified frequency domain characteristics of Ca^2+^ dynamics in each color-coded cluster throughout the entire 2.7-sec time-series recording (**Fig**. **3A**).

**Fig. 3.**
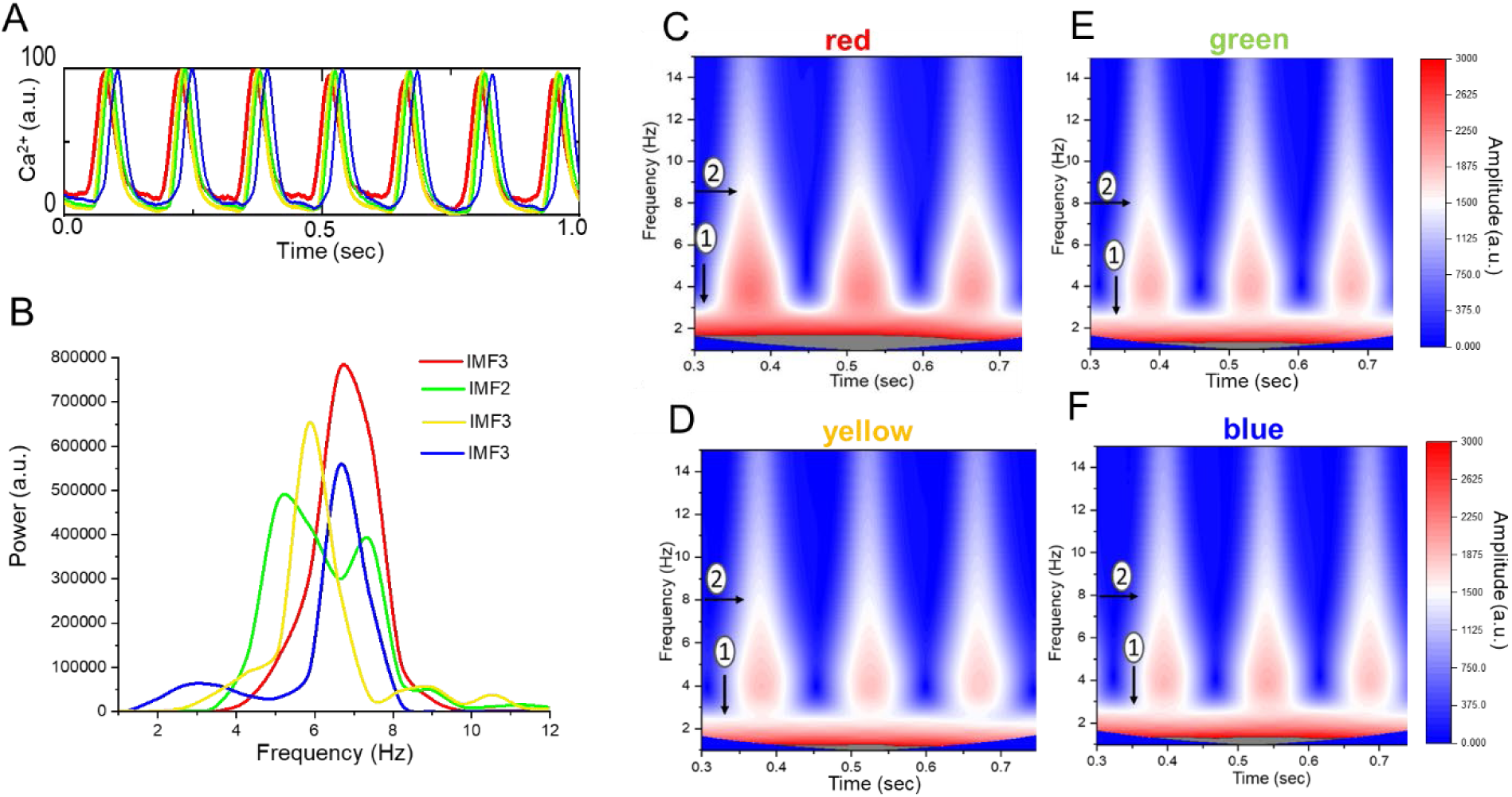
Frequency components of total energy expenditure throughout SAN tissue. **(A)** Upper Panel, illustrates the superimposed normalized (on a scale from 0 to 100) Ca^2+^ dynamic waveforms of the average Ca^2+^ dynamics of the four distinct, color-coded clusters during the entire 2.7 sec. time series recording; lower Panels present the decomposition of the Ca^2+^ dynamics in Panel A into its intrinsic mode functions (IMF) by empirical mode decomposition (EMD) and Hilbert-Huang Transforms (HHT) from which *instantaneous* frequency ranges were extracted[31] (. Complete frequency ranges for all IMFs are detailed in **Fig. S4**. **(B)** The instantaneous frequencies corresponding to the predominant power derived from IMFs of the four color-coded clusters (Fig. S4). **(C-F**) Short-Time Fourier Transform (STFT-FD (with window size fixed in the frequency domain) of four regional color-coded signals. **t**ime-frequency-power representation of the red, yellow, green, and blue color-coded cluster respectively between 0.3-0.75 sec of the 2.7 sec. time series. The color-coded bar graph represents the frequency power (Amplitude). The low-frequency power zone is designated by oval 1 and the high-frequency power zone by oval 2.

**Panel B** shows the major frequency composition of each color-coded cluster. Note that a large range of frequencies from low (2Hz) to high (15 Hz) are embedded within the Ca^2+^ dynamic of four color-coded clusters (**Panel B**). The dominant frequency of highest power (energy) within the Ca^2+^ dynamic of each color-coded cluster emerged at 6.6Hz in the red cluster, 5.9Hz in the yellow, and about 5.1Hz in the green cluster. Although the dominant frequency of the blue cluster was similar to that of the red cluster (around 6.6Hz, **Fig. 3B**), the blue cluster exhibited much lower energy (power) at this frequency. The entire range frequencies of all of the clusters is shown in **Fig. S4**.

In order to determine frequency-*time*-power relationships of the Ca^2+^ dynamic of each of the four color-coded regions, we converted regional signals in **Panel A** to the time-frequency and amplitude domains. **Panels C-F** show the converted regional Ca^2+^ signals during first three impulses in Panel A. A range of frequencies from low (indicated by oval 1) to high (indicated by oval 2), in red, yellow, green, and blue clusters, respectively. Note, however, that in the red cluster (**Panel C**) high and low frequency components and the frequency power (see the bar code) were larger, and the abrupt transition from lower to higher frequencies in this cluster began earlier during the foot of each CaT (**Panel C**) compared to other clusters (**Panels D-F**). It is clear however, that the frequency ranges of the four clusters overlap over a range of time points during each impulse.

To further clarify this relationship, we performed principal component analysis (PCA) every 17.3 ms (covering 10 data acquisition points) across the initial 190 ms of the Ca^2+^ time-series recording. An example of the PCA analysis during this first impulse is illustrated in **Fig. S5**. The first two principal components, PC1 and PC2, accounted for over 90% of the total variance in the Ca^2+^ dynamics within the entire first global CaT. During the initial 0-36 ms of the recording **(Fig. S5, Panels B and C),** and *prior to* the initiation of the foot of the global CaT, PC1 and PC2 accounted for approximately 96% and 61%, respectively, of the variance. As the foot of the global CaT began to form (between ∼45-65 ms), the contribution of the red cluster increased to 99% (**Fig. S5D**). Beyond the foot, during the rapid upstroke, and up to the peak of the global CaT, the yellow and green clusters made the greatest contribution. The blue cluster, followed by the green cluster made the greatest contribution during relaxation of the global CaT. See supplement for further information.

Supplementary **Fig.S6** shows LCO dynamics (1^st^ derivative) of the entire 50 regions of interest (ROIs), grouped into four color-coded clusters (**see Fig. S6**) throughout the entire time series (2. 7 sec) Ca²⁺ recording. Note that the ellipse representing the yellow cluster in **Fig S6**, overlaps substantially with that of the green cluster, and the green cluster overlaps with the blue, while the red ellipse connects all three clusters. Thus, the Ca²⁺ dynamics across *all* SAN functional clusters *collectively* influence the dynamic of the global SAN CaT (as illustrated in **Figs. S5 and S6**).

### 4.4. Stability of Ca^2+^ Dynamics within SAN Spatial Domains

The different times at which the Ca^2+^ dynamics of each cluster were initiated hinted that the stability of Ca^2+^ dynamics differed among clusters. We analyzed the stability behavior within the network of SAN functional color-coded clusters during impulse initiation (as described in the methods). **Figure 4** illustrates the normalized amplitudes of SAN Ca^2+^ impulses (**Panel A**) and 1st derivatives (velocities, **Panel B**) of four color-coded SAN clusters superimposed on SAN global CaT during the first 300 ms of time-series recordings (during which two global CaT emerged). **Panels B and C** show that the Ca^2+^ dynamic within each color-coded spatial domain consistently transitioned between stable (closed circles) and unstable states (open circles). Following a prior impulse, the Ca^2+^ dynamic of all four clusters reached the stable fixed points (within approximately 40 ms), before the Ca^2+^ dynamic of the red cluster diverged (after ∼7 ms) toward an unstable point, as its velocity steadily increased during impulse formation ∼45-63 milliseconds of the recording.

**Fig. 4.**
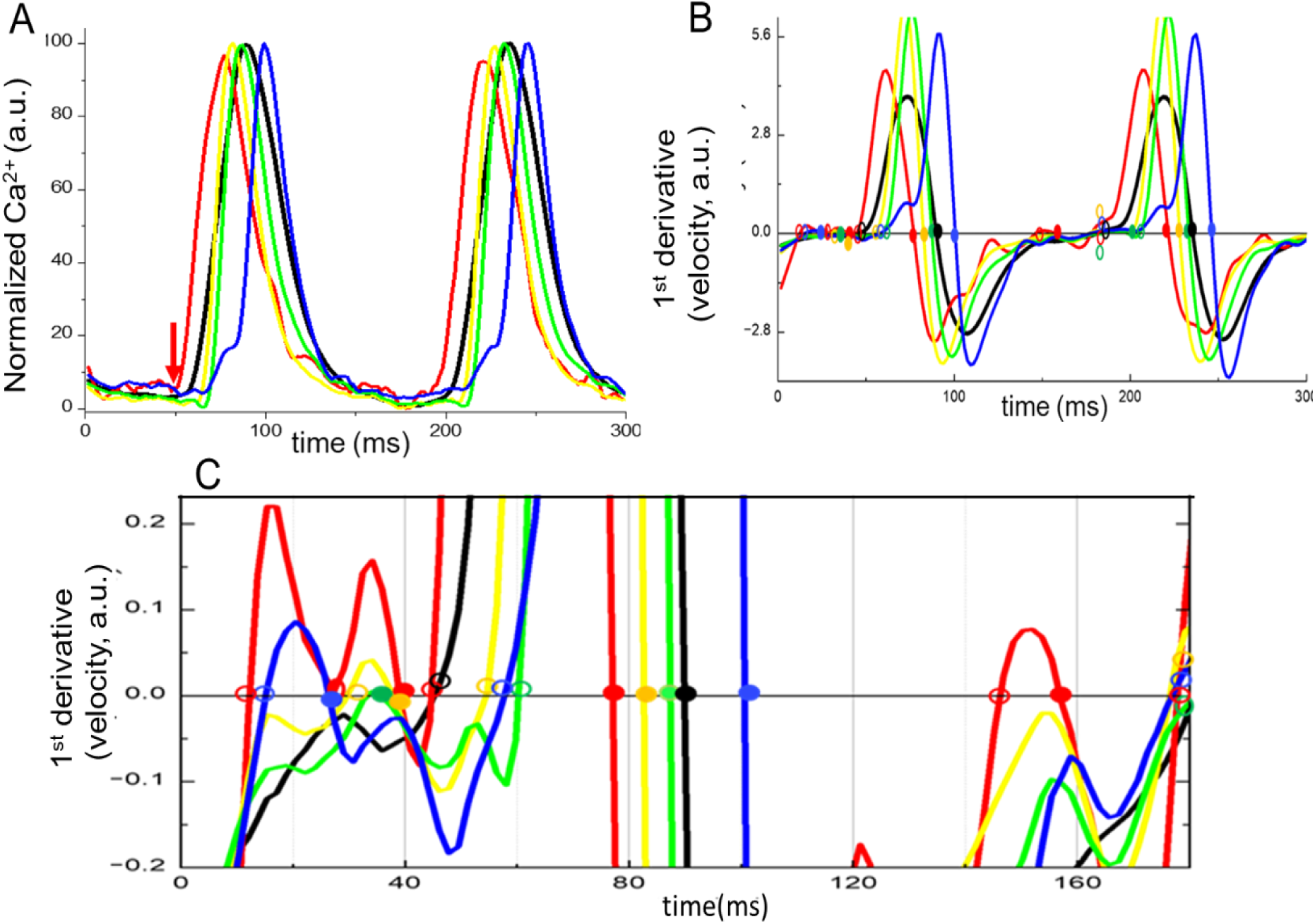
Stability of LCOs within SAN spatial domains. **(A)** The normalized amplitude of the SAN Ca^2+^ dynamics across four distinct color-coded clusters during a time period of 300 ms (encompassing two impulse formations during a time series). **(B)** Velocity change, represented as the 1^st^ derivative of the Ca^2+^ dynamics of the clusters in Panel A. The circles indicate fixed points at which time the velocity is zero; closed circles denote stable fixed points, while open circles represent unstable fixed points. **(C)** Fixed and unstable points of Ca^2+^ dynamics the four color-coded clusters during the initial cycle in Panel A illustrated at increased resolution. If a fixed point is stable, the velocity (dCa^2+^/dt) of the points in its vicinity will be *negative* (dCa^2+^/dt < 0), i.e. the fixed point is considered to be a ‘sink’ and the signal is moving *toward* equilibrium. If the velocity of points near the fixed point is positive (dCa^2+^/dt > 0), the fixed point is unstable, or a ‘source’, from which the signal diverges exponentially from equilibrium.

The timing of this unstable fixed point in the red cluster aligned with the onset point of the foot of the 1st global CaT, which occurred between 45-50 ms (arrow in Panel A, see also global CaT foot in **Figs. 1**). The yellow, green, and blue curves transitioned toward the unstable fixed points at ∼55, 60, and 58 ms, respectively. Following the transition, the four SAN Ca^2+^ spatial domains (red, yellow, green, and blue) all reached the next ‘quasi-stable fixed point’(at the time of a maximum of 1^st^ derivative, and not discussed in this study), returning to the stable state at ∼77, 83, 87 ms red, yellow and green clusters, respectively, but much later in the blue cluster at 101 ms (**Panel C** closed circles).

Note, that the times to achieve equilibrium for the yellow and green clusters approximately coincided with the time to achieve the peak of the global Ca^2+^ signal (∼85ms). In contrast, the red cluster reached its fixed point before and the blue cluster after the peak of the global Ca^2+^ signal had already occurred (**Panel A**, black curve). However, Ca^2+^ dynamics in all spatial domains eventually regained stability with their Ca^2+^ peak values occurring at different times. Importantly, following the completion of the first global CaT, and before the onset of the next CaT, oscillations in the stability of the Ca^2+^ dynamic *only* observed in the red cluster, following which the Ca^2+^ dynamic of the red cluster, eventually moved to the vicinity of an unstable fixed point (∼180ms) to initiate the foot of the global CaT in the next cycle. The Ca^2+^ dynamic of the three other clusters reached their unstable points recapitulating the events during the initiation of the prior global CaT.

### 4.5. Hidden non-linear structures within the SAN Ca^2+^ dynamics

Although the aforementioned linear analyses (**Figs.1-4**) were instructive with respect to understanding how heterogenous Ca^2+^ signals form global impulses, these analyses do not completely capture complex, non-linear interactions within the SAN Ca^2+^ dynamic. We performed a number of detailed non-linear analyses that detected hidden structures within each cluster differed from each other, including rotor-like energy transfer, and degrees of stochasticity and recurrence within a given cluster over time.

#### 4.5.1. Ca^2+^ Dynamic Energy Transitions Differ Among within SAN Functional Clusters

We performed functional data analysis (FDA)[32] the dynamic of energy transitions in each colorcoded cluster throughout the time-series. Figure 5B illustrate the averaged amplitudes and derivatives of the Ca^2+^ dynamic in each of the four color-coded clusters in Panel A during a 500 ms segment of the 2.7-sec time-series of Ca^2+^ recording, and their acceleration vs velocity diagrams are shown in Panel C. Functional heterogeneity in energy transfer could be clearly observed in the FDA trajectories of the Ca^2+^ dynamics.

**Fig. 5.**
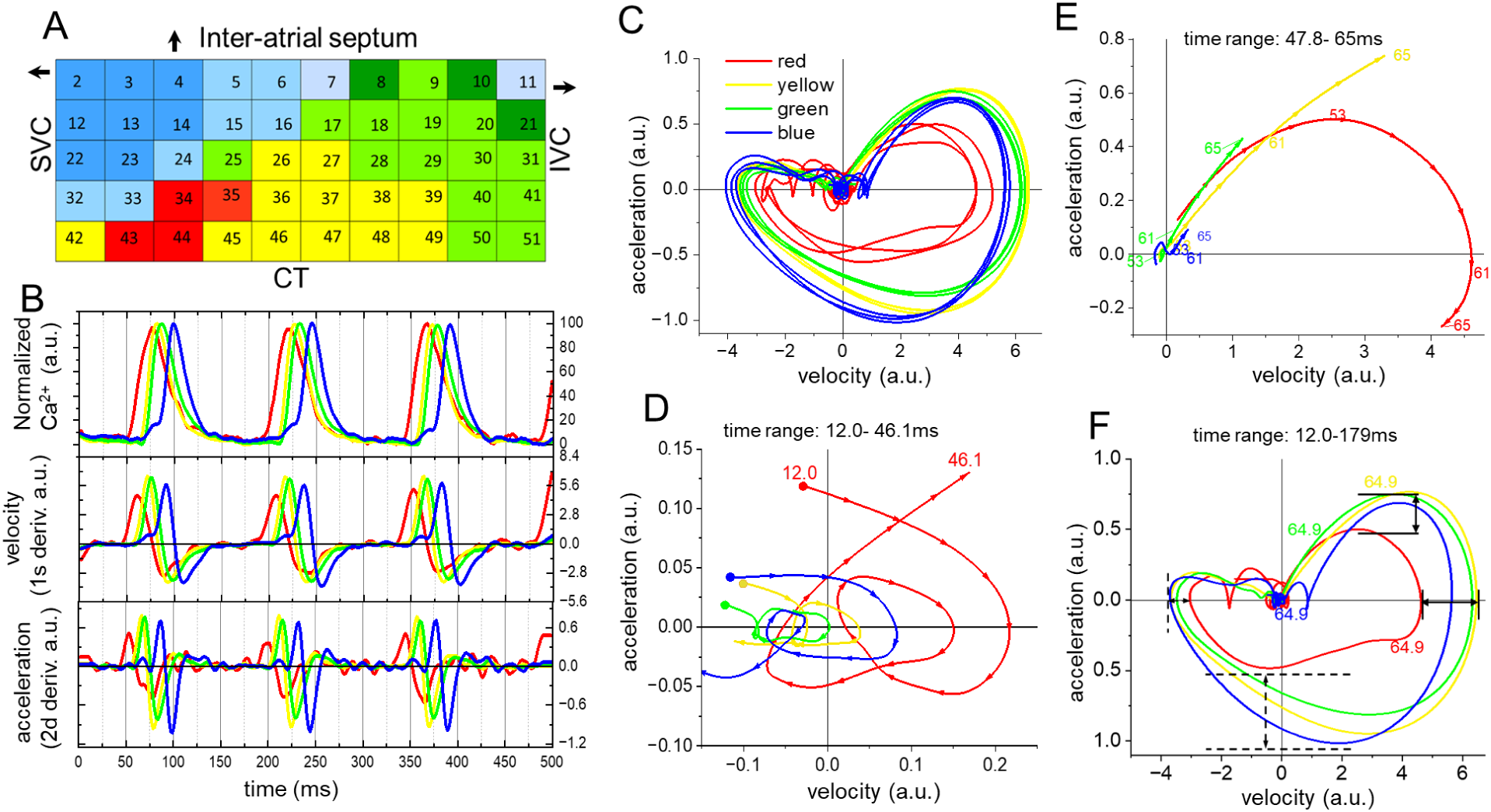
Functional data analysis (FDA) of energy transfer within Ca^2+^ dynamics of the four color-coded clusters deduced from the time-dependent kinetic changes in their Ca^2+^ dynamics. **(A)** A color-coded map illustrating the spatial distributions of SAN functional clusters. **(B)** Averaged normalized Ca^2+^ dynamics (upper Panel) of the four distinct color-coded areas in Panel A, along with their 1^st^ (middle Panel) and 2^nd^ (lower Panel) derivatives, in a 500 ms time window of a time series of Ca^2+^ recoding during which three CaTs emerged. **(C)** Phase-plane diagrams of acceleration (2^nd^ derivative) and velocity (1^st^ derivative) of the CaTs in Panel B during 500 ms of a Ca^2+^ time-series recording (during which 3 impulses in each color-coded cluster). For the simplicity **Panels D-F** show only the 1^st^ impulse formation for each of four color-coded clusters, presented at higher magnification: in **(D),** the low energy state during the pre-foot of the global CaT (between ∼12 - 46 ms); in **(E)** during formation of the foot of the global SAN CatT (∼46.1-65 ms); and in **(F)** the entire trajectories of the four clusters during the formation of the first global CaT in **Panel B**, depicting the ranges of peak maximum and minimum acceleration and velocity (solid and dashed black lines and arrows). See also **Video 3**. Note, that when *both* velocity and acceleration are *zero*, the system is in true equilibrium, and no energy is being transferred. When *either* acceleration or velocity are *non-zero*, the system is in motion, and energy is being transferred from potential to kinetic energy. When *either* acceleration or velocity is *negative*, kinetic energy is being transferred back to potential energy and vice versa. If the potential and kinetic energies within the FDA function form a perfect circle (when acceleration and velocity simultaneously cross at 0:0) a system have achieved a true equilibrium, note that this is never the case for the SAN cluster dynamics.

Panel D shows a zoom-in of the low energy state during the first impulse formation in Panel C (0-46ms, pre-foot), that includes events near zero crossing. Panel E is a zoom-in during the foot formation covering time intervals from approximately 46.1 ms to 65 ms during the first impulse formation. Panel F shows the completion of subsequent trajectory beyond 65-ms of the first impulse in Panel C. Note that the completed phase-plane plots of the Ca^2+^ dynamics (**Panels C, F**) form circular-like trajectories of *unequal* size, *neither* of which forms a perfect circle. The absence of *0:0* crossings indicates that the SAN Ca^2+^ dynamics never achieve a true equilibrium. Phase-plane analysis (**Fig. 5C-F**) indicates that the kinetic transitions of Ca^2+^ signals within regional Ca^2+^ clusters near the termination of one impulse and initiation of a subsequent impulse hovered around 0 velocities, meaning that little or no energy is being transferred at that time, (as the system approaches equilibrium at the zero crossing).

A rotor-like oscillatory behavior (‘knot’) around the zero crossing (**Panel D)** is followed by trajectories that project throughout the entire velocity-phase plane. Panel F clearly shows differences in the kinetic variation among four color-coded functional clusters (**see black arrows in Panel F**) during impulse formation, particularly between red and the other three clusters. It is important to emphasize that the red cluster’s Ca²⁺ dynamics were captured only within the specific region observed in this video, which represents 8% (4 ROIs) of the selected SAN area. The yellow cluster occupied 24% (12 ROIs), while the green and blue clusters each accounted for 34% (17 ROIs each) of the observed area (**Panel A**). Although the spatial extent of the red cluster was the smallest, and its Ca²⁺ dynamics near the CT had the lowest potential and kinetic energy, the acceleration-velocity trajectory of this cluster showed an early increase (entering the upper right quadrant in **Panels C-F**), preceding significant kinetic changes in the other clusters. The timing of the energy transition observed in the red cluster aligns with the initiation of the global calcium transient (CaT) (**Fig. 1**), and is likely implicated in the ignition of the global SAN CaT via synchronization of LCOs within this cluster. This is followed by a rapid acceleration of Ca^2+^ in other functional clusters, impacting a rapid increase in acceleration and velocity of the global Ca^2+^ impulse. The kinetic trajectory of the blue cluster (near the inter-atrial septum, **Fig. 5A**) emerged following a delay after the trajectory initiation of the other clusters, although the magnitude of its energy transition was comparable to that of the yellow and green areas (**see Fig. 5C-F**).

Supplementary Fig. 7 compares energy transitions during the 1^st^ global CaT in the time-series (**Fig. S7A-C**) and the ROI correlations during time segments of the same impulse (Fig. **S7E-G)**; and **Fig. S8** shows energy transitions for all 19 CaTs. Phase-plane diagrams of acceleration vs velocity of the integrated (global) CaT followed similar trajectories as those shown in **Fig. 5**.

**Supplementary Method Fig. 1:**
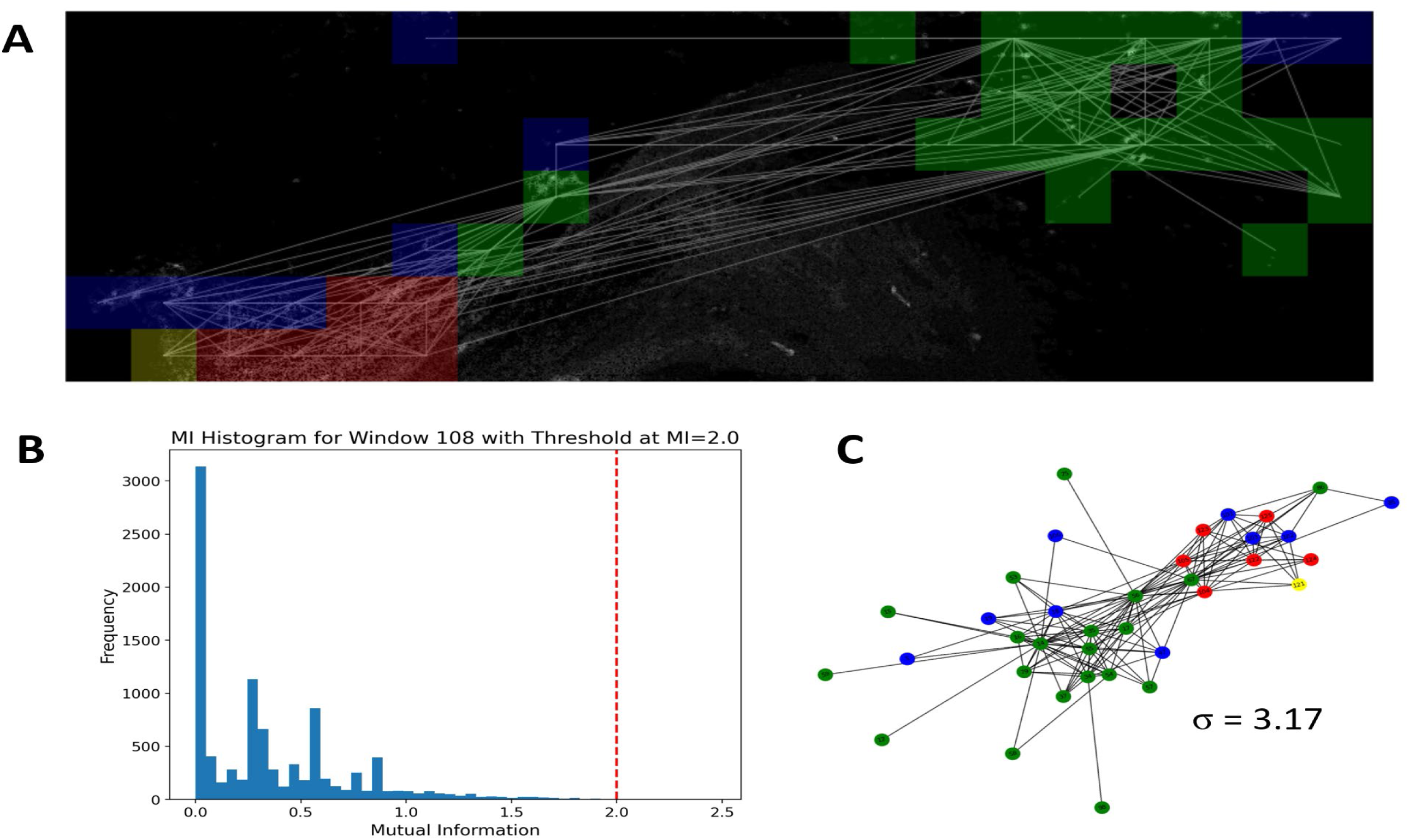
Small-world network analysis of the integrated (global) SAN Ca2+ signal. (A) Spatial representation of functional connectivity in the SAN, with colored regions indicating different functional clusters and white lines representing connections above the mutual information threshold. (B) Histogram of mutual information (MI) values for a representative time window, with the red dashed line indicating the threshold (MI=2.0) used for network construction. (C) An example of a small-world network graph for representative time window, derived from thresholded MI values, with nodes colored according to their functional cluster and σ indicating the small-worldness metric for this particular network configuration.

#### 4.5.2. Recurrence Characteristics of the SAN Ca^2+^ Dynamic System

We next performed Recurrence Quantification Analysis (RQA) which detects hidden patterns and recurrences in a time series without assuming signal stationarity, making it particularly useful for short time series and discontinuous data.

The **Fig. 6** illustrates the necessary components for generating recurrence plots (RPs) of Ca²⁺ dynamics (shown for the red cluster, **Panels E and F**) based on mutual information (**Panel B**), the false nearest neighbor method (**Panel C**), and time-delay state-space reconstructions (**Panel D**).

**Fig. 6.**
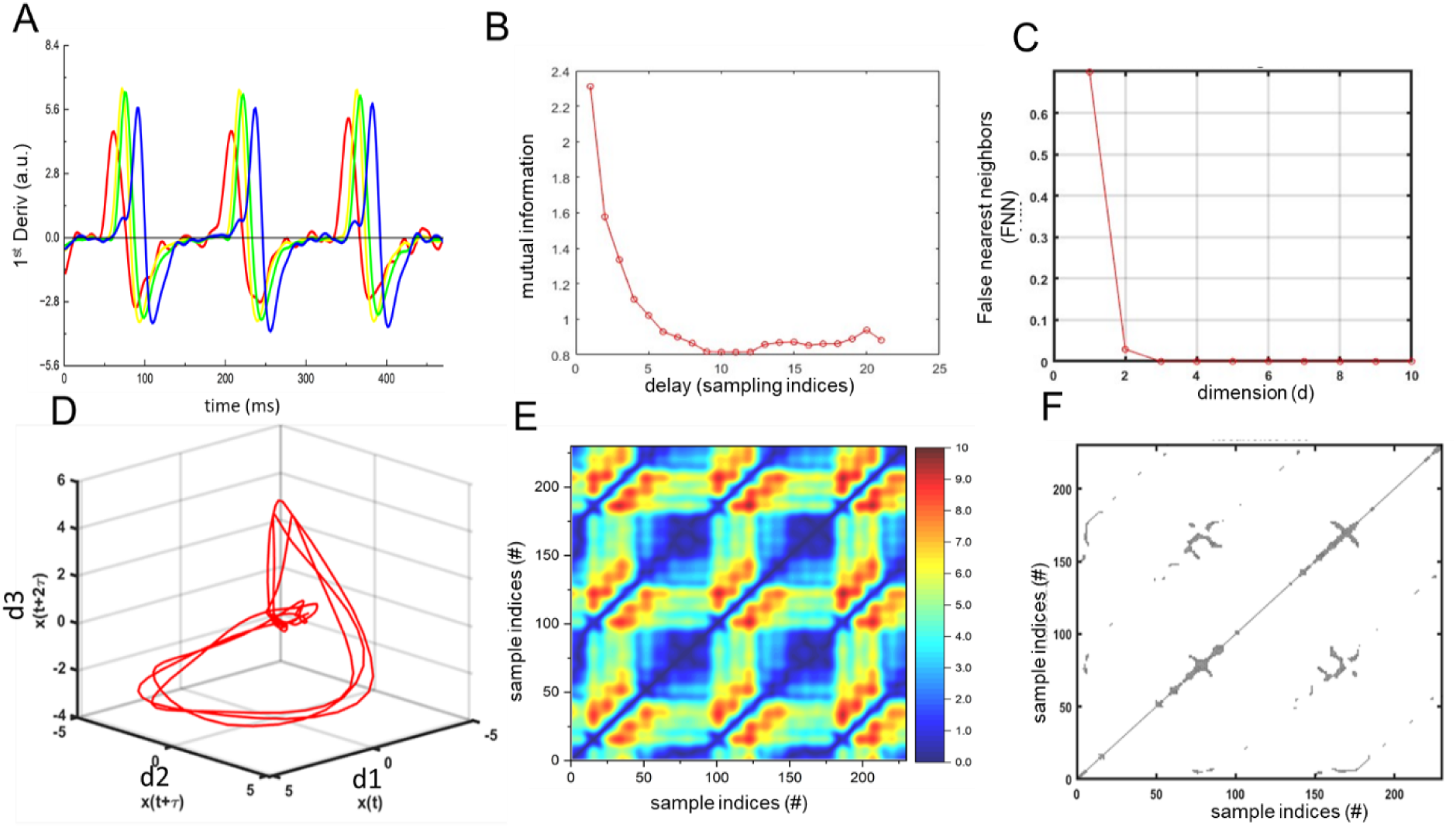
Recurrence Quantification Analysis (RQA) of the Ca^2+^ dynamics. **(A**) Superimposed Ca^2+^ dynamics (1^st^ derivative) of the red, yellow, green and blue clusters during three consecutive global SAN CaTs. The optimal time delay (10 sampling points, ∼17.03 ms) and optimal embedding dimensions of three (d = 3) used to create the state-space plots were estimated from mutual information (**B)** and false nearest neighbors (FNN) (**C)** for the red functional cluster. The time-delay embedding of the state-space **(D)** was used to recontract *unthresholded **(E)*** and *thresholded* **(F)** recurrence plots (RP) of the red cluster. MATLAB toolbox[33, 34] was used to perform this type of analysis.

The state-space reconstructions of the four color-coded clusters were then transformed into the unthresholded RP (**Panel E**, shown for the red cluster), to which the threshold was applied to construct the thresholded RPs (TRPs) (**Panel F**, shown for the red cluster, see details in the figure legend), which quantitively estimate the recurrence parameters[33, 34].

**Figure 7** displays state space plots (**upper Panels**) and their corresponding recurrence plots (**lower Panels**) for the red, yellow, green, and blue clusters respectively. Visual inspection of the thresholded RP (see methods) of the red cluster showed fewer recurrence points than those of the other clusters, indicating that the red cluster harbors more *stochastic* behavior than other clusters. The basin of attraction (attractor) of the phase space plots of the red cluster, arrow, in Fig. 7A-G also showed a higher variability compared than that of the three other clusters, that was in line with higher instability of initial conditions (fixed points **Fig. 4C**) in this cluster. The recurrence structures of the red cluster’s thresholded RPs (TRPs) were less periodic and more scattered than those of the other clusters, suggesting higher stochasticity (Fig. 7B).

**Fig. 7.**
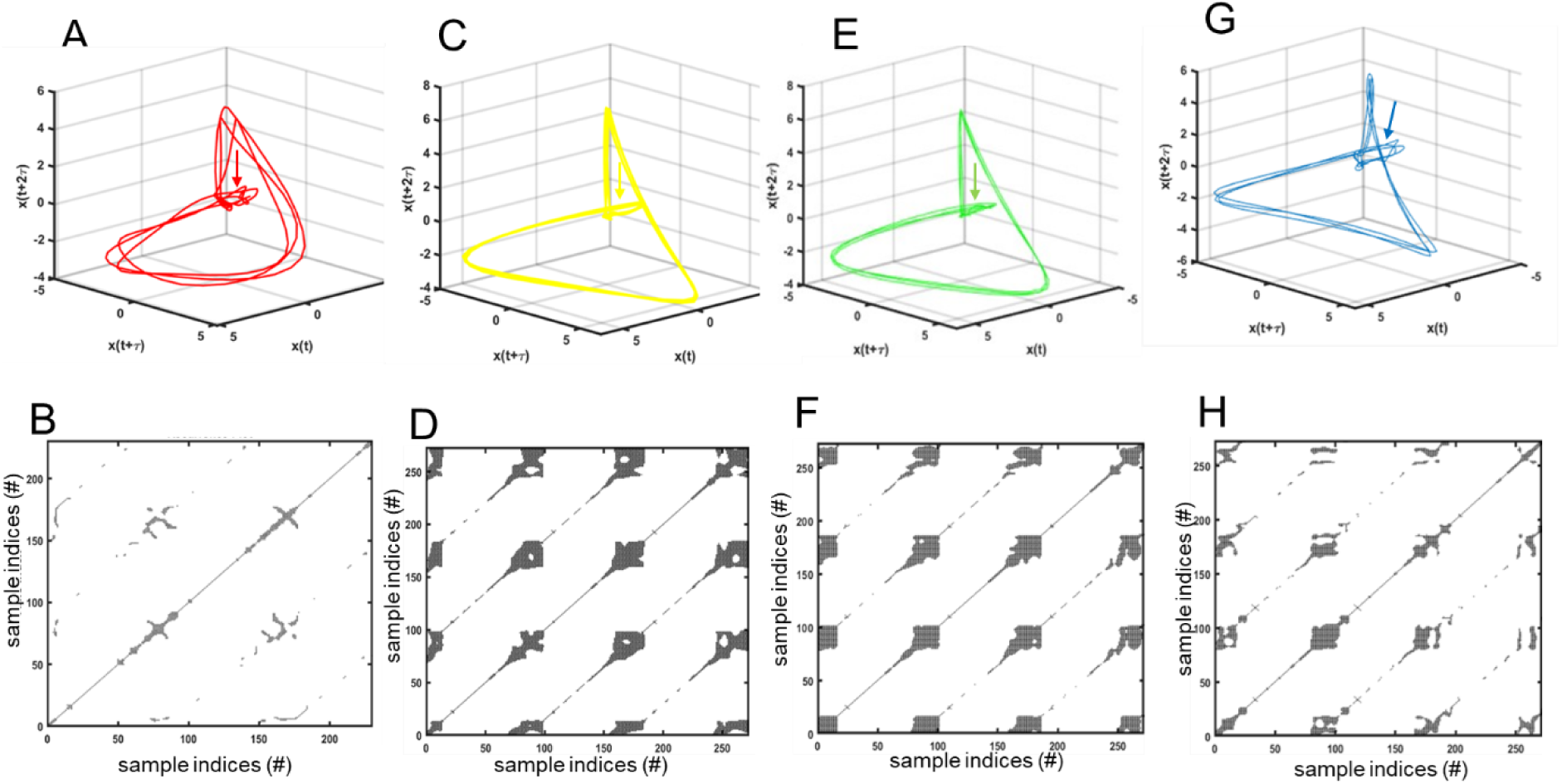
Time-delayed state-space reconstruction and recurrence plots (RP) of the Ca^2+^ dynamic of SAN regional clusters. **(A&B)** State-space plot and thresholded recurrence plot of the Ca^2+^ dynamic of the red cluster (also shown in Figure 6D and 6F respectively), (**C & D**) of the yellow cluster, (**E&F**) of the green cluster, and (**G&H**) of the blue cluster. For all four clusters, a time lag of 10 sampling points (t =17.03 ms) and an embedding dimension of three (d = 3) for state-space reconstruction were employed. See Fig.6 legend and methods for a detailed description of analysis.

The recurrence structures of the red clusters RP were less periodic and more scattered than those of the other clusters, suggesting higher stochasticity (Fig. 7B). However, this does not translate into physiological noise, which would become manifest in Panel B as stationary dots. The recurrence density of diagonal lines was lower in red RP than those in other clusters, suggesting that the red cluster experiences more dynamic state transitions. The RPs of three other cluster **(Panels D, F, H)** exhibit clear diagonal lines that repeat periodically with well-defined structure with some large gaps and elongated diagonal lines, suggesting a *quasi-periodic*, oscillatory system with phase shifts. The results of the quantitative threshold RP are listed in **Table 1**.

**Table 1.** Quantitative analysis of cluster recurrence.

| Recurrence Parameters | Clusters |
| --- | --- |

|  | Red | Yellow | Green | Blue |
| --- | --- | --- | --- | --- |
| Recurrence Rate (RR) | 3.50 | 20.49 | 18.85 | 11.60 |
| Determinism (DET) | 55.97 | 97.54 | 97.64 | 89.52 |
| Longest diagonal line length (LMAX) | 28.00 | 59.00 | 47.00 | 41.00 |
| Entropy (ENT) | 2.97 | 4.26 | 4.26 | 3.21 |
| Trend in recurrence point Density (TND) | 59.22 | 94.34 | 94.08 | 87.00 |
| Laminarity/Trapping Time (LAM TT) | 4.17 | 10.78 | 11.49 | 6.97 |
| **Total Recurrence Score** | **153.83** | **286.40** | **273.31** | **239.30** |

Note that the RPs of yellow and green cluster structure exhibited the most *deterministic* behavior with some non-stationarity that was defined from RP statistics and visualized as a strong recurrence of pattern at specific intervals on RPs. The recurrence parameters of the blue cluster were lower than those of yellow and green clusters, but higher than those in the red cluster, suggesting a higher degree of stochasticity within the blue cluster structure compared to the yellow and green clusters. In summary, RQA indicate that the yellow and green clusters exhibit more periodicity than red and blue clusters, and that the highest degree of stochasticity (lowest total recurrence score **in Table 1**) is embedded in the red cluster.

### 4.6. Ca^2+^ information sharing among SAN functional clusters

While the functional energy transfer analyses and recurrence provided a great deal of information with respect to the non-linear Ca^2+^ dynamics within each cluster (**Figs. 5-7**), these analyses do not address information sharing among SAN clusters that leads to a formation of the SAN global CaT **(Fig. 1A).**

#### 4.6.1. Pearson correlation coefficient analysis

We employed Pearson correlation analysis to determine trans-spatial relationships of LCO dynamic (first derivative) among of the normalized LCO amplitude of 50 ROIs forming four color-coded SAN functional clusters as a function of rolling window (∼15ms) throughout the first 350 ms of the time-series Ca^2+^ recordings (**Fig. S9**). We focused only on correlations coefficient that were greater than 0.99 to construct an adjacent correlation network. Prior to the initiation of each impulse (0-15ms, see vertical lines in Panel B for time reference) no ROIs were correlated at this level (Panel C). During the next 17-32-ms (Panel D), two small clusters formed near the crista terminals (red color in the ROI map in Panel A). In the following time segment (34-49ms), Ca^2+^ signal amplitudes within a large number of ROIs formed one large homogeneous cluster, a few small clusters (**Panel E**), and only 28% remained un-clustered in **Fig. S9.** During the next segments (**Panel F**, 51-66ms), the clustering patterns again changed: two major clusters and two small clusters were formed, with ∼ 20% ROIs remaining *unclustered*. During this segment, the amplitude of the integrated (global) SAN CaT rapidly increased (**Fig. S9B**). Note that the range between 34-63ms approximately encompasses the time at which the point instability had occurred in all of the clusters (Figure 3) and the time at which the attractor basin formed in the recurrence plot (Figure 4).

During the next time segments (68-84ms), one large cluster encompassing a large number of ROIs, and a one small cluster were observed, and 20% of total ROIs were *unclustered* (**Figs. S5G**). After this time segment, the global SAN CaT began to decay (**Panel B,** black trace, (85-101ms, Panel H), large clusters of ROIs in the prior segment reorganized into two medium-sized correlation clusters a few very small clusters, and ∼20% remained uncorrelated . At this time, the integrated (global) SAN CaT had decayed rapidly to ∼50 % of its peak amplitude (**Fig. S10B**). In the following time segments (103-118MS), ROI Ca^2+^ signals became largely unsynchronized, and only one medium-sized cluster and three small clusters remaining, and ∼28% of ROIs were uncorrelated (**Fig. S10**); during this time, the global SAN CaT decayed rapidly to ∼18% of its peak amplitude. The next time segments, (120-135ms, panel J) showed only a few small correlation clusters, and in the next time segment that aligned with the beginning time of next CaT (∼136-151ms) ROIs were completely uncorrelated again.

Figure **S11** shows the correlations among the kinetics of the average Ca^2+^ dynamics of the four color-coded SAN functional clusters and between them with the global SAN CaT during the entire 2.7-second time-series recording. The global CaT more strongly correlated with yellow (r=0.89) and green (r=0.98) functional clusters than with red (r=0.57) and blue (r=0.53) functional clusters (**Fig. S11**).

#### 4.6.2. Cross-recurrence Quantification Analysis

To compare interrelationship between different functional clusters, we performed a cross-recurrence quantification analysis (CRQA).[35] In contrast to standard correlation methods, CRQA provides a comprehensive evaluation of the moments when two signals exhibit recurrence, capturing synchronization patterns over time, rather than merely assessing overall similarities or dissimilarities. Since CRQA directly compares signals point by point, it can identify leading-following relationship during interaction of two signals.

Supplementary **Fig. S12** shows pairwise comparison of Ca^2+^ dynamics (rate of change, upper panels) and the respective thresholded cross-recurrence plots (TCRPs) (**lower Panels**) for all functional clusters. Cross-recurrence quantitative parameters are stated **in Table 2**.

**Table 2.** Cross-recurrence quantitative analysis (CRQA) of TRCPs between pairs of functional clusters.

| Parameters | red&yellow | red&green | red&blue | yellow&green | yellow&blue | green&blue |
| --- | --- | --- | --- | --- | --- | --- |
| Recurrence rate (RR) | 5.0 | 5.0 | 5.0 | 5.1 | 5.1 | 5.1 |
| Determinism (DET) | 67.2 | 67.6 | 67.0 | 84.8 | 81.8 | 81.6 |
| Longest diagonal line length (L_max) | 11.0 | 11.0 | 18.0 | 15.0 | 17.0 | 16.0 |
| Average diagonal line length (L_aver) | 3.0 | 3.0 | 3.2 | 3.6 | 3.5 | 3.4 |
| Entropy (ENTR) | 2.0 | 2.0 | 2.2 | 2.5 | 2.4 | 2.4 |
| Average horizontal line length (TT_H) | 3.1 | 3.0 | 3.3 | 4.3 | 4.8 | 5.1 |
| Longest horizontal line length (H_max) | 11.0 | 11.0 | 11.0 | 15.0 | 15.0 | 25.0 |

The higher the DET of a cluster pair, the more predictable the structure of the pair, suggesting that this pair *will follow* another signal rather than driving that signal. Recurrence plots for the CQRA follow the same format as those shown for the recurrence analysis in **Fig7.** The Longer L_max and L_aver between pairs, the stronger the coupling is between pairs; longer diagonal structures of pairs indicate **stronger** dynamic interactions between paired clusters; the longer horizontal lines (TT_H, H_max), the more time the cluster dwells in a given state, suggesting it might follow rather than drive overall change within the system of clusters **(Fig. S12**).

Based on our CRQA, the red functional cluster is likely the leading cluster because it has:

1. The red functional cluster is likely the leading cluster because when it is paired with any other cluster, it has: lower determinism (∼67%) across all its pairs, suggesting it is more chaotic and less structured; shorter horizontal lines, suggesting that it doesn’t stay in the same state for long time, which is typical of a driving signal.
2. The yellow, green, and blue pairs show higher determinism and more structured recurrence, suggesting that these cluster pairing may follow the red cluster, rather than lead it.

In summary, the Ca^2+^ dynamic in the red functional cluster appears to be the driver of the SAN global CaT, whereas the Ca^2+^ dynamic the yellow, green, and blue clusters follow the lead of the red cluster.

#### 4.6.3. Small-world properties and functional connectivity in SAN pacemaker activity

Small world signaling analysis, a tool within the neuroscience toolkit (REF), have been employed to search for remarkably weak ties among neuronal cells are precisely organized to maximize information transfer with minimum wiring cost. Because the cytoarchitecture and function of the SAN resembles that of neuronal tissue[17, 18], we tested whether Ca^2+^ dynamics within the system of network of SAN functional clusters exhibits small-world organization. The cluster coefficient, σ, was σ > 1 throughout the 2.7-second time-series recording, indicating the presence of small-world signaling. Figure 7 shows peak σ peak occurred at about the time of the formation of the foot of the SAN global CaT Ca^2+^, and the lowest sigma (σ) peak occurred at about the time when the global Ca^2+^ impulse achieved its peak (**Fig 8**, **Panel A**). Interestingly, the peak σ values varied from from cycle to cycle, ranging from 1 to 5 (**Figure 8**, **Panel A**).

**Fig. 8.**
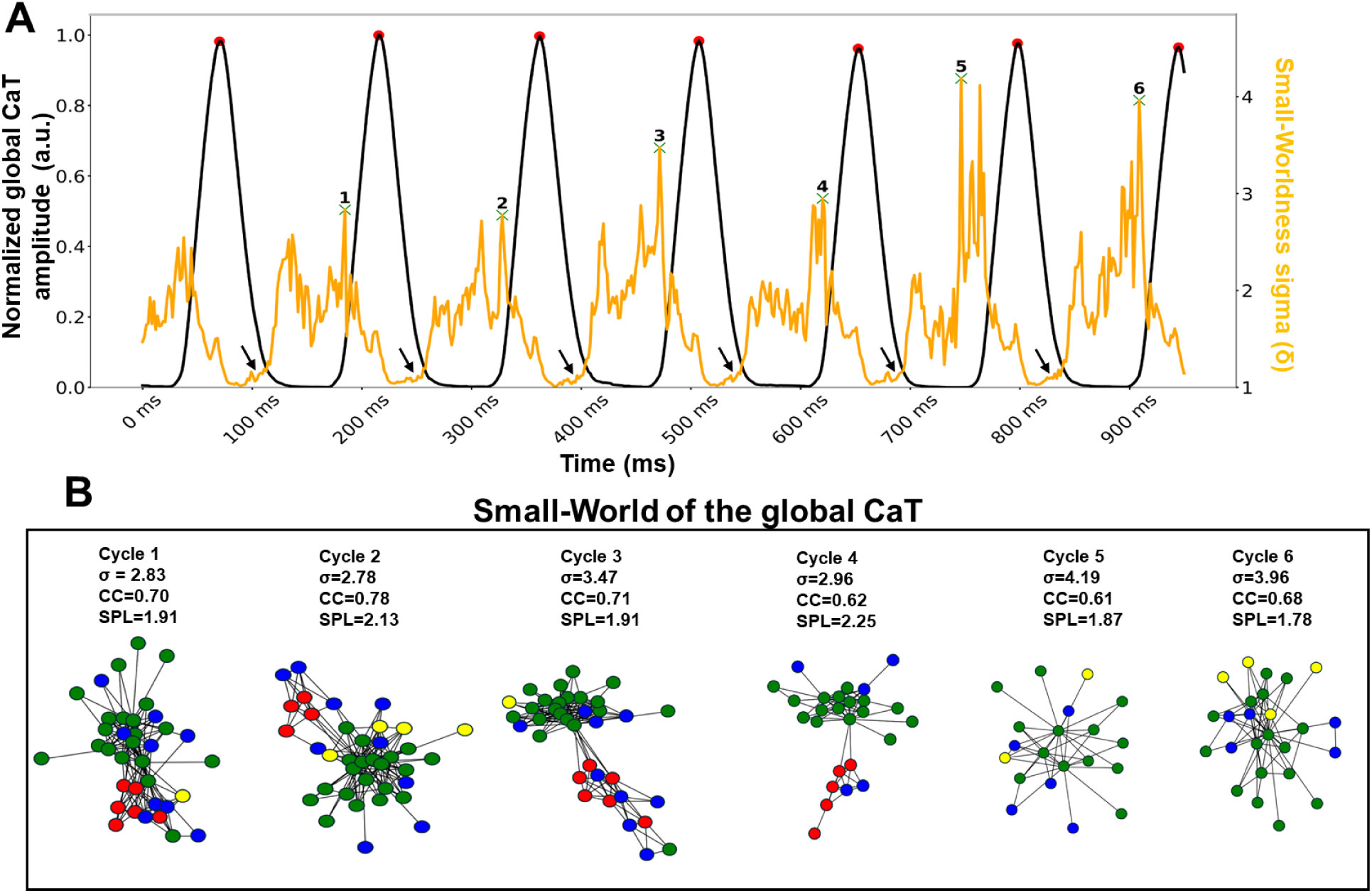
Small World Dynamics within integrated global SAN CaTs. **(A)** The normalized (0-1) global CaT signal (black color) and small-worldness, σ, (orange color) over initial 1000ms time series of a 2.7-second Ca2+ recording. Red dots mark peaks of the global Ca^2+^ impulses, while green crosses mark σ peaks. Numbers 1-6 highlight notable σ peaks between the six cycles depicted. **(B)** Small world signaling networks during each of the six consecutive cycles in Panel A, with nodes color-coded to their respective color-coded functional cluster (red, yellow, green, blue). The peak σ for each cycle is listed alongside the cycle number and the network configuration at the time of peak σ is illustrated for each cycle. Clustering coefficient (C) and average path length (L) of the network, comparing these against randomized networks that preserved the degree distribution of the original network. The small-world index *σ* = (*C*/*C_rand_*)/(*L*/*L_rand_*) where *C_rand_* and *L_rand_* represent the corresponding metrics from a random network (for 100 random constructions), was calculated for the Ca^2+^ signaling interactions between SAN tissue pixels (see methods **Figure S1** and methos **Video S2**).

Panel B shows the characteristics of the correlation coefficients (CC) and the shortest path length (SPL), point to a small-world network structure that balances efficient local information processing with rapid long-range communication. The dynamics of “small-worldness” closely align with the coordinated behavior of color-coded clusters: σ begins to progressively grow during the relaxation phase of a given SAN global Ca^2+^ impulse, and reaches its highest value around the time of the maximum 1^st^ derivative (around the peak of the red cluster) of the *next* global Ca^2+^ impulse. Thus, each cardiac impulse emerges as a unique solution of signal processing within the dynamic LCO smallworld network, known by its highly efficient information processing at low wiring and energy costs, while preserving identity (i.e. specialization) of each cluster.

In other terms, there is no time during the SAN global impulse corresponding to complete systole or diastole within or among the SAN functional clusters (the kinetics of Ca^2+^ dynamics never achieve a complete steady state, i.e., the Ca^2+^ signaling fire within the SAN *never* goes out). The animation in **Video 4** visualizes the evolving functional connections between different regions of the SAN over time and emergence of the global CaTs.

### 4.8. Is the emergence of the next SAN global CaT predetermined or stochastic?

Our Ca^2+^ dynamic analyses, **Figs. 1-8**, indicated that the SAN global CaT is a self-sustained regenerative impulse function, with the status of Ca^2+^ signals at the end of one impulse becoming the initial conditions for the next impulse, having both stochastic and predetermined characteristics. This raises the issue whether or not the SAN global CaT is predetermined or stochastic. Orbit plot analysis is a powerful visualization tool to analyze and successfully decompose the dynamics behavior within complex systems. Prior studies, utilizing orbit analysis of microelectrode recordings of SAN impulses had been interpreted to indicate that the SAN electrical impulse formation is deterministic (chaotic) in nature [36, 37]. We constructed orbit plots, which embed additional dimension of time-shifting segments (7ms) throughout 2.7 sec of Ca^2+^ dynamics[36, 38]. Fig**. S13** shows orbit plots constructed from the spatially averaged Ca^2+^ dynamics within the four color-coded clusters. This difference in the initial condition created variability of orbit plots within a given cluster, suggesting that SAN Ca^2+^ dynamic is not purely deterministic but as shown in the other analysis, recurrence and cross-recurrence analysis, partially stochastic. See supplement for further details. The Lyapunov exponent (λ), a measure of how quickly nearby Ca^2+^ dynamic trajectories in the phasespace diverge over time to estimate the degrees of stochastic and deterministic behavior embedded within a system **(Fig. S15).**

The Lyapunov exponent, λ, of the network of the SAN functional clusters was 0.082 (**Fig. S15**), intermediate between the strongly chaotic Lorenz attractor (λ = 0.146) and a noisy sinusoidal wave (λ ≈ 0.0001). This indicates moderate chaotic dynamics with sensitivity to initial conditions, but more structured than the Lorenz attractor and not purely random, validating the interpretation of the complex dynamics in stability and recurrence analyses (**Figs 6-7 and Fig. S12**), and orbit plots (**Figs. S13-14**).

### 4.9. Deduction of the SAN electrical impulse from the kinetic transitions within the SAN global Ca^2+^ dynamic

The approximate SAN electrical impulse (AP) could be deduced from the Ca^2+^ dynamics of the regional clusters and their relationship to the AP recorded with a sharp microelectrode within the right atrium adjacent to the crista terminalis (CT). **Figure 9A** shows that the peak of the atrial AP occurred slightly before the peak of the Ca^2+^ impulse of the yellow cluster, well before the completion of the Ca^2+^ dynamic of the entire SAN. The SAN electrical impulse must have occurred prior to the time at which the AP was recorded in the atrium (**Panel A**). **Figure 9A** shows the relationship of the deduced SAN electrical impulse (the dotted trace) to the measured atrial AP and to the Ca^2+^ dynamics of the SAN regional clusters.

**Fig. 9.**
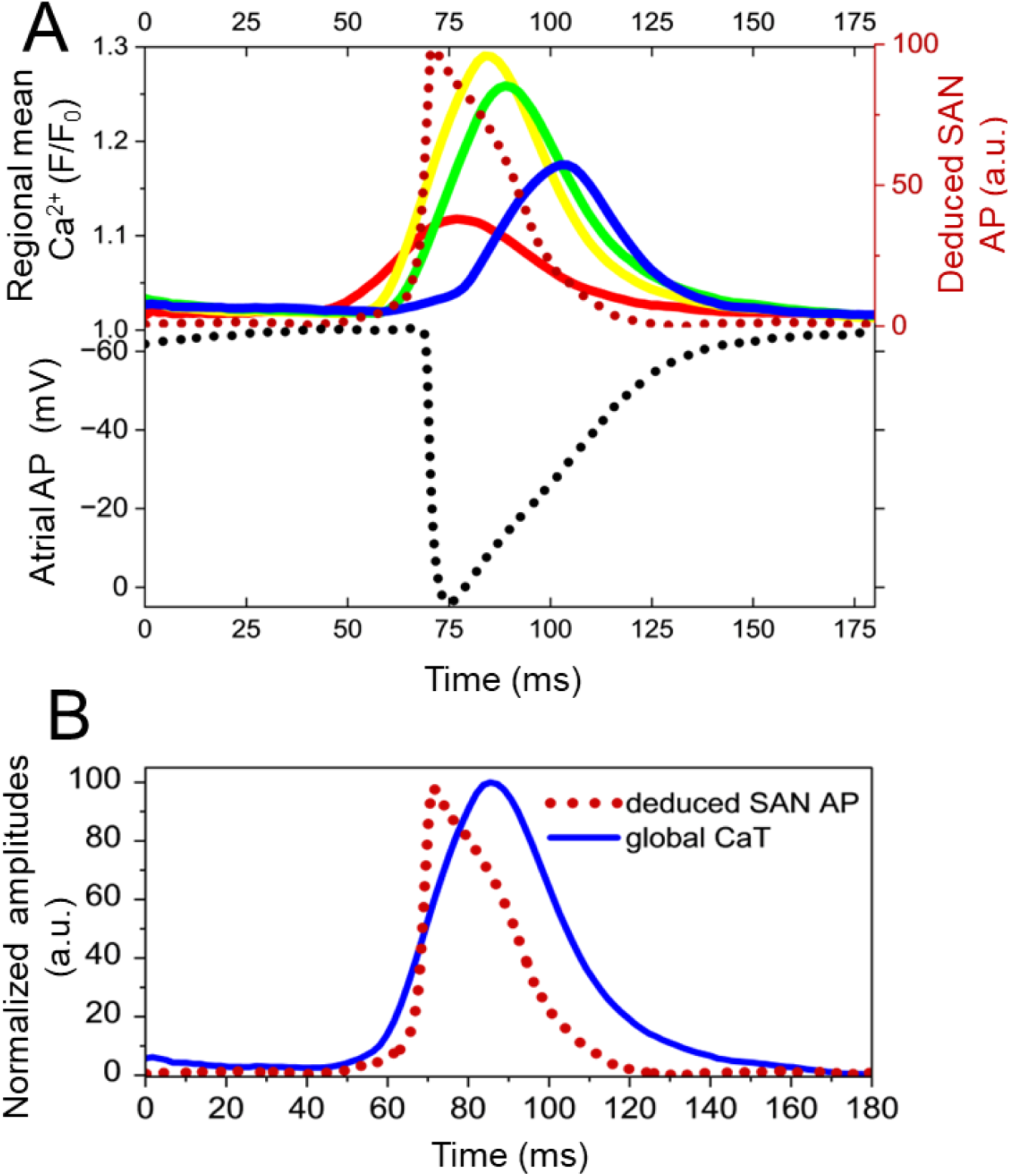
Deduction of SAN electrical impulses from measurements of the Ca^2+^ dynamics of four SAN regional clusters and membrane potential in right atrial electrode. **(A)** Relationship between deduced SAN AP and the Ca^2+^ dynamics of the SAN four regional clusters and atrial AP (measured, as a time reference point, via a sharp electrode placed in the atrium near the crista terminals recorded simultaneously with SAN Ca^2+^ dynamics. (**B)** Superimposed normalized deduced SAN electrical impulse and the global CaT.

It is evident that the Ca^2+^ dynamic of the red cluster increases slowly between ∼45-63ms, i.e., prior to the initiation of the rapid upstroke of the deduced SAN impulse. We refer to this time period as the ignition phase of the deduced SAN AP, analogous to the ignition phase (during diastolic depolarization) in single SAN pacemaker cells[39]. During this phase, there was a noticeable contribution of the other clusters within the SAN network to the deduced SAN AP, but their contributions were not as marked as the red cluster. Following this phase, the deduced SAN AP rapidly increases to its peak amplitude within approximately 4-5 ms (**Panel A**). The rapid upstroke of the deduced SAN electrical impulse is followed by (triggers) a rapid increase in the Ca^2+^ dynamic in the yellow cluster, closely followed by a rapid increase in the green cluster. The Ca^2+^ signal within the blue cluster initially slowly increased to a plateau at ∼77 ms and then, following substantial delay (∼7 ms), manifested a fairly rapid increase to a peak amplitude at ∼100 ms (**Panel A**). The rapid increase in the blue Ca^2+^ dynamic suggests that it was triggered by a *delayed* electrical impulse entering into the blue cluster. The relationship of the deduced SAN electrical impulse to the global CaT is illustrated in **Panel B**.

### 4.10 Panoramic confocal imaging of SAN, Connexin 43 and HCN4 immunolabeling

The SAN electrical impulse is initiated and conducted via translation of energy within the Ca^2+^ dynamic of loosely electrically connected SAN pacemaker cells expressing high levels of HCN4, but not connexin 43 (CX43)[17]. These studies had identified zones in which cells expressing both HCN4, and more tightly electrically coupled cells expressing CX43, are intertwined[17]. To glean spatial perspective of the SAN Ca^2+^ dynamic to the emergence of the deduced SAN electrical impulse in the present study, we preformed *panoramic* confocal imaging of SAN HCN4 and CX43 immunolabeling. The reconstructed 3D-panoramic image of dual HCN4 and CX43 immunolabeling (**Figure 10 A and B**) shows how the network of interconnected CX43^+^/HCN4^−^-immunoreactive cells engages the meshwork of the HCN4^+^/CX43^−^ -immunoreactive pacemaker cells. CX43 antibody-labeled membrane proteins observed as green dots in **Fig. 10** align at the perimeter or at the ends of cells and define cell borders. Postprocessing of the series of Z-stack images with orthogonal sectioning extracted 3D-images of the Cx43^+^-“sleeves” intertwined with pacemaker cells meshwork in the endo-epicardial projection planes (**Fig.10, Panels C and D**).

**Fig. 10.**
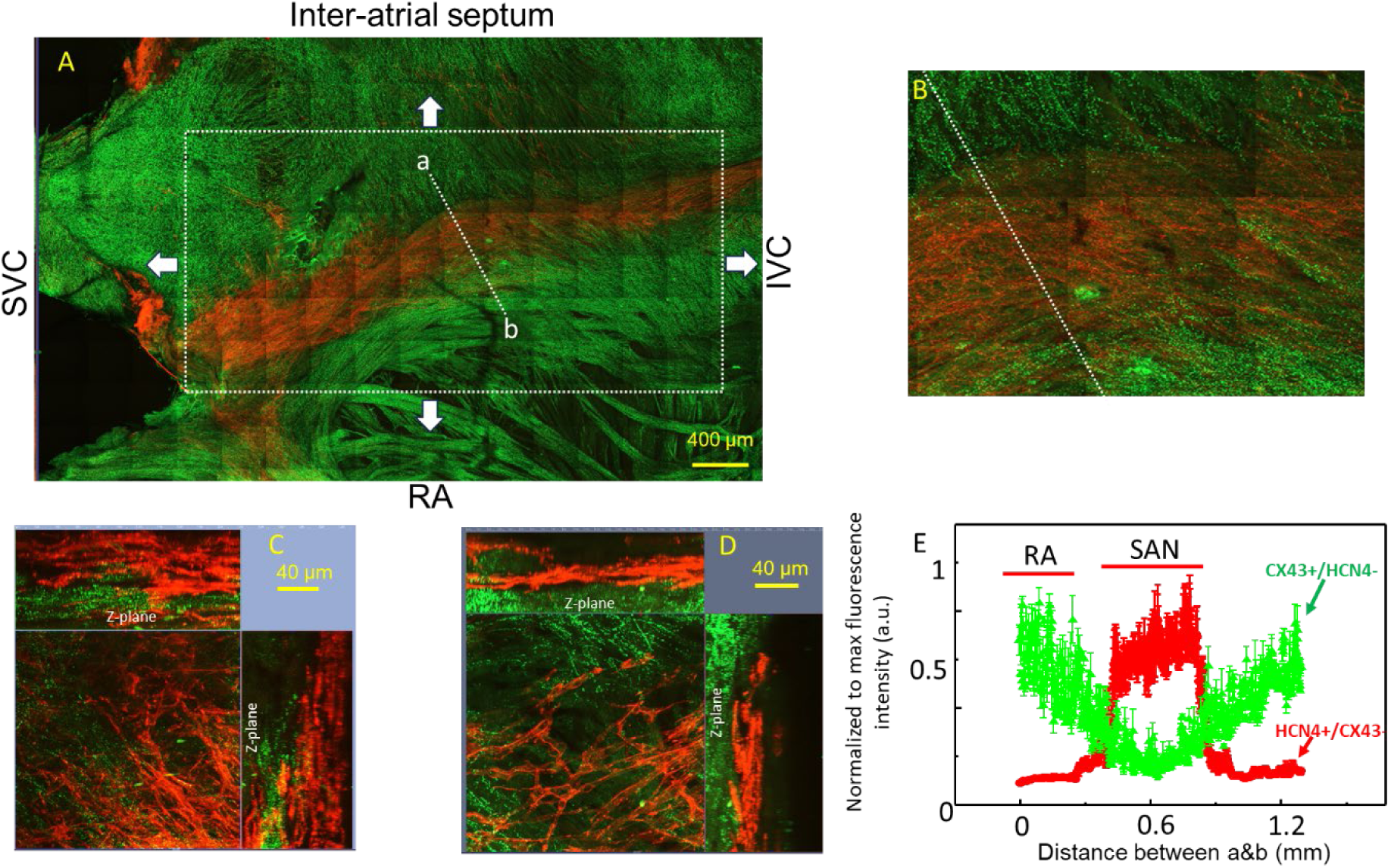
Panoramic 3D confocal SAN image reconstruction. **Panel A:** A panoramic 3D confocal image reconstruction of the meshwork of HCN4^+^ (**red**) expressing pacemaker cells that do not express CX43, embedded within a network of CX43^+^ (**green**) cells that do not express HCN4^−^. **Panel B** is zoomed **Panel A**. **Panels C** and D: images of reconstructed virtual cuts of intertwining HCN4^+^/CX43^+^ zones within SAN tissue from the endocardial to the epicardial surface. The image in **Panel C** is from the central part (the “body”) of the SAN, and **Panel D** is from the tail of the SAN, close to inferior vena cava. SVC-superior vena cava, IVC-inferior vena cava, CT-crista terminalis. RA-right atrium. Panel E: the region of interest in Panel A containing the dashed white line is digitally amplified. The box encompassed by the dashed white line, shows the area in which Ca^2+^ responses illustrated in prior Figs. 1-9 were recorded, normalized to the maximum amplitude. **Panel E** shows the fluorescence, normalized to their maxima, the CX43 (green) and HCN4 (red). The dashed white line shows the optical cut in which HCN4 and CX43 immunolabeling was quantified in this whole mount preparation.

We refer to areas in which HCN4^+^ and CX43^+^ immunolabeling overlap to be an “intertwining zone,” and envision that within this intertwining zone, an electrical impulse initiated within the loosely, electrically coupled HCN4^+^ pacemaker cells engages the CX43^+^ network to initiate the SAN AP. The average density (normalized to maximum) of CX43 (green) and HCN4 (red) protein immunolabeling, measured across a distance beginning from the end of the dashed line within the atrial trabeculae to the center of SAN (Panel A) in 3 different preparations is shown in **Fig. 10E**. The overlapping of HCN4 and CX43 fluorescence intensity reflects the gradient in CX43 and HCN4 protein expression within a given locus. Because the density of the CX43 expression and degree of SAN intercellular electrical coupling are linearly connected[40], indicates that as the density of CX43 expression increases from the central SAN to the atria, the level of electrical coupling among those cells increases proportionally, facilitating rapid propagation of electrical signals away from the SAN center.[40]

## 5. Discussion

We applied novel linear and non-linear analytical paradigms to the time-series of SAN Ca^2+^ images to extract a comprehensive profile of how heterogeneous (stochastic) local Ca^2+^ oscillations (LCO) in and among SAN pacemaker cells (see Video S1) become synchronized and desynchronized, forming rhythmic SAN impulses (global CaTs).

Our results indicate that information processing within the LCO ensemble is linked to the formation of incessant highly rhythmic, albeit unique SAN impulses. Phase analysis of local Ca^2+^ dynamics in isolated mouse SAN delineated a network of functional pacemaker cell clusters, distinguished by their local Ca^2+^ dynamic amplitudes, kinetics, and phases (**Figs. 2-3, 5, S2, Video 1-2**). Cross-talk of local Ca^2+^ dynamics within the network (**Figs.8, S5-6, S9-10, Video 4**) culminated in rhythmic SAN global Ca^2+^ transients (CaTs), having a mean rate and rhythm identical within a given SAN to that recorded by a reference sharp electrode in the right atria (**Fig. S1A**), indicating that CaTs are induced by global SAN electrical impulses.

A number of detailed non-linear analyses indicated the presence of hidden structures within each cluster that differed from that of other clusters (**Figs. 5-7, S12-14, Table 1**). Initial conditions of each impulse initiation and subsequent LCO ensemble evolution during each cycle differed from each other, due, mostly to a stochastic process (having a degree of uncertainty) of the Ca^2+^ dynamics within a small pacemaker cluster located near the crista terminalis (**Figs. 4, 7-8, S12, Table 1-2**). That cluster exhibited the highest degrees of intrinsic power, earliest rotor-like energy transfer, most frequent point-to-point instability (**Figs. 3-5, S4**), earliest acrophase (**Fig. 2**), and greatest impulseto-impulse variability within the SAN cluster network (**Fig. S1**). Further, cross-talk among *individual signaling clusters* varied during each cycle, forming a unique sequence of small-world Ca^2+^ signaling networks which in turn generated *a unique* spatiotemporal solution for Ca^2+^ dynamic synchronization linked to each global CaT (**Fig. 8, Video 4**). Based on Ca^2+^ dynamic analyses of the regional clusters and their interactions, we deduced the SAN electrical impulse (AP) (**Fig. 9**). Emergence of SAN impulses from local Ca^2+^ dynamics recapitulates, at a higher level of organization, the emergence of spontaneous impulses in isolated, single, SAN pacemaker cells shown in prior studies [8–11, 41–45].

### 5.1. Regional energy, frequency distributions, point stability transitions of regional Ca^2+^ dynamics within the non-liner global SAN Ca^2+^ dynamic

Initial energy expenditure began as a rotor-like trajectory of acceleration and velocity within the Ca^2+^ dynamic in each cluster, and was reflected initially as a state of low energy transfer, which then progressively increased during the impulse formation (**Fig. 5**). The greatest energy expenditure occurred during the rapid rate of increase of the global CaT (**Figs. 1, 2, 5**, **S7-8)** and was reflected in a greater expansion of the rotor-like trajectory acceleration and velocity in all clusters.

The rotor appeared first within the cluster (red cluster) (**Fig. 5D**) located near the CT that initiated the ignition phase (foot) of the global CaT. This cluster had the *highest* power and oscillated around a slightly higher dominant frequency and greatest degree of stochasticity than that of other regional functional clusters (**Figs. 3B, 6-7, Table 1**), resembling Ca^2+^ “hot spots” that have been observed in other tissues [3, 46, 47]. The characteristics of this (red) SAN cluster may be likened to pacemaker/rhythmogenic neurons in the Central Pattern Generator networks of the central nervous system, similar to those recently identified to have a key role in regulating the zerbrafish heart rhythmic functionality[48].

The slower velocity at which the Ca^2+^ dynamic increased within this cluster compared to that of other clusters within the network, likely reflects a lower degree of coupling of pacemaker cells within this cluster to each other compared to other clusters. Due to its small spatial area the Ca^2+^ dynamic within this cluster (**Fig. 2A**, red), achieved a lower amplitude (c**.f. Fig. 2B**) compared to that of other regional clusters within the SAN cluster network. Indeed, confocal SAN Ca^2+^ imaging in prior studies discovered that the area in which the earliest Ca^2+^ impulse could be detected was near to the CT, as in the present study, and expressed HCN4, but not Cx43[18]. Although the *total* energy expenditure during self-ordering of spontaneous LCO’s to form the Ca^2+^ dynamic of the red cluster was substantially less compared to than that of other regional clusters (**Fig. 5C-F**), it was sufficient to drive the rate and rhythm of the Ca^2+^ dynamics in other clusters, in which energy expenditure during the formation of a CaT was much greater.

The appearance of the rotor in the red cluster was followed by the appearance of the rotors in the other clusters (**Fig. 5**). Initiations and expansions of rotor-like trajectory in each cluster varied during different SAN global CaTs during a time-series (**Fig. 5C**). Importantly, energy transfer within the SAN Ca^2+^ dynamic never achieved true equilibrium in any cluster during a time-series of impulses (The SAN Ca^2+^ “fire” never goes out!).

### 5.2. Cross talk among SAN functional cluster Ca^2+^ dynamics creates the global SAN Ca^2+^ dynamic

Transitions in stability and energy of Ca^2+^ dynamics among clusters mirrored each other following time delays (**Figs. 4-5**). Importantly, point stability oscillations and energy transitions of the Ca^2+^ dynamic within the red cluster had begun to initiate the *subsequent* CaT, even before such transitions in other clusters had fully stabilized from their trajectories within the *prior* CaT (**Fig. 4**). Different phase relationships within and among the SAN functional cluster network during the time period encompassing at the end of a previous cycle and the beginning of the next cycle, created variable initial conditions of each SAN global CaT. Therefore, ourresultssuggestthatcollectiveintelligence[49]across multiple SAN functional components work together within a network to create global SAN impulses. Cross-talk among the Ca^2+^ dynamics within *different clusters* varied throughout an impulse and from one impulse to the next. This cross-talk formed unique, dynamic small-world Ca^2+^ signaling networks, creating *a unique* spatiotemporal Ca^2+^dynamic synchronization solution for each global CaT (**Fig. 8**). PCA indicated that all clusters within SAN network contributed to the SAN impulse, but the magnitude of each contribution varied during the time of the impulse formation (**Figs. S5-S6**), with the small cluster near the CT (color-coded as red), as noted above, contributing to the majority of the energy during the pre-foot and foot of the global CaT prior to the rapid upstroke (**Fig. S5B-D**). Pearson correlation analysis showed that time-dependent synchronization/desynchronization of the Ca^2+^ dynamics within each cluster formed global Ca^2+^ impulses within the time-series (**Fig. S9-S10**); and within a given temporal epoch during an impulse, the kinetics and amplitudes of functional Ca^2+^ dynamics in *different* regional pacemaker cells clusters, became correlated with each other. This pointed to the creation of a time-dependent, correlated *network function among the clusters*. The magnitude of correlations also differed between clusters, e.g. the yellow and green strongly correlated to each other than to other clusters; the red and blue clusters negatively correlated with each other, consistent with different functional properties of regional clusters (**Fig. S11**).

The existence of small-world signaling throughout the Ca^2+^ time-series recording (**Fig. 8A**), suggests that the Ca^2+^ dynamic network architecture is capable of balancing local connectivity with global integration, ultimately forming *a unique* spatiotemporal LCO synchronization solution to create each impulse (**Fig. 8B).** This behavior of SAN tissue resembles that of functional brain networks in which information transfer is maximized with minimal wiring cost by self-similar modules of small world weak ties[50].

Earlier studies, based on orbit plots, concluded that SAN electric impulses were purely deterministic[36, 37]. Orbit and recurrence plots of Ca^2+^ dynamics of the present study (**Figs. S6-7, Figs. S13-14)**, however, reveals the SAN network nonlinear Ca^2+^ dynamic features both stochastic and deterministic characteristics that differ from one impulse to the next. The difference in the interpretation as to whether spontaneous SAN impulses are purely deterministic, or in part stochastic and in part deterministic is due to the failure of assessment of Ca^2+^ dynamics in the prior study [36] . Indeed, recurrence and cross-recurrence analysis of regional Ca^2+^ dynamics (**Figs. 6-7, S12, Tables 1, 2**) indicated that the stochastic characteristics within the global SAN Ca^2+^ dynamic within time-series likely stems mostly from the stochastic nature of LCOs in the red compared to other clusters and that the deterministic-like features are in parted by the Ca^2+^ dynamics of the other clusters (yellow, green, blue).

The Lyapunov exponent of SAN Ca^2+^ dynamics (**Fig. S15)** also strongly suggests that the SAN Ca^2+^ signaling dynamic exhibits chaotic features, though less sensitive to initial conditions than a purely deterministic system dynamic. In other terms, our SAN Ca^2+^ dynamic is intermediate between highly periodic and stochastic. The stability of SAN impulse formation within a time series, in the face of both chaotic dynamics and stochastic processes is achieved through the bounded nature of the system; while individual cycles may vary due to both deterministic chaos and random fluctuations, the overall behavior remains within certain physiological limits. This delicate balance between chaos, stochasticity, and stability allows the heart to respond flexibly to physiological demands while maintaining overall reliability, a perfect arrangement for the critical task of regulating the heartbeat.

### 5.3. Potential mechanisms of the non-linear SAN Ca^2+^ dynamic

Numerous potential mechanisms that underlie the non-linear SAN Ca^2+^ dynamics that confer both its stochastic and deterministic characteristics (memory) are likely related to these nanoscale mechanicalelectro-chemical-thermal factors[9, 51, 52] impacting molecules of the coupled-clock system within SAN pacemaker cells, and molecules within the large number of interstitial cells residing within the SAN[18]. Different cells may express different levels of Ca^2+^ signaling proteins, e.g. ion channels and pumps, which can affect the response to stimulation and the dynamics of Ca^2+^ signaling. Even within the same cluster, there may be variations in the distribution of inputs from neighboring cells or regions, leading to differences in the spread and strength of the Ca^2+^ signal, resulting in variable trajectories of Ca^2+^ dynamics. Feedback and cross-talk between pathways can result in variations in the response of different cells within a given cluster. Environmental factors e.g., local differences in temperature, pH, or oxygen levels can also affect the dynamics of Ca^2+^ signaling. Studies aimed to more fully define these factors and specifically how they impact the overall coordination and variability of Ca^2+^ signaling within the complex SAN cellular network ought to be given high priority in futures studies of SAN impulse formation.

The apparent memory encoded within stochastic and deterministic characteristics within the network of SAN regional clusters is similar to a type of memory in single SAN pacemaker cells that is attributable, in large part, to the Ca^2+^ content of the SR, the Ca^2+^ clock within the coupled clock pacemaker system[9], the impact of Ca^2+^ released from the SR on target molecules within the membrane clock that are activated by Ca^2+^. Because synchronization of activation states of coupled clock molecules is modulated by phosphorylation, the phosphorylation status of clock molecules modulates both the availability of the number of Ca^2+^ ions to be pumped into the SR and the rate at which Ca^2+^ is released via ryanodine receptors (RyR)[9], thus, modulating the SR Ca^2+^ content. At the initiation of each AP cycle, Ca^2+^ leak from the SR creates local, spontaneous Ca^2+^ oscillations (LCOs), that self-organize into an ensemble Ca^2+^signal that interacts with membrane clock molecules to create spontaneous diastolic depolarization of the cell membrane. This is followed by a synchronized release of Ca^2+^from the SR that is triggered by the rapid upstroke of the SAN AP. Some of the Ca^2+^ released from the SR is extruded from the cell, while Ca^2+^ ions enter the cell through other ionic mechanisms during each AP cycle. But the SR Ca^2+^ content (memory stored in the SR), even in an apparent quasi-steady state, is not exactly the same from one AP cycle to the next[45]. Thus, cycle-to-cycle variability in spontaneous LCO characteristics and the ion channel current activation reflects cycle-to-cycle variability of the memory within the coupled-clock system in SAN pacemaker cells[45]. Cycle-to-cycle variations in memory encoded within SAN pacemaker cells embedded in SAN tissue is a plausible explanation for why energy transfer trajectories or orbit plots do not return to the exact initial points, and manifest cycle-to-cycle variability[45] in a given apparent quasi-steady state throughout time-series Ca^2+^ recordings.

### 5.4. SAN electrical impulses deduced from the complex non-linear SAN Ca^2+^ dynamic

The approximate phases of the SAN electrical impulse can be deduced from the complex non-linear SAN Ca^2+^ dynamic (**Figs. 5-9, Figs. S12-S15**) from the kinetics and amplitudes of Ca^2+^ dynamics within the SAN cluster network (**Figs. 1-4, S3-S5**). Incessant, spontaneous, spatiotemporal synchronization of LCOs within a small cluster (color-coded red) made the major contribution to the foot of the global CaT (**c.f. Figs. 2-9, Figs. S4-S6, S12-S14**).

The similarities of the initial slow foot of the global SAN CaT and the initial slow foot of the Ca^2+^ transient in single pacemaker cells, suggests that: 1) although not directly measured in the present study, the integrated low-amplitude initial Ca^2+^ signaling creates the concomitant increase in the deduced SAN with the red cluster that formed the foot of the global SAN CaT (**c.f. Fig. 2**, colorcoded red).

Thus, the deduced slow increase in the local depolarizing electrical field potential (LFP) during the foot of the global CaT in SAN tissue is analogous to the slow diastolic depolarization aka, the “pacemaker potential”[41], in individual cells that emerges from self-organization and synchronization of heterogeneous LCOs.

It is reasonable to speculate that a small LFP generated largely by Ca^2+^ dynamics within a small, red cluster of SAN pacemaker cells, initiates the diastolic depolarization (DD) of the SAN AP. Indeed, a slowly rising small foot of SAN AP (coinciding with the formation of the slow foot of the SAN global CAT in the present study) had previously demonstrated directly in studies using voltage-sensitive fluorescent indicators[53, 54]. Panoramic tiled confocal SAN images of the present study indicate that beyond a zone in which HCN4 and CX43 expressing pacemaker cells intertwine the density of the density of CX43 expression increases (**Fig. 10**). It is also reasonable to speculate that an LFP, forms within the intertwining meshwork of HCN4^+^/Cx43^−^ expressing cells[17] and HCN4^−^/Cx43^+^ expressing cells. As the amplitude of the Ca^2+^ dynamic, largely within the red cluster, continues to exponentially increase, a rapid depolarization of the LFP within the intertwining zone creates the rapid increase in the electrical impulse in the adjacent clusters (yellow and green), and the subsequent rapid phase of the SAN global CaT, due to an increase in the amplitude of the Ca^2+^ dynamic of the yellow and green clusters, is triggered by the rapid upstroke in the SAN electrical impulse (AP) (**Fig. 9A**).

The rapid acceleration of the deduced SAN electrical impulse was very brief, achieving a peak in ∼5ms from its onset, and occurred prior to the time at which the peak of the global CaT was achieved (**Fig. 9B**), approximately coincided with the time at which the peak of AP was recorded in the right atrium (**Fig. 9A**). This is in agreement with observation that electrical impulses, measured via voltage-sensitive fluorescent indicators in mouse and the SAN of other species activate the atria prior to the time at which the SAN AP is completed[53]. The SAN electrical impulse is conducted away from the intertwining area in the direction in which density of Cx43 expression (electrical conductance) progressively increases (**Fig. 10**). The non-radial spread of the SAN Ca^2+^ dynamic observed here, would be expected to result from the non-radial SAN AP spread from the leading pacemaker site described previously[55].

Interestingly, the timing and kinetics of the Ca^2+^ dynamic within the blue SAN functional cluster, approximately located in the mid-SAN, and extending toward the septum (**c.f. Fig.2A, colorcoded blue**), differed from those of each of the other clusters. Specifically, the Ca^2+^ dynamic of the blue cluster began, as in the red cluster, to slowly increase during the ‘pre-foot’ of the global SAN CaT (as clearly seen in **Fig. 2D-E, blue**), continued to increase slowly, similar to that of the yellow and green clusters, but then plateaued (stalled), as if the conduction of the SAN electrical impulse that triggered a rapid increase in Ca^2+^ in the yellow and green clusters were “blocked” from entering into the blue cluster (**Fig.2**). Indeed, the location of this cluster within the SAN approximated that, which in prior studies, has been referred to as a ‘block zone’, into which the SAN electrical impulse inmany species fails to be conducted[56–58]. Although bands of fibrous tissue, or branches of the SAN arterial system are often-cited causes of impulse block into this area of the SAN [57]. Poor excitability of cells within this area is likely to be the functional mechanism for an apparent block of impulse spread into this area [57, 59]. The peak of the Ca^2+^ dynamic of this cluster also occurred beyond the peak of the global SAN CaT (**Fig. 2**). An increase in Ca^2+^ during this time may contribute to the repolarization of the SAN AP, via impacting on Ca^2+^-dependent K^+^ channels[60–62] and other Ca^2+^-dependent electrogenic surface membrane molecules.[63]

### 5.5. Is reentry into the SAN/medial right atrium an intrinsic component of normal SAN electrical impulses?

After a substantial delay, however, the Ca^2+^ dynamic of this cluster began to increased rapidly, as if it were triggered by a delayed AP that had been conducted into this cluster, (in the yellow and green clusters), reaching its peak amplitude at a time that was well beyond the time at which the peak of the AP reference electrode in the atrium was recorded. Sinus node reentry in the rabbit heart has been previously directly demonstrated in response to an ectopic beat[64, 65]. Delayed excitation of the blue cluster may have resulted from an electrical input originated within the interventricular septum, reentering the blue cluster, possibly via precaval bundles or Bachmann’s bundles, the later being the main pathway via which excitation of the left atrium occurs[66, 67]. Although excitation of the left atrium is delayed compared to that of the right atrium, contraction of both atria occurs nearly at the same time, possibly due to a delayed impulse into the SAN blue cluster [66, 68–70]. As noted previously, it is not possible to define the exact anatomic medial border of the SAN [53, 56, 58, 67] and it is possible that the blue cluster demarcated in the SAN image studied in the present work is actually part of the right atrium, and not part of the SAN.

If a conduction block of the impulse *into* the blue cluster were to be complete and bi-directional, a delayed APoccurrence in his cluster might not be expected to be conducted back into the adjacent (red) cluster, and would, therefore, not be expected to impact the ignition process of the next global CaT (occurring at this time in the red cluster, **Fig. 2**). In other terms, complete failure of a delayed reentry of an electrical impulse from the blue into the red cluster, would prevent the creation of a complete reentrant circuit that would lead to (“or dictate”) a stereotyped timing of the initiation of SAN global CaTs. However, impulse to impulse variability in the initiation phases of orbit plots (**Figs. S12-S13**), energy transfer in FDA (**Figs. 5**), and variability of point instability of the Ca^2+^ dynamic in the red cluster **(Fig. 4),** creating variability in timing of the initiation of subsequent global CaTs (**Fig. 1, Figs. S1, S8)**, strongly suggest that reentry of a stereotyped electrical impulse into the red cluster does not appear to have a *major* impact on the synchronization of LCOs in this cluster that initiates the next SAN impulse.

It is possible, however, that the functional block during the early SAN impulse phase is *unidirectional*, and that *variable penetration* of a strong electrotonic influence, such as an AP occurrence within the blue cluster at a later time in the cycle, might influence the LCO dynamic within the red cluster. If this were to be the case, a *parasystolic-like* micro reentry mechanism [71], might modulate the Ca^2+^ dynamic of the red cluster and therefore modulate the timing of the which the next impulse would occur. “Normal” mouse impulse to impulse SAN variability, in this instance, would reflect, not only variability in the kinetics of incessant, spontaneous resynchronization LCOs within the red cluster, but also to variable penetration of electrical impulses from the red to the blue cluster. Future studies are required to inform on the validity of this speculation.

### 5.6. Limitations and segue to future research

A limitation of the present study, as in prior studies with microelectrodes and voltage-sensitive fluorescence probes, is that Ca^2+^ dynamics were assessed in 2D. Presently, however no techniques to measure Ca^2+^ in 3D with appropriate time sampling commensurate with Ca^2+^ kinetics.

### 5.7. Clinical Perspectives

In order to understand how pathophysiological mechanisms impact on SAN function, leading to arrhythmias associated with aging and cardiac disease, its necessary to understand how SAN impulses are normally generated in the absence of these conditions. The present study provides novel perspectives on the Ca^2+^ dynamics involved in the SAN impulse formation in mice. Because SAN structure and function in mice are similar to those in other species, including humans[57], perspectives gleaned from how Ca^2+^ dynamics within a network of SAN functional clusters differentially contribute to the formation of SAN global impulses in mice are of clinical relevance with respect to:

- To understand how responses to stimulation or inhibition of autonomic receptors, and other neurotransmitters cause changes in the rate and rhythm at which SAN impulses are generated
- Mechanisms of initiation and maintenance of SAN reentrant arrhythmias
- Pathophysiology of SAN aging, the emergence of sick sinus syndrome.

Disruption in the origin of electrical impulses in the SAN underlie numerous types of cardiac arrhythmias. Prior studies of arrhythmias have generally utilized linear approaches to analyze SAN impulse formation. Present results show how nonlinear analysis can easily been applied reliably to detect and visualize hidden functional structures within the SAN that are essential for the early detection and diagnosis of many arrhythmias.

## Institutional Review Board Statement

The animal study was reviewed and approved by the Animal Care and Use Committee of the National Institutes of Health (protocol #457-LCS-2027).

## Abbreviations

AP: action potential
Acrophase: period from a reference point to the maximum value of a signal
CaT: action protentional-induced Ca^2+^ transients
Chronopix: chrono-pixel
CT: crista terminalis
CX43: Connexin 43 (Gap junction alpha-1 protein)
dCaT/dt: 1^st^ derivative of CaT (1^st^ derivative, velocity)
d2CaT/dt2: 2^nd^ derivative of CaT (2^nd^ derivative, acceleration)
d3CaT/dt3: 3^rd^ derivative of CaT (3^rd^ derivative)
IROC: instantaneous rate of change
F/F0: normalized Ca2+ fluorescence (max/min)
FDA: functional data analysis
Functional clusters: regions of interest with similar Ca^2+^ dynamics
HCN4: hyperpolarization-activated cyclic nucleotide-gated channel Ignition phaseslow foot of each CaT
IVC: inferior vena cava
IMF: empirical mode function
LCO: spontaneous, stochastic, local Ca^2+^ oscillation
Lyapunov exponent λ: defines the rate of separation of two initially close trajectories
NCX: Na+/Ca2+ exchanger
PCA: principal component analysis
ROI: region of interest
SAN: sinoatrial node
sigma (σ): small-world metric that relates the clustering coefficient to the shortest path length
Small-world network: quantifies efficient information seclusion and integration
SVC: superior vena cava

## Supplementary Materials

### 2. Methods

#### 2.1. Sinoatrial node tissue (SAN) preparation

Our experiments conformed to the Guide for the Care and Use of Laboratory Animals, published by the US National Institutes of Health. The experimental protocols were approved by the Animal Care and Use Committee of the National Institutes of Health (protocol # 457-LCS-2027).

The sinoatrial node (SAN) tissue was dissected from the heart of young (3 month-old) C57BL/6J mice (Charles River Laboratories, USA) (n=4) as previously described ^1^. Mice were anesthetized with Isoflurane (inhalation at 2-3% for induction and 1-1.5% for maintenance) and monitored until the loss of reflexes to tail pinch confirmed the adequacy of anesthesia. The heart was rapidly excised and immediately transferred to a cold (4°C) oxygenated standard Tyrode solution containing (in mM): 130 NaCl, 24 NaHCO3, 1.2 NaH2PO4, 1.0 MgCl2, 1.8 CaCl2, 4.0 KCl, 5.6 glucose, equilibrated with 95% O2 / 5% CO2 (pH 7.4 at 35.5°C). The heart was pinned onto a silicon platform, and the right atrium was carefully opened to expose the SAN region, including the crista terminalis, intercaval area, and interatrial septum, with the superior and inferior vena cava (SVC and IVC) kept intact. The preparation was then pinned to the bottom of a *oxygen*-controlled experimental chamber with the endocardial side facing upward, ensuring minimal stretch to flatten the SAN tissue.

#### 2.2. Data acquisition of local Ca2+ signals/oscillations (LCO) during impulse formation in SAN ex vivo

The SAN preparation (n=4) was incubated with Calbryte 520 AM (25 μM) a fluorescent calcium indicator, for 1.5 hours at room temperature. After washout of dye for 20 min, local Ca2+ signals (LCOs) were recorded in Tyrode solution (as above) with 5x magnification at 35.6C and acquisition rate 1.7ms for 2.7 sec via a stationary fixed stage upright microscope (Axio Examiner D1 equipped with zoom tube (0.5–4x), Carl Zeiss Microscopy LLC) and PCO edge 4.2 camera as previously discussed ^1^. The epifluorescence microscope with a CoolLED pE-300ultra light source was set to 40% power to optimize signal-to-noise ratio and maintain tissue viability. The excitation light was directed through a 498 nm dichroic mirror, and emitted fluorescence was collected through a 530±20 nm filter.

To minimize motion artifacts during recordings, preparations were treated with 10 μM cytochalasin D to inhibit mechanical contraction ^1^. Residual motion artifacts were corrected during post-processing using ImageJ. The high temporal (500–700 Hz) and spatial resolution allowed for the construction of spatiotemporal maps of SAN cell network activation. Ca2+ signal intensity remained stable for *the entire experiment*.

#### 2.3. Atrial cell action potential recording

Because action potential (AP) characteristics differ in the SAN pacemaker cells among regions of the SAN ^1^, to deduce the timing of SAN action potential (AP), we placed reference electrode in atria near the crista terminalis (CT). To record the transmembrane potential (AP), we used sharp microelectrodes (40–100MΩ) fabricated from aluminosilicate glass capillaries (1.5mm OD 0.86mm ID) with a horizontal pipette puller (Sutter Instruments, CA, USA), and back-filled with 3M KCl. Each microelectrode was inserted from the endocardial side of the tissue, and APs were recorded with a high-impedance amplifier (A-M Systems, USA) with a virtual bridge circuit for current injection. The electrical signal was digitized with Digidata 1440 and analyzed with PCLAMP10 software (both from Molecular Dynamics, USA).

#### 2.4. Data analysis

Original Ca2+ signals recorded during 2.7 sec were preprocessed in ImageJ and then a local Ca2+ signal pixels intensity (LCO) within the image was extracted by applying a 50 equaled size grid (68×100 pxl). Next, the smoothing of the extracted LCO data was performed, using Savitzky-Golay method, to remove noise or irregularities in the of the origin LCO signal, followed by normalization of the maximum LCO signal of each of the 50 ROI to its minimum signal (F/Fo) over time. The Ca2+ signals were presented as F/Fo values or as a percent of LCO change in each ROI (F-min(F))/(max(F)-min(F)). Time series were generated from the space-averaged LCO signal within each ROI for each frame in the sequentially recorded stack of images. When necessary, a first-order differencing of time series was applied to transform to transform nonstationary time series into stationary ones.

We applied a diverse set of linear and nonlinear analytical analyses of local Ca2+ (LCO) dynamics using multiple software programs: Microsoft Excel, OriginLab, MATLAB 2024a, R, pCLAMP 10.4, our custom-made programs to analyze APs and Ca2+ transients dynamics (Victor Maltsev).

##### 2.4.1. Cosinor-based analysis

(Germaine Cornelissen) was applied to measure the acrophases in each ROI, the times from a reference point (0) to the time of maximum amplitudes, during the first 200 ms of impulse formation to pinpoint initiation of Ca2+ signal change among ROIs.

##### 2.4.2. Amplitude and derivative analyses

were applied to assess heterogeneity in amplitudes and kinetics of LCOs during different phases of impulse formation within and between LCOs.

##### 2.4.3. Tangent analysis

was applied to identify the slow phase of the global Ca2+ signal.

##### 2.4.4. Pearson correlation analysis

was applied as a function of window rolling (∼15ms) to the normalized LCO amplitude of 50 ROIs to identify the strength of a linear correlation between LCOs of ROIs.

##### 2.4.5. Network cross-correlation analysis was

applied to identify functional LCOs clustering patterns (network solutions) within and between LCOs (r =>0.99, Fruchterman–Reingold Force-directed plots) during impulse formation.

##### 2.4.6. Functional data analysis (FDA)

was applied to illustrate the nonlinear dynamic of the energy transfer in the Ca2+ signal during impulse formation.

##### 2.4.7. Embedded phase-plane analysis

was applied to study network solutions in SAN global and regional Ca2+ signals by assessing the deviation from the limited cycle and the presence of chaos during each Ca2+ impulse ^2,3^.

##### 2.4.8. Recurrence Characteristics of the SAN Ca^2+^ Dynamic System

We performed Recurrence Quantification Analysis (RQA) on the four regional cluster Ca^2+^ dynamics via a modified MATLAB toolbox ^4,5^. RQA detects hidden patterns and recurrences in a time series without assuming signal stationarity, making it particularly useful for short time series and discontinuous data. To perform RQA a one-dimensional time-series data initially projected into a multidimensional state space, representing the system’s state at a given time, prior to transforming it into a recurrence plot (RP) using time-delay embedding. The RP visually represents the results of the RQA, i.e., quantification of patterns through specific metrics such as recurrence rate (RR), determinism (DET), longest diagonal line length (LMAX), entropy (ENT), trend in recurrence point density (TND), and laminarity/trapping time (LAM/TT)^4,5^. The optimal time delay and embedding dimensions were estimated from a mutual information and false nearest neighbors (FNN) functions for the red cluster are shown in B and C, respectively. The time-delay embedding was used to recontract *unthresholded* and *thresholded* (€=0.3) recurrence plots (RP).

##### 2.4.9. Principal component (PCA) analysis

was used to determine the direction of the maximum variances within SAN Ca2+ signals of the four color-coded regions during three critical phases of the global Ca2+ impulse (CaT) formation. The following markers were used as reference points for PCA in the regional clusters: the ignition phase (where the second derivative of the global CaT, d²Ca/dt², transitions from zero to its maximum), the fast phase (where d²Ca/dt² reaches its peak and progresses to the CaT maximum), and the decay phase (where the CaT returns to baseline). Four principal components were extracted, where PC1 and PC2 accounted for the most variances in the Ca2+ dynamics.

##### 2.4.10. Lorenz attractor analysis

was applied to quantify the degree of chaos in the system via estimation of the Lyapunov exponent of deterministic chaos (λ = 0.146), a non-chaotic system (λ ≈ 0.0001). Instead of a butterfly’s height, we are tracking the amplitude of the Ca2+ signal. Plotting the current amplitude value against its previous values reveals the patterns of Ca2+ signals change over time that are not usually obvious from the original time series.

To estimate λ in the Ca2+ signal, we employed a two-dimensional phase space reconstruction technique with lag = 1, minimum time separation = 4, tau = 1, and minimum neighbors = 10. The unit lag and tau ensured high temporal resolution, allowing analysis of the system’s state at consecutive time points without skipping any data. The minimum time separation of four helped mitigate the effects of short-term correlations (by ensuring that, we compare points that are at least four time steps apart). The criterion of 10 minimum neighbors for divergence calculations enhanced the statistical reliability of our Lyapunov exponent estimation by ensuring sufficient data points for analysis. These settings helped to optimize dynamics while maintaining statistical robustness with two-dimensional embedding.

##### 2.4.11. Fixed points stability analysis

An understanding of fixed points is to determine the behaviors of dynamic systems because these points represent the solution where the system could come to equilibrium. First, the fixed points of LCO of different ROIs were marked on 1^st^ derivative curves (when *f*(x*) = 0, velocity is not changing over time). Next, to assess stability, we evaluate the behavior of the system in the vicinity of the fixed points via linearization of the infinite polynomial function near a fixed point (Taylor series expansion).

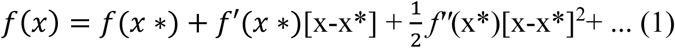

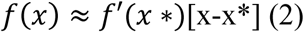

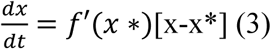

Integration of some steps of equation 3, generates equation 4, which represents the equation of motion near fixed points.

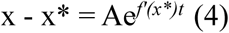

The sign of the 1^st^ derivative in the vicinity of fixed points, informs on a behavior of the system. A fixed point is stable (a ‘sink’) if perturbations lead the system back towards it, while it is unstable (a ‘source’) if perturbations push the system away. Therefore, if dCa/dt > 0, the solutions of the system will be unstable at this fixed point because the distance from the fixed point will exponentially increase; if dCa/dt < 0, it is a stable fixed point since the distance from the fixed point decays to zero. The sensitivity of the system can be significantly increased during the transition in the phase plane, where curves are near unstable fixed points.

##### 2.4.12. Empirical Mode Decomposition (EMD)

The 2.7-second time series recording of Ca2+ signals was analyzed using Empirical Mode Decomposition (EMD) ^6^. EMD is an adaptive decomposition technique used to extract frequencies from nonstationary signals. Compared to Fourier transform, EMD takes into account the non-linearity of the signal by iteratively sifting the original signal to extract oscillatory components known as Intrinsic Mode Functions (IMFs). The residual after the sift represents a non-oscillatory trend component. IMFs are characterized by several features, including local symmetry around zero and containing the same number of extrema as zero-crossings (or differing by no more than 1).

##### 2.4.13. Hilbert-Huang (HHT) transform analysis

that does not require signal stationarity was applied to compute the instantaneous phase, frequency, and amplitude of each IMF and quantify the power within a signal distributed across frequencies ^7^.

##### 2.4.14. Modified modified Short-Time Fourier Transform (STFT-FD)

In contrast to standard SFFT in which the window size is fixed in the time domain, STFT-FD applies a window that varies in size based on frequency, fixing the window size in the frequency domain ^8^. This technique provides better resolution at different frequency ranges within the signals, especially when analyzing signals with rapidly changing frequencies.

##### 2.4.15. A small-world network analysis

a powerful tool from network science was applied to investigate Ca2+ dynamics. To enhance the signal quality and separate calcium activity from noise in raw imaging, data series of preprocessing steps was applied. The penalized matrix decomposition (PMD) method was first applied, which decomposes the data into low-rank signal and noise components. Specifically, the patch-wise PMD (TV, TF) algorithm was used, applying total variation (TV) regularization spatially and trend filtering (TF) temporally. This step effectively denoised and compressed the data, improving the signal-to-noise ratio. Following the PMD, we applied a simple threshold to the data to further isolate signal from noise. We then used a median filter to smooth the thresholded data, which helped eliminate any remaining salt-and-pepper noise that might have persisted after the PMD and thresholding steps. The combination of PMD, thresholding, and median filtering significantly enhanced the visibility of calcium activity patterns and cellular structures, as evident in the improved clarity and contrast of the processed images in Supplementary Method Fig. 1 and Movie S2.

For each rolling time window during the 2.7 sec time series of SAN Ca2+ recording, weighted graphs based on mutual information matrices (Fig. S1B) were constructed (edges with top values above 2.0 were thresholded; and finally clustering coefficients and path lengths metrics were computed (Fig. S1A,C). These metrics were then compared to random graph models to derive the small-worldness metric σ, is the ratio of the average clustering coefficient and average shortest path length. For each time window, weighted graphs based on mutual information matrices were constructed; edges with top values above 2.0 were thresholded; and clustering coefficients and path lengths metrics were computed. These metrics were then compared to random graph models to derive the small-worldness metric σ. By calculating small-worldness (σ) over sliding windows throughout the cardiac cycle, network topology was constructed. Our analysis combined mutual information calculations and theoretical graph measures to analyze the functional connectivity and network property of the integrated (global) Ca2+ signal in the SAN (Fig. Method S1A).

##### 2.4. 16. Recurrence Quantification Analysis (RQA)

RQA detects hidden patterns and recurrences in a time series without assuming signal stationarity, making it particularly useful for short time series and discontinuous data. Unlike traditional timefrequency analysis, RQA reveals system behavior beyond frequency content, capturing aspects such as similarities, stability, determinism, noise and transitions between states. To perform RQA a onedimensional time-series data initially projected into a multidimensional state space, representing the system’s state at a given time, prior to transforming it into a recurrence plot (RP) (unthresholded and thresholded) using time-delay embedding (MATLAB Toolbox^4,5^). The RP visually represents the results of the RQA, i.e., quantification of patterns through specific metrics such as recurrence rate (RR), determinism (DET), longest diagonal line length (LMAX), entropy (ENT), trend in recurrence point density (TND), and laminarity/trapping time (LAM/TT)^4,5^.

##### 2.4. 17. Cross-recurrence Quantification Analysis (CRQA)

To compare interrelationship between different functional clusters, we performed a cross-recurrence quantification analysis (CRQA).^9^ In contrast to standard correlation methods, CRQA provides a comprehensive evaluation of the moments when two signals exhibit recurrence, capturing synchronization patterns over time, rather than merely assessing overall similarities or dissimilarities. Since CRQA directly compares signals point by point, it can identify leading-following relationship during interaction of two signals. MATLAB toolbox^4,5^ was used to perform this type of analysis.

###### Atrial cell action potential

Because action potential (AP) characteristics differ in the SAN pacemaker cells among regions of the SAN ^1^, to deduce the timing of SAN action potential (AP), we placed reference electrode in atria near the crista terminalis (CT). To record the transmembrane potential (AP), we used sharp microelectrodes (40–100MΩ) fabricated from aluminosilicate glass capillaries (1.5mm OD 0.86mm ID) with a horizontal pipette puller (Sutter Instruments, CA, USA), and back-filled with 3M KCl. Each microelectrode was inserted from the endocardial side of the tissue, and APs were recorded with a high-impedance amplifier (A-M Systems, USA) with a virtual bridge circuit for current injection. The electrical signal was digitized with Digidata 1440 and analyzed with PCLAMP10 software (both from Molecular Dynamics, USA).

## Supplementary Movies

**Supplementary Video S1.** From Bychkov et al., JACC: Clinical Electrophysiology (2020) with permission.

**Supplementary Video S2.** A novel filtering process of Ca^2+^ events, included trend filter denoising (based on spatially-localized-penalized matrix decomposition used on 2D neuronal dendrite activity), thresholding and median filter were applied to raw Ca^2+^ images to enhance the signal quality and separate Ca^2+^ activity from noise in small-world analyses.

**Supplementary Video S3.** 3D diagrams of the global CaT (F/F_0_) acceleration, velocity, and time across 19 impulses as shown in supplementary **Fig. 8C**.

## Supplementary Figures

**Table S1.** Mean Intervals Global CaT and atrial AP.

| ID | CaT | atrial AP |
| --- | --- | --- |
| SAN 1 | 145.10 | 145.30 |
| SAN2 | 185.84 | 184.70 |
| SAN 3 | 237.67 | 236.90 |
| SAN 4 | 232.00 | 232.00 |
| **mean** | **200.15** | **199.73** |
| **SD** | **43.42** | **43.25** |

**Fig. S1.**
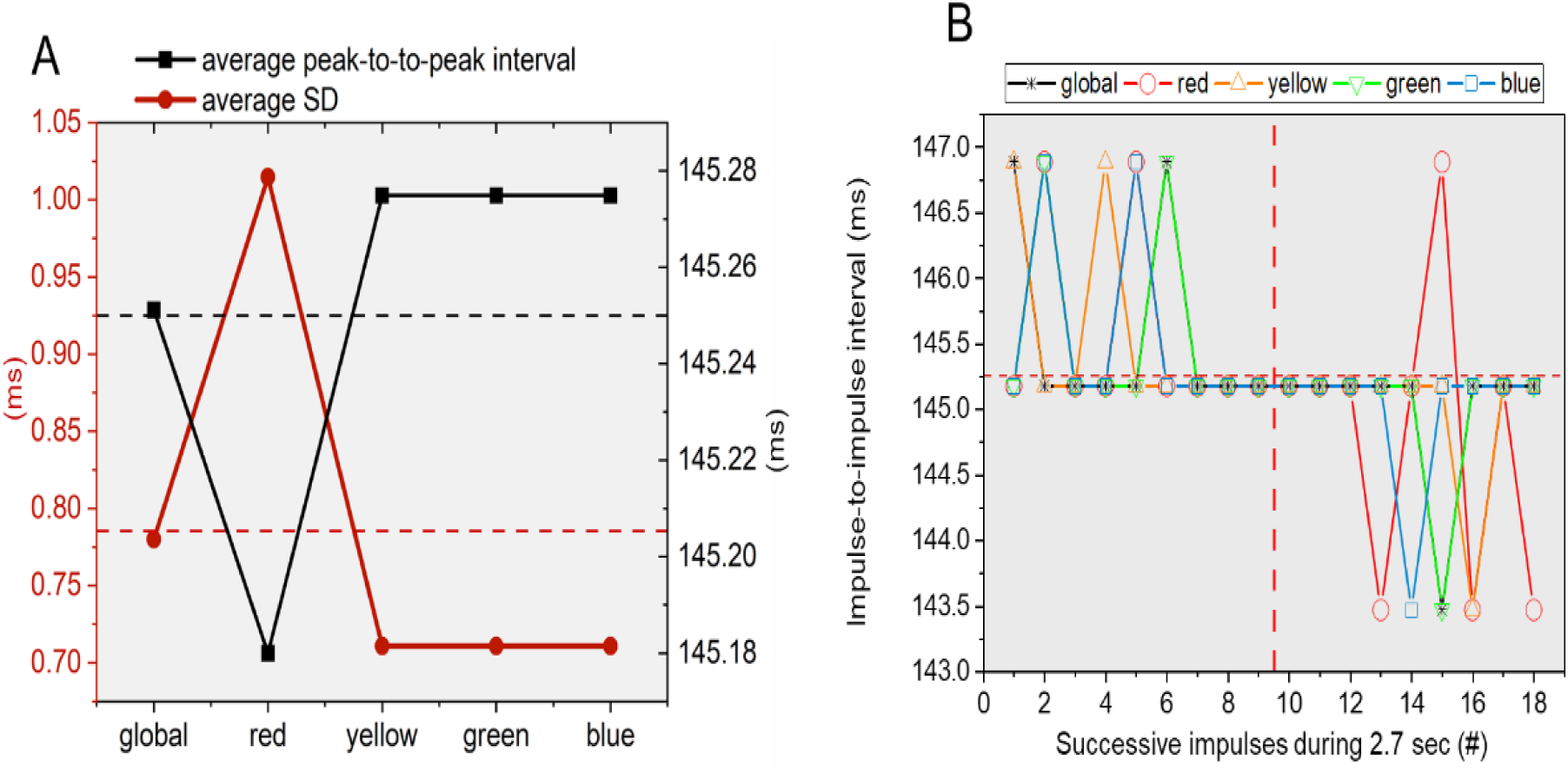
Cycle to cycle variability. (A) Average (right axis) and standard deviation (SD, left axis of the global CaT and each of four color-coded functional cluster. (B) Average peak-to-peak interval variability of the Ca^2+^ dynamic within each of four color-coded clusters and global CaT during 2.7-second recordings.

**Fig. S2.**
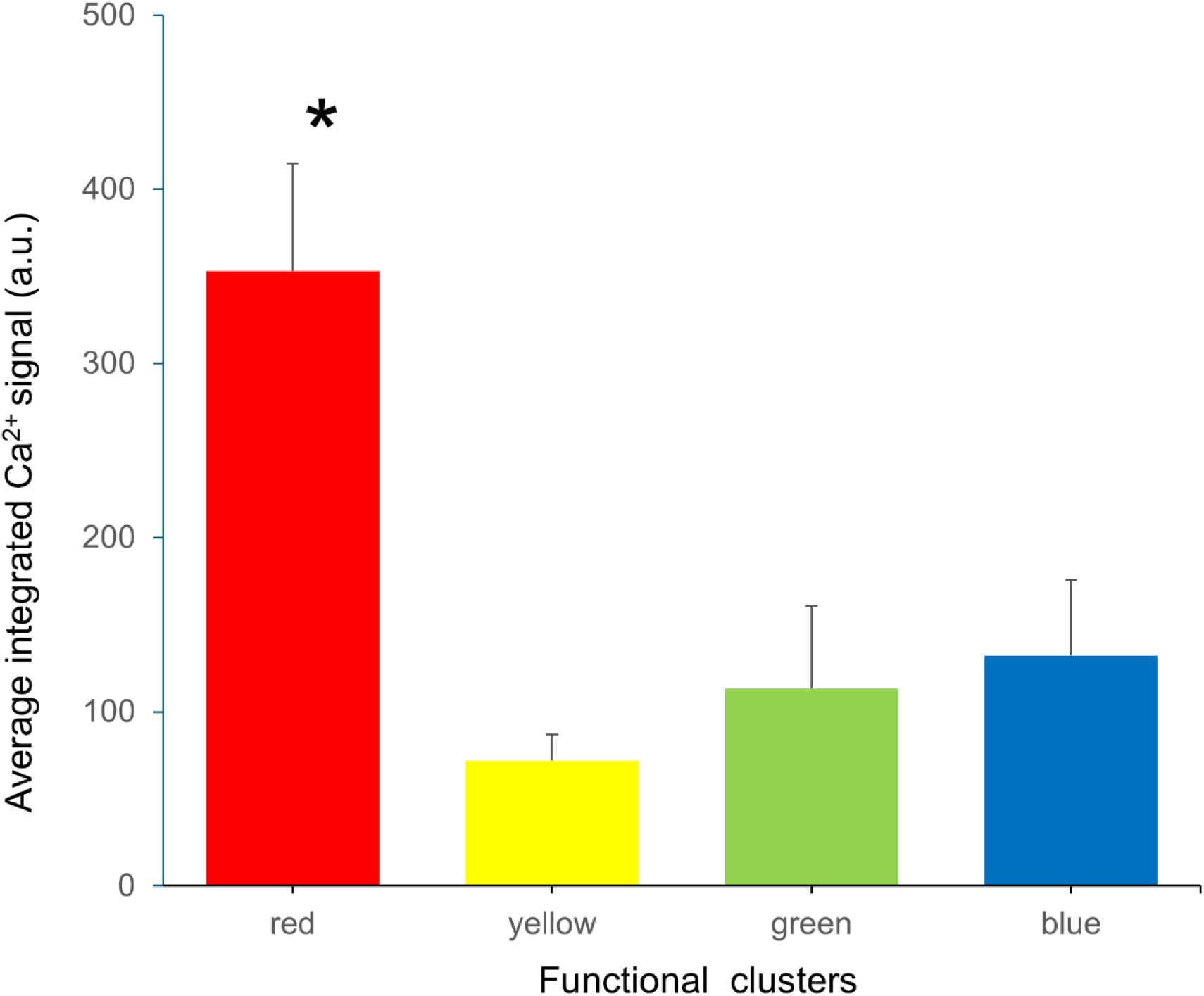
Averaged integrated Ca^2+^ signals (calculated in three cycles prior to initiation of the foot of the global CaT). ANOVA Tukey test p<0.05 vs yellow, green and blue average integrated Ca2+ signal.

### Frequency components analysis of the SAN global CaT

We quantified frequency domain characteristics of Ca^2+^ dynamics of the global CaT (**Fig**. S4) and of the each color-coded cluster (**Fig**. S4) throughout the entire 2.7-sec time-series recording by empirical mode decomposition (EMD) and Hilbert-Huang Transforms (HHT)^6^ (see methods)

**Fig. S3.**
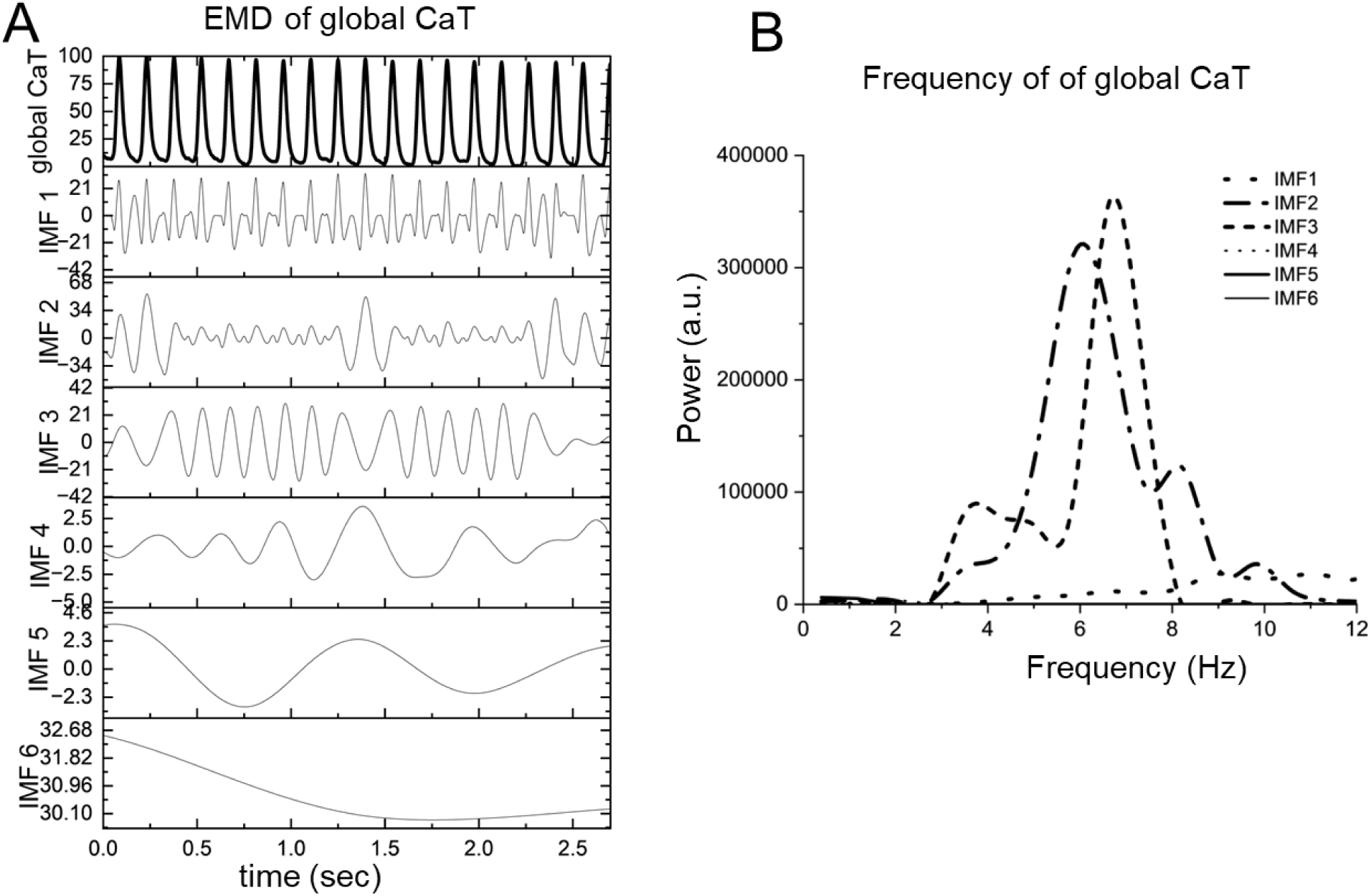
Frequency components of total energy expenditure throughout SAN tissue of the global CaTs. CaTs (Panel A top tracing) were decomposed into empirical mode functions (IMF, Panel A lower traces) by empirical mode decomposition (EMD) from which *instantaneous* frequency ranges were extracted by Hilbert-Huang Transforms (HHT)^6^. (B) The instantaneous frequencies corresponding to the IMFs in Panel A.

**Fig. S4.**
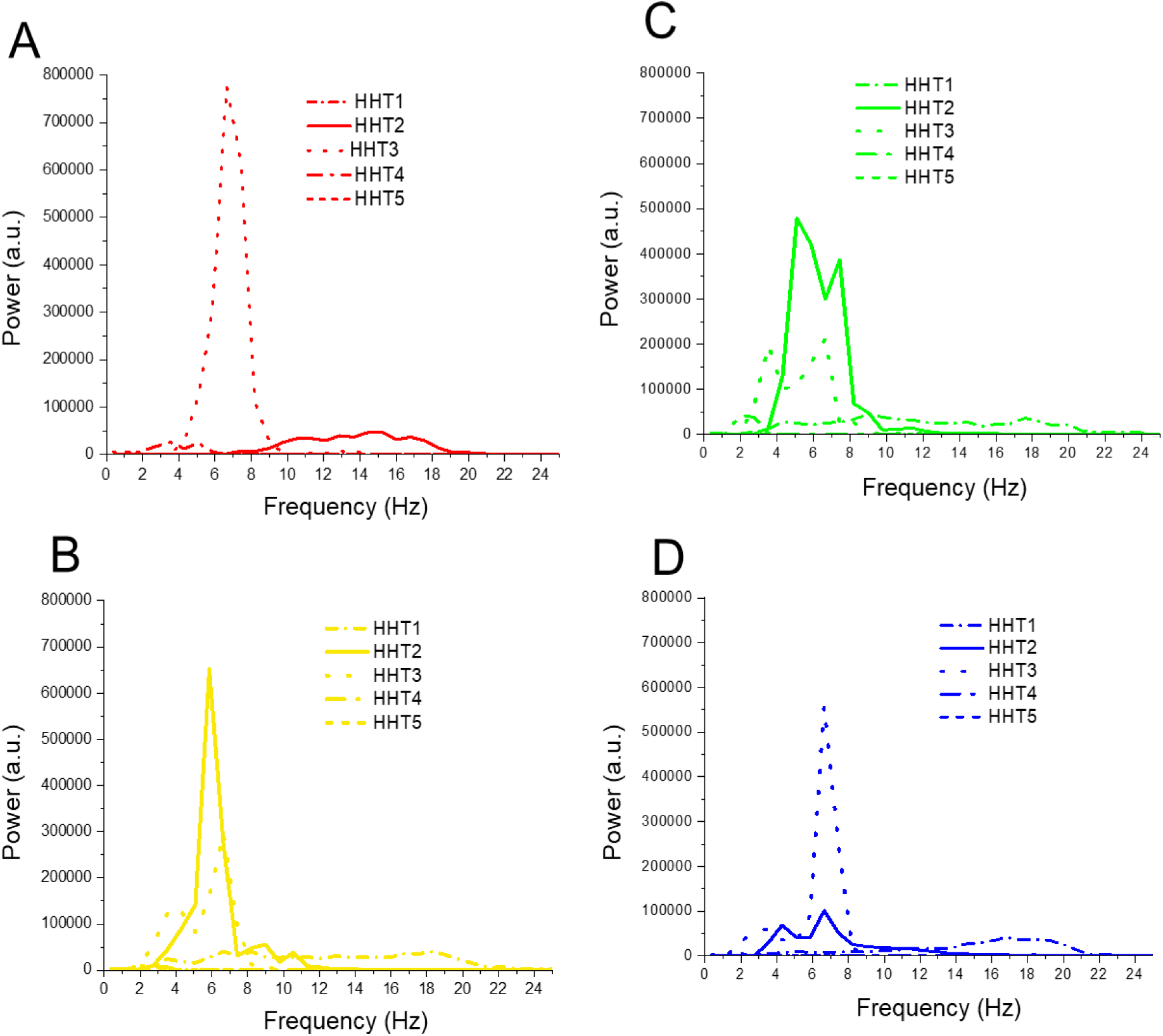
Frequency components of regional clusters. Complete frequency ranges for all IMFs of the four color-coded regional Ca2+ signals (red, yellow, green and blue) in Fig. 3A during the entire 2.7 sec time series recordings.

The series of Ca^2+^ impulses were decomposed into several empirical mode functions (Fig. S3, IMF, Panel A lower traces) from which instantaneous frequency ranges were extracted (Panel B). **Panel B** shows the power (amplitude) distribution across instantaneous frequencies within the four color-coded clusters (**Fig. S3**).

Figure S4 shows the complete frequency ranges for all IMFs of the four color-coded regional Ca2+ signals (red, yellow, green and blue) in Fig. 3A during the entire 2.7 sec time series recordings (see main text of Fig. 3 for details).

### Principal Component Analysis (PCA)

We performed principal component analysis (PCA) on biological noise every 10 points (approximately 17.3 ms) across the initial 190 ms of the Ca^2+^ time-series recording to determine how the Ca^2+^ dynamics within regional clusters collectively contributed to the occurrence of a global Ca^2+^ transient (CaT). A total of four principal components (PCs) were extracted from the analysis. The first two principal components, PC1 and PC2, accounted for over 90% of the total variance in the Ca^2+^ dynamics within the entire first global CaT. Therefore, PC3 and PC4 were excluded from these results. During the initial 0-17ms and 18-36ms of the recording **(Fig. S5, Panels B and C),** and *prior to* the initiation of the foot (pre-ignition) of the global CaT, PC1 and PC2 accounted for approximately 96% and 61% of the variance within the global Ca^2+^ dynamic. On average during this time, the red cluster contributed 78.5% to PC1, while the blue cluster contributed ∼19% to PC2 (P**anels B-C**).

**Fig. S5.**
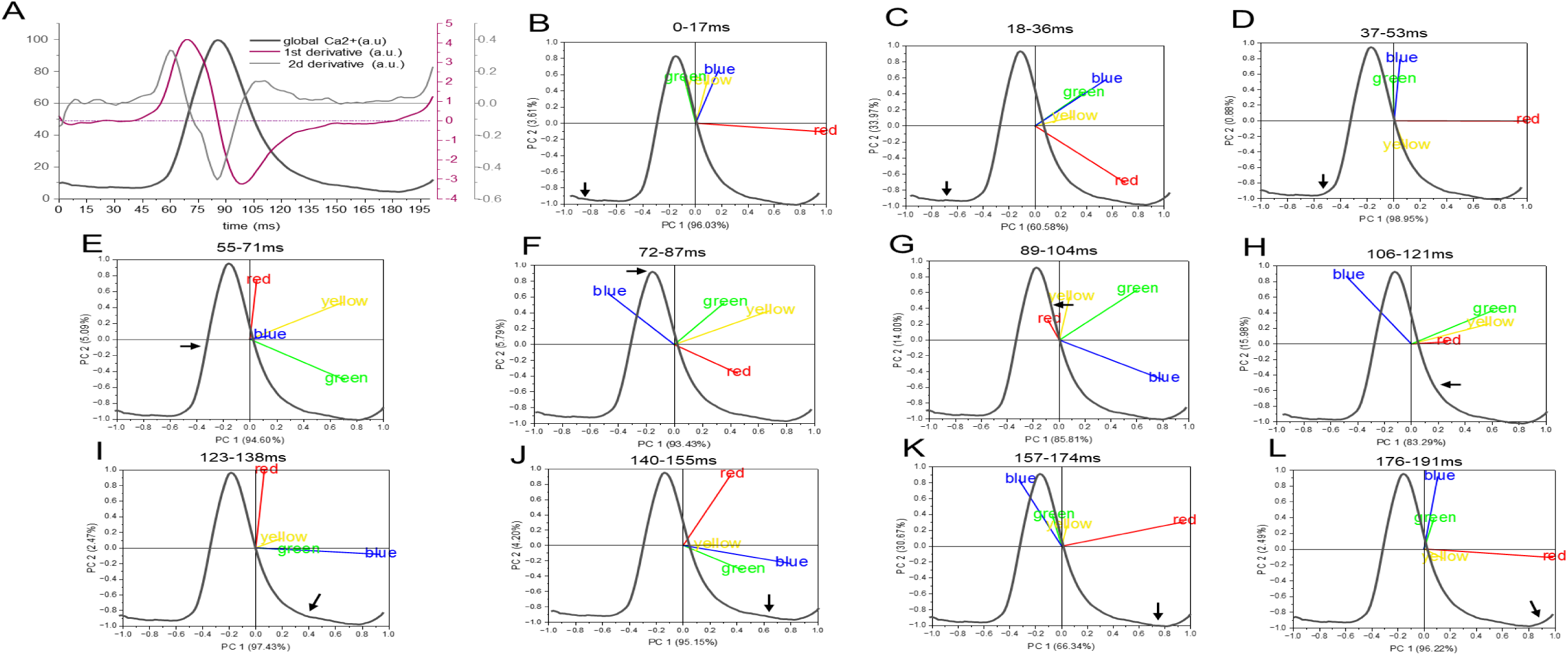
Principal component analysis (PCA) of the Ca^2+^ dynamics of the four color-coded SAN clusters during the 1st impulse formation (as depicted in Panel A) during 2.7 ms time series Ca^2+^ recording. (Panel A) The normalized waveform of the first global impulse (CaT) and its 1^st^ and 2^nd^ derivatives. (Panels B-L) Four principal components were extracted from each time window segment of 10 points (∼17.3 ms) across 190 ms. Since the 1^st^ two principal components accounted for most (>90 %) of the total variance within the Ca^2+^ dynamics are only shown and plotted as PC1 on the x-axis, and PC2 on the y-axis. Global CaT was superimposed on each PC plot for visualization of CaT dynamics at the time segment from which PCA were extracted. Arrows show the end time of each segment.

As the foot of global CaT (defined by the third derivative in **Fig. 1B**) began to form, the contribution of the red cluster Ca^2+^ dynamics to the PC1 increased to 99%, (**Panel D**). Following the end of the ignition phase, during the rapid upstroke of the global CaT (65-73 ms), the Ca^2+^ dynamics of the yellow and green clusters contributed about 95% to PC1of the global CaT dynamic, while the red cluster Ca^2+^ dynamic was responsible for 5% of the variations in PC2; the contribution of the blue cluster was minimal during this period (**Panel E**). This segment included the transition of the global CaT from acceleration (45-65ms) to maximum velocity (**Fig**. **1B**, ∼73 ms). The contribution of the Ca^2+^ dynamic, the yellow cluster, dominated PC1 until the global CaT reached its peak value at approximately 85 ms after its onset (**Panel F**). Immediately following the peak of the global CaT (**Fig. 1B**, ∼85 ms), the Ca^2+^ dynamic of the blue cluster made the largest contribution to PC1 (86%), while the green cluster accounted for 14% of PC2 **(Fig. S5G).** This trend reversed in the following segments during the mid and late decay phases of the global CaT (**Fig. S5H**). Near the termination of the first global CaT (∼140 ms), the Ca^2+^ dynamic blue cluster contributing 97% (**Fig. S3I**). **Panels J-L** illustrate the principal components from the end of the first global CaT to the beginning of the second CaT, highlighting the dominant contributions of the red and blue clusters in PC1 and PC2.

To assess how the Ca²⁺ dynamics of regional clusters contribute to the overall Ca²⁺ dynamic observed in the global calcium transient (CaT) in more detail, we conducted an analysis of the Principal Component *Spectra* (PCS) of the first derivative of Ca^2+^ signal (F/Fmin) of the 50 regions of interest (ROIs), grouped into four color-coded clusters throughout the entire time series Ca²⁺ recording (2.7 sec). Note that the ellipse representing the yellow cluster in **Fig S6**, overlaps substantially with that of the green cluster, and the green cluster overlaps with the blue, while the red ellipse connects all three clusters. Thus, the Ca²⁺ dynamics across *all* SAN functional clusters *collectively* influence the dynamic of the global SAN CaT (as illustrated in **Figs. S5 and S6**).

**Fig. S6.**
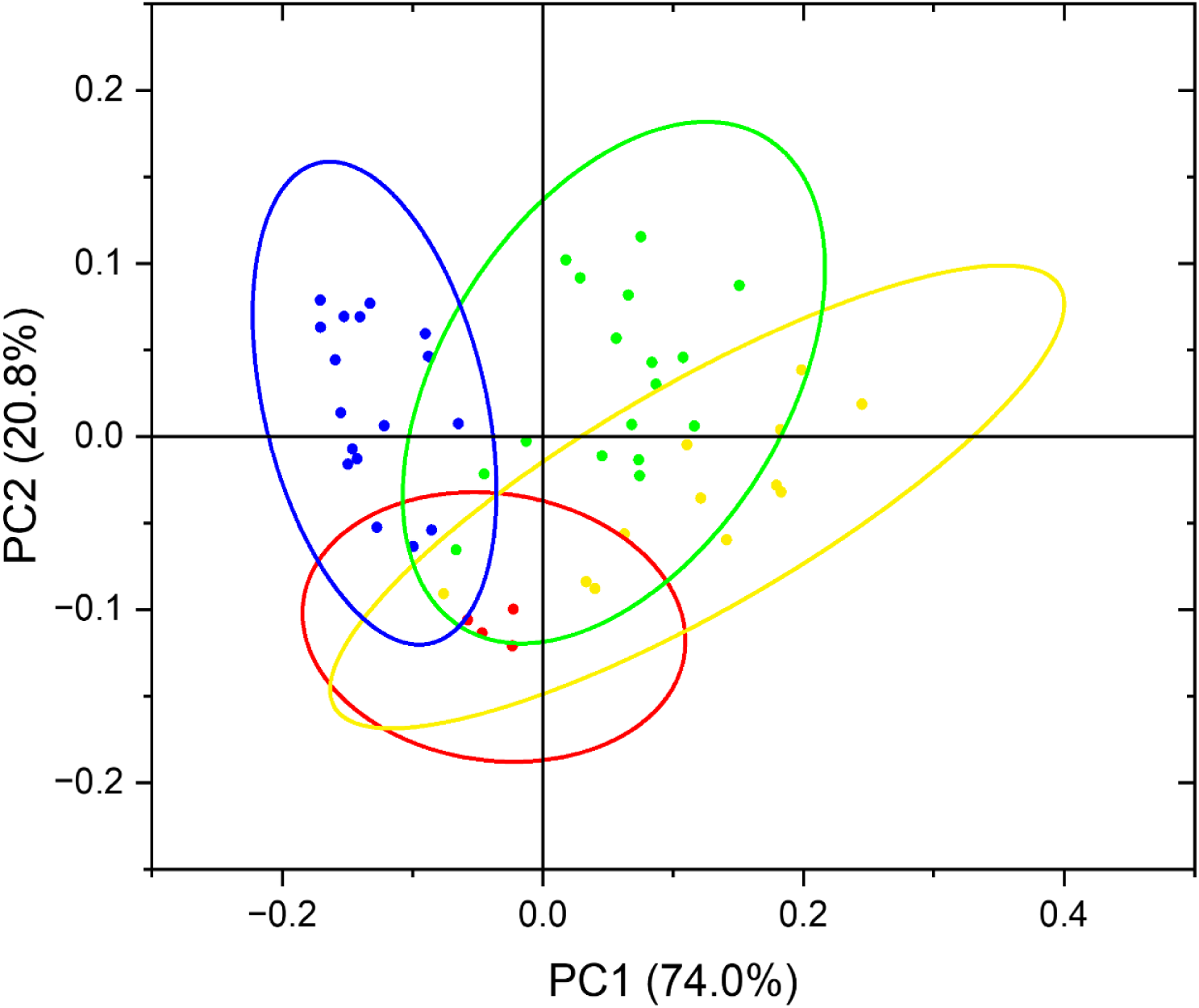
Principal component *spectra* (PCS) of the Ca2+ dynamic. Ca2+ signal (F/Fmin) (first derivative) of the 50 regions of interest (ROIs), **grouped** into four color-coded clusters throughout the entire time series Ca²⁺ recording (2.7 sec).

### Energy transition during the global SAN CaT

We performed functional data analysis (FDA)**^10^** the dynamic of energy transitions in the each colorcoded cluster throughout the time-series (Fig. 5 see main text), and in the global CaT (Figs. S7-S8). Fig. S7 shoiws energy transitions during the 1^st^ global CaT in the time-series (Fig. S8A top panel) and the ROI correlations during time segments (see below, Fig. S9) of the same impulse. Prior to the initiation of the global CaT (foot) and at the time when there was no or small correlation (Fig. S7E-F) among Ca^2+^ dynamic of any ROIs, energy transition was very low (**Fig. S7A**).

In contrast, during the foot of the global SAN CaT (∼50-65ms) when a large number of correlations among ROI Ca^2+^ dynamics emerged (**Panel G**), the energy expenditure was high (**Panels A, B** at above 0:0,). Similar pattern of phase plane dciagram of acceleration vs velocity were observed during the integrated (global) CaT of all 19 impulses are shown in **Fig. S8,** <u>Video S3.pptx</u>. However, there was a beat-to-beat variability in both, acceleartion and velocity.

**Fig. S7.**
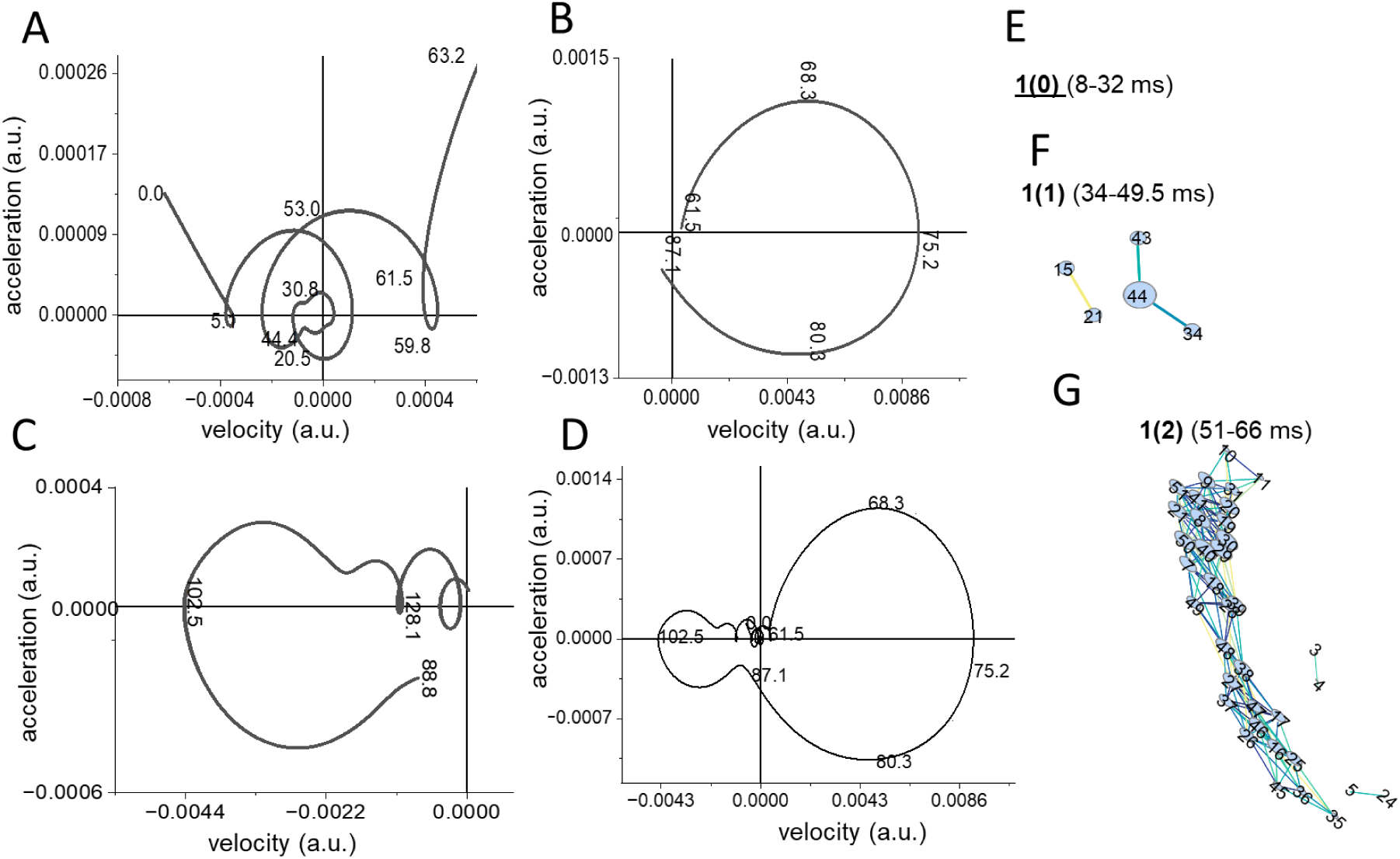
Energy transfer during global CaT. The relationship between acceleration and velocity of the Ca^2+^ dynamic of the global SAN CaTs and concurrent time-dependent correlations among ROI Ca^2+^ dynamics. FDA phase-plane diagrams of the global CaT acceleration (the 2^nd^ derivative) plotted against velocity (the 1^st^ derivative) during the 1^st^ global SAN CaT. The functional data analysis is segmented into three distinct sections based on the shifts along the x/y-axes, and the corresponding concurrent correlations of ROI Ca^2+^ dynamics are in Panels E-G . Panel A illustrates a phase-plane diagram between 0-63ms, Panel B between 63-87ms, and Panels C between 89-130ms of impulse formation., Panel D illustrates all sections.

**Fig. S8.**
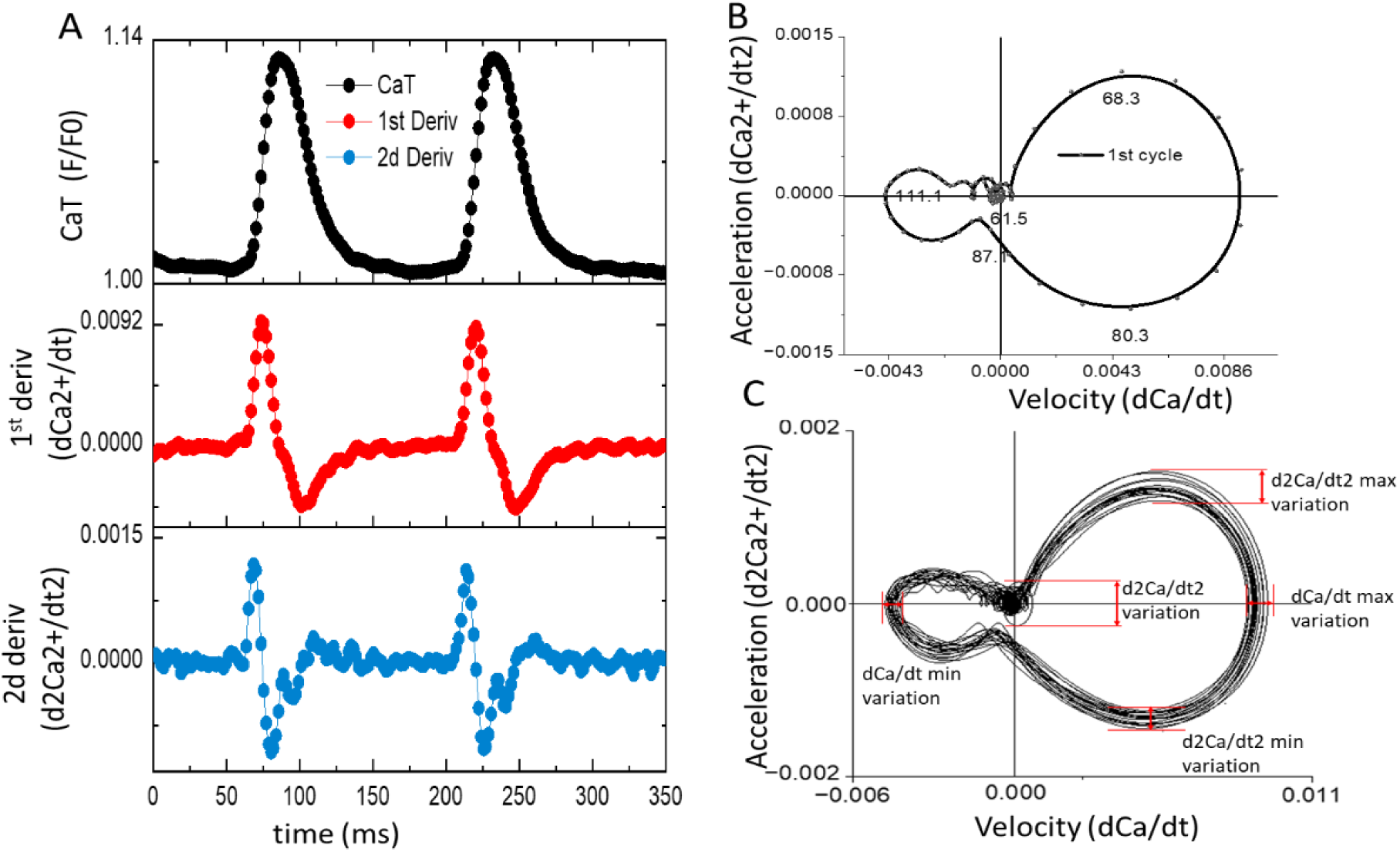
Energy transfer during global CaTs in all 19 impulses. See Fig. S7 for details and Video S3.

#### Pearson Correlation Coefficient Analysis

We employed Pearson correlation analysis to determine trans-spatial relationships of LCO dynamic among the 50 ROIs forming four color-coded SAN functional clusters as a function of rolling window (∼15ms) throughout the first 350 ms of the time-series Ca^2+^ recordings (Fig. S9). We focused only on correlations coefficient that were greater than 0.99 to construct the adjacent matrix.

The normalized amplitudes of Ca^2+^ signals within each of the 50 ROIs overlaid by the global SAN Ca^2+^ impulse are shown in **Fig. S9B** The color-coded map of the 50 ROIs clustering is displayed in **Panel A**. Amplitude correlation patterns among ROIs (**Panels C-K**) differentially emerged and decayed with time during the 350 ms time-period, during which two complete integrated (global) SAN CaT emerged (black trace in Panel B). Correlation patterns prior to and during the two impulses are shown side by side in **Panels C-J**. Blue and green numbers at the top of each time segment indicate the 1^st^ and 2d impulses.

Prior to the initiation of each impulse (see vertical lines marked as 1(0) and 2(0) in Panel B for time reference), ROI LCO amplitudes were completely *unclustered* (**Panel C**), suggesting stochastic behavior among LCO comprising 50 ROI. During the next time segment (see vertical marked as 1(1) and 2(1) in Panel B) two small clusters formed in 1(1), one of which included only 2 ROIs and the other included 3 ROIs; and in 2(1) only one cluster formed (Fig. S9D). These clusters in both 1(1) and 2(1) formed near the crista terminals (red color in the ROI map in A). In the following time segment (1(2) and 2(2)), Ca^2+^ signal amplitudes within a large number of ROIs formed one large homogeneous cluster, a few small clusters (**Fig. S9E**), and only 28% remained *unclustered* in 1(2). Note, that during this specific time segment (see vertical lines in Panel B for time reference), the amplitude of the integrated (global) SAN CaT slowly increased (**Panel B**).

During the next segments (**Panel B** marked as 1(3) and 2(3) for time reference), the clustering patterns again changed: two major clusters and a few small clusters were formed, with ∼ 20% ROIs remaining *unclustered* (**Fig. S9F**). During this segment, the amplitude of the integrated (global) SAN CaT rapidly increased near its peak (**Fig. S9B**). During the next time segments 1(4), 2(4), one large cluster encompassing a large number of ROIs, and a few small clusters were observed, and 20% and 6% of total ROIs were *unclustered* (**Fig. S9G**). During this segment, the integrated (global) SAN CaT began to decay (**Panel B,** black trace). In the following segments, 1(5), 2(5), large clusters of ROIs in the prior segment reorganized into two medium-sized correlation clusters a few very small clusters, and ∼20% remained uncorrelated (**Fig. S9H**). At this time, the integrated (global) SAN CaT had decayed rapidly to ∼50 % of its peak amplitude (**Fig. S9B**). In the following time segments 1(6), and 2(6), ROI Ca^2+^ signals became largely unsynchronized, and only one medium-sized cluster and three small clusters remaining, and ∼28% of ROIs were uncorrelated ; during this time, the integrated (global) SAN CaT decayed rapidly to ∼18% of its peak amplitude. The next time segments, 1(7) and 2(7) showed only a few small correlation clusters, and in 3(0) ROIs were completely uncorrelated again (**Fig. S9K**). Thus, during the 350 ms recording, there was an indication of memory-like behavior indicated by similar, but not exact, clustering patterns of ROIs in 1(0)-1(7) 2(0)-2(7): initially ROIs were uncorrelated, and with time a gradual shift to a higher level of correlation among ROIs occurred, followed by loss of correlation, eventually, again, with ROIs becoming completely uncorrelated.

Figure S10 shows the correlations among the kinetics of the average Ca^2+^ dynamics of the four color-coded SAN functional clusters and between them with the global SAN CaT during the entire 2.7-second time-series recording. The correlation between the yellow and green functional clusters was higher (r=0.90) than between the red and yellow (r=0.78) or red and green (r=0.53) functional clusters. The red and blue functional clusters negatively correlated (-0.091) with each other. There was some small positive correlation between the yellow, green, and blue functional clusters. The global CaT more strongly correlated with yellow (r=0.89) and green (r=0.98) functional clusters than with red (r=0.57) and blue (r=0.53) functional clusters (**Fig. S10**).

**Fig. S9.**
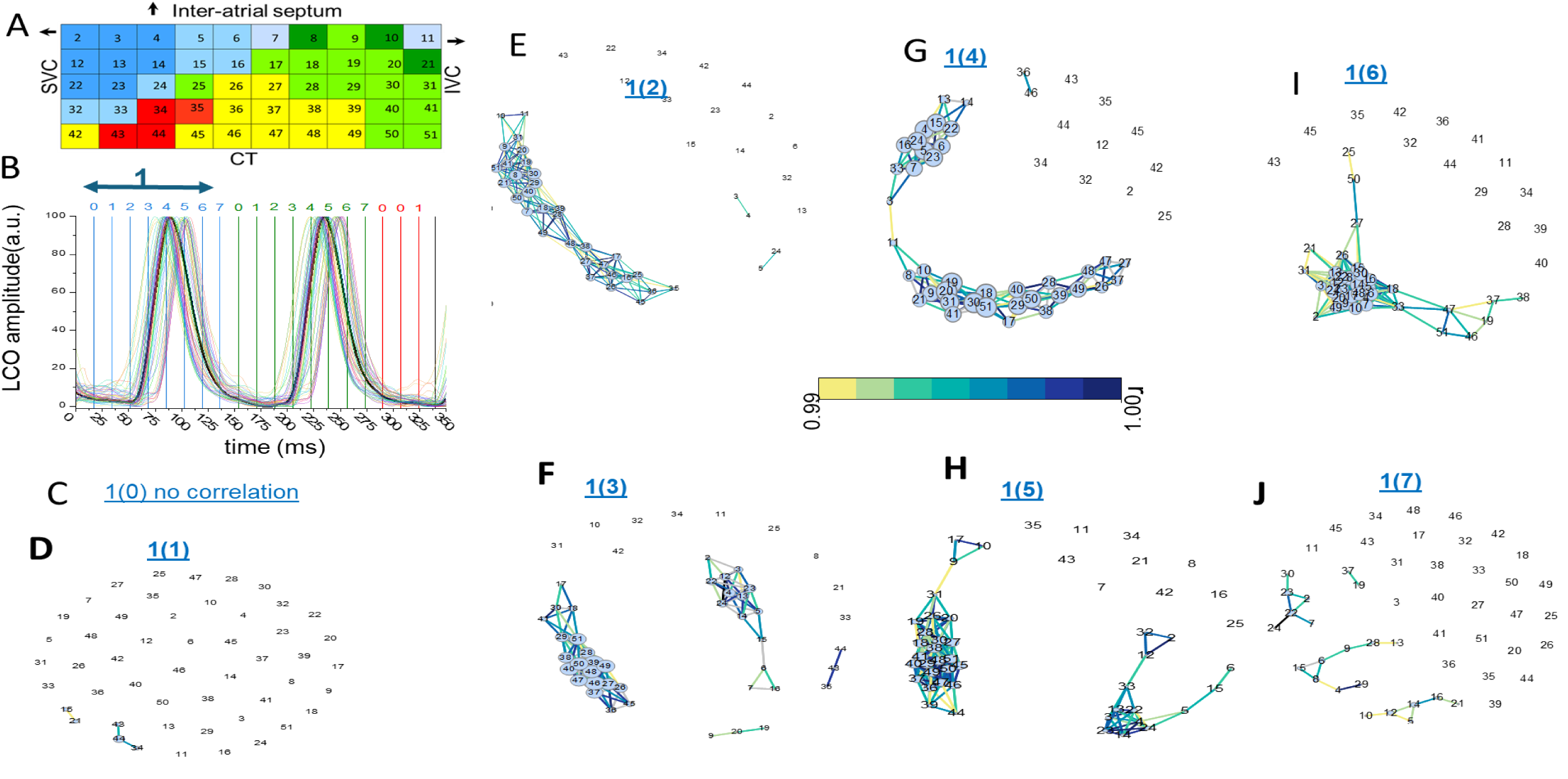
Spatio-temporal cluster correlations of SAN local Ca^2+^ oscillation (LCO) dynamics during the first impulse formation. (A) A color-coded map illustrating the spatial distributions of clusters of SAN functional regions of interest (ROIs). (B) The normalized (0-100%) LCO amplitudes of the Ca^2+^ dynamics of the 50 ROPanel A during the initial 350 ms of a time series Ca^2+^ recoding, during which global CaTs emerged. Each ∼17 ms segment during the 1st impulse) is referred to by numerical values 0-7. The normalized global Ca^2+^ signal waveform (CaT) is shown in black. (C-K) Pearson correlation analyses performed in a continuous rolling window (∼17 ms). Plots of network correlation coefficients (r ≥ 0.99) for ROI LCO dynamics during each 17ms window in each segment from Panel B are presented side-by-side in Panels C-K. The network for impulse 1 (Panel B) is indicated by blue circles, while the network for impulse 2 (Panel B) is represented by green circles. Color-coded (see heat map in the center of the figure) lines connecting ROIs within the network, reflect the strength of their correlations during a given window.

**Fig. S10.**
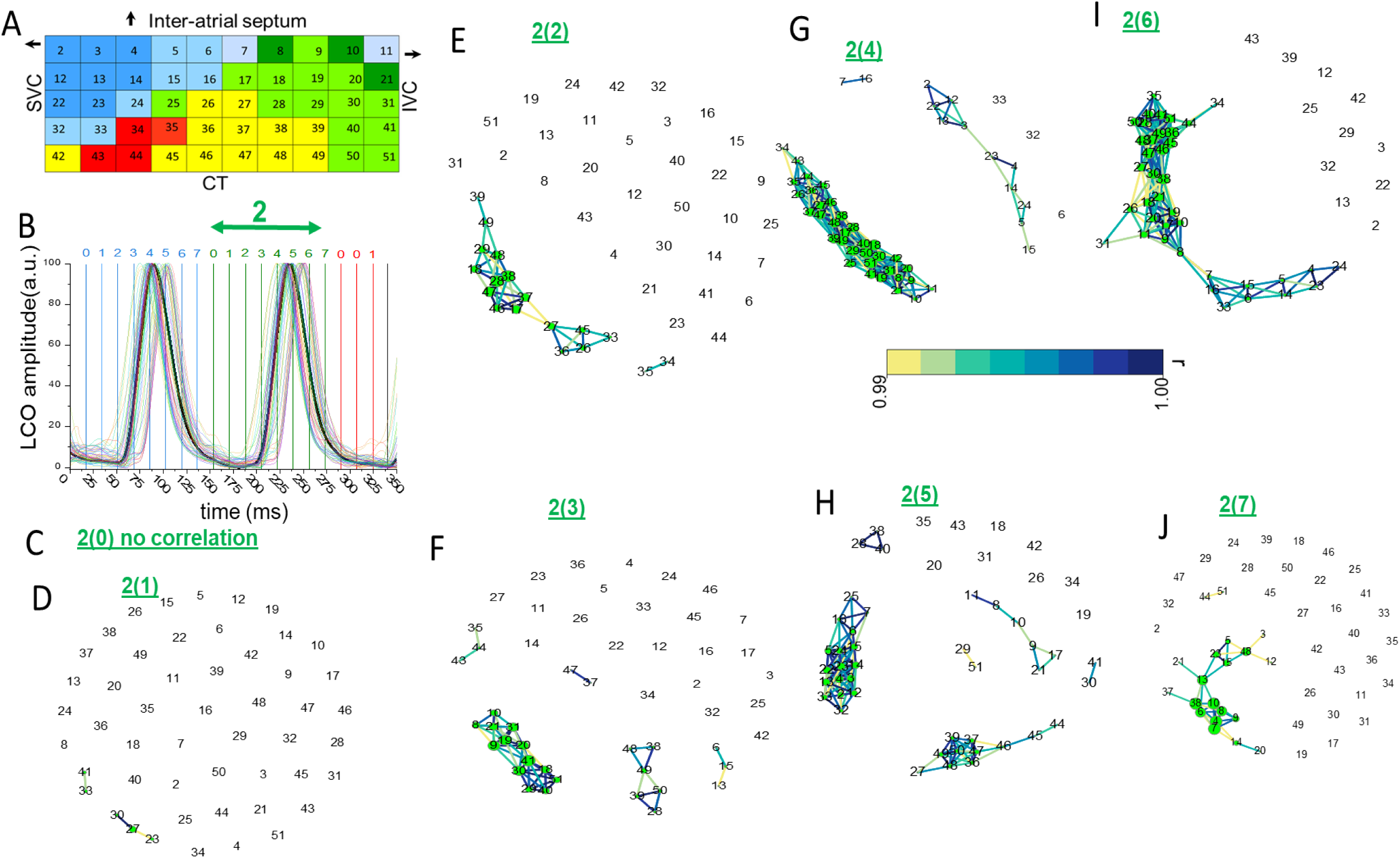
Spatio-temporal cluster correlations of SAN local Ca^2+^ oscillation (LCO) dynamics during the second impulse formation. The network for 2d impulse (Panel B) is represented by green circles. See details in **Fig. S9**.

**Fig. S11.**
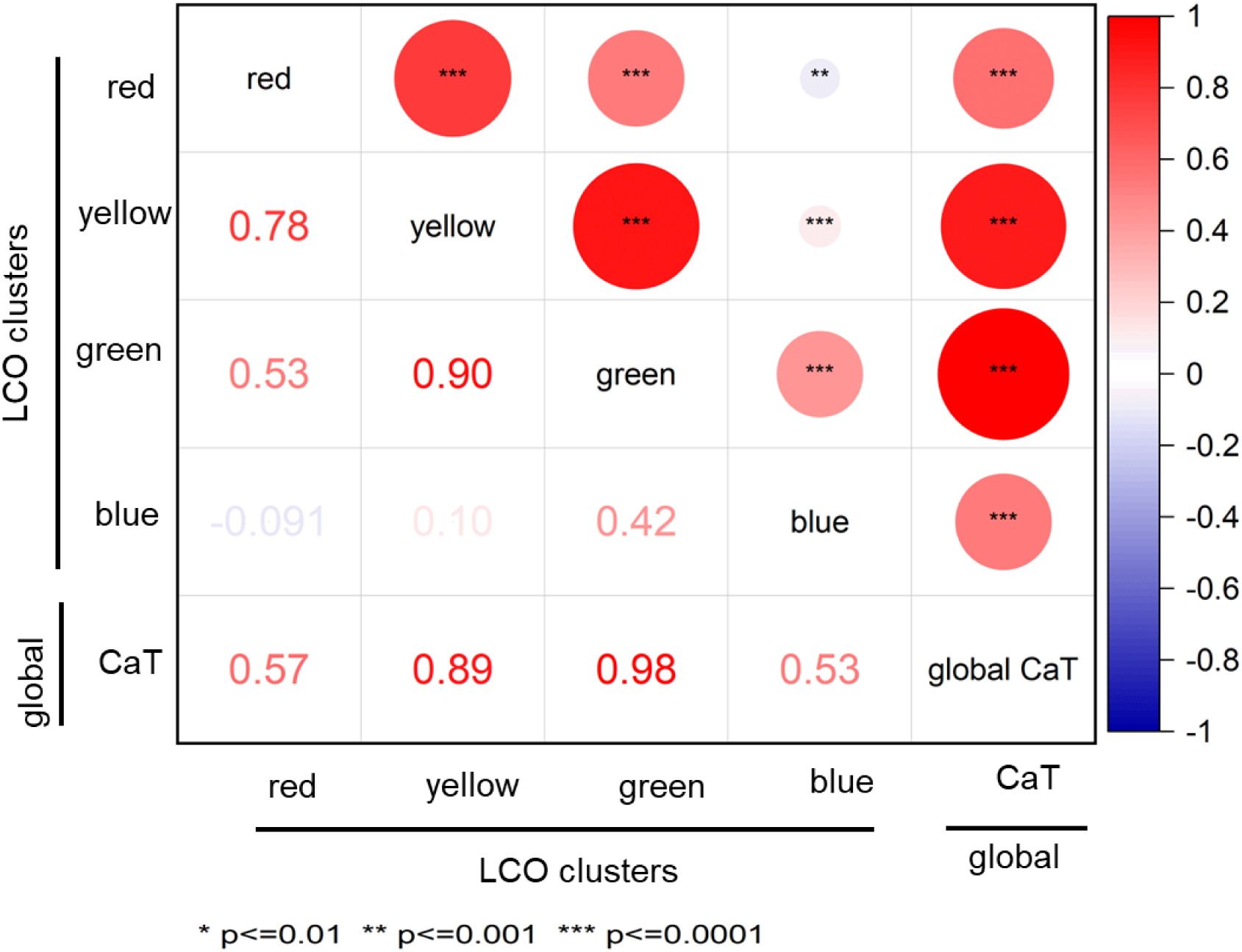
Correlation matrix of the kinetics of the average Ca^2+^ dynamic within four distinct SAN color-coded functional clusters, and global CaTs during an entire 2.7-second time-series recording. The upper triangular portion of the matrix highlights statistically significant correlations, while the lower triangular section depicts the magnitudes of the corresponding correlation coefficients. Positive correlations are represented by red circles; negative correlations are indicated by a blue circle. The size of each circle is proportional to the strength of the correlation, and the vertical color bar serves as a reference for the correlation coefficients.

**Fig. S12.**
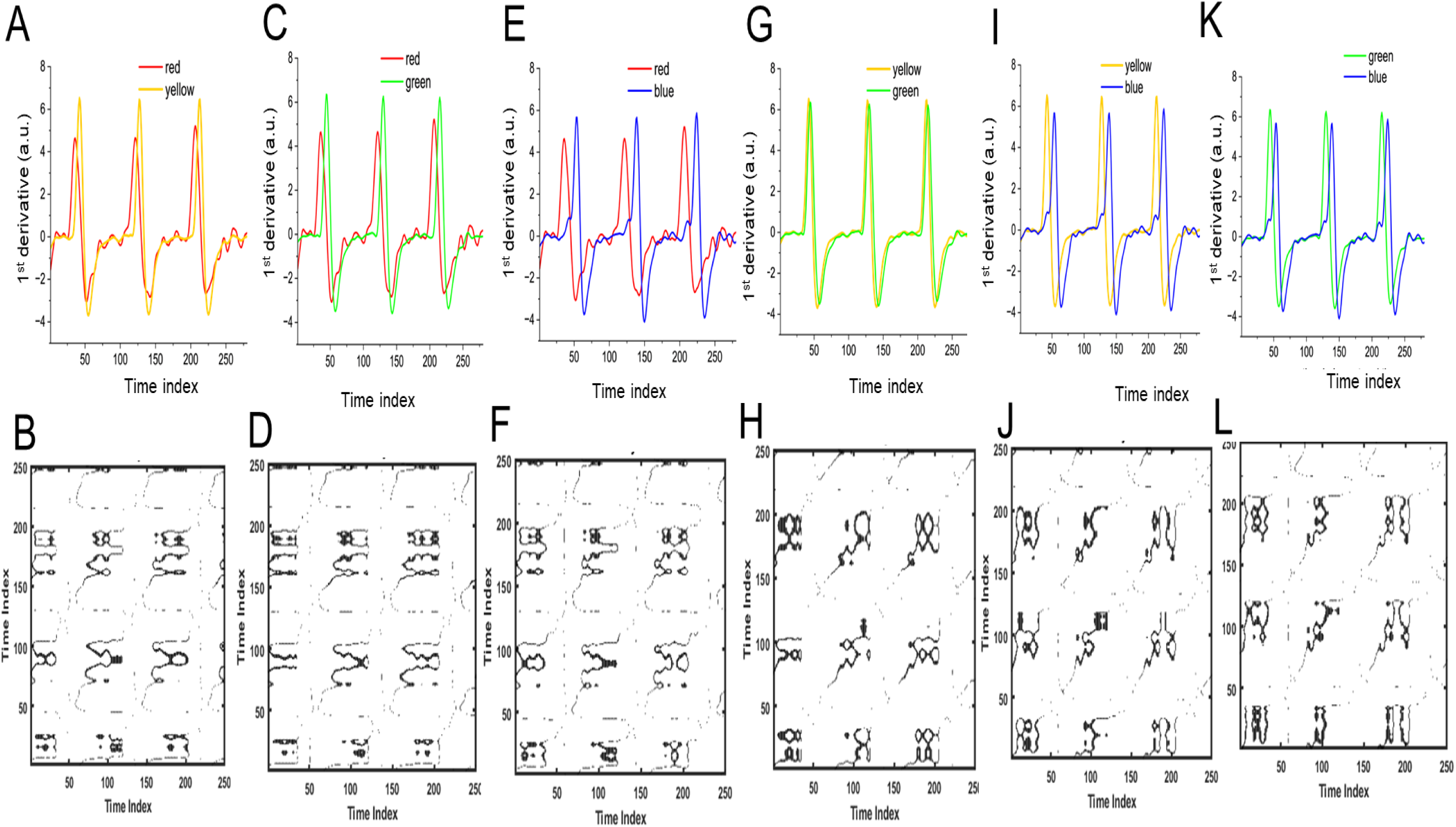
Cross-recurrence quantification analysis between four color coded clusters. Each of upper panel shows superimposed Ca^2+^ dynamics (rate of change) of two different clusters. Lower panels represent pairwise comparison its respective thresholded cross-recurrence plots (TCRPs) for all pairs of functional clusters. MATLAB toolbox^4,5^ was used to perform this type of analysis.

### Orbit plots analysis

We constructed orbit plots, which embed additional dimension of time-shifting segments (7ms) throughout 2.7 sec of Ca^2+^ dynamics^3,11^. Fig. S13 shows orbit plots constructed from the spatially averaged Ca^2+^ dynamics within the four color-coded clusters (**three cycles, Panel A**) as the unlagged and lagged (7ms) versions of itself during 500 ms (**Panel B,** three impulses), 180 ms (**Panel C**, one impulse) and 65 ms (**Panel D,** ‘pre-foot’ and ‘foot’ formation) of impulse generation, and Panel F illustrates the orbit plots for the 19 impulses of the global CaT depicted in Panel E.

**Fig. S13.**
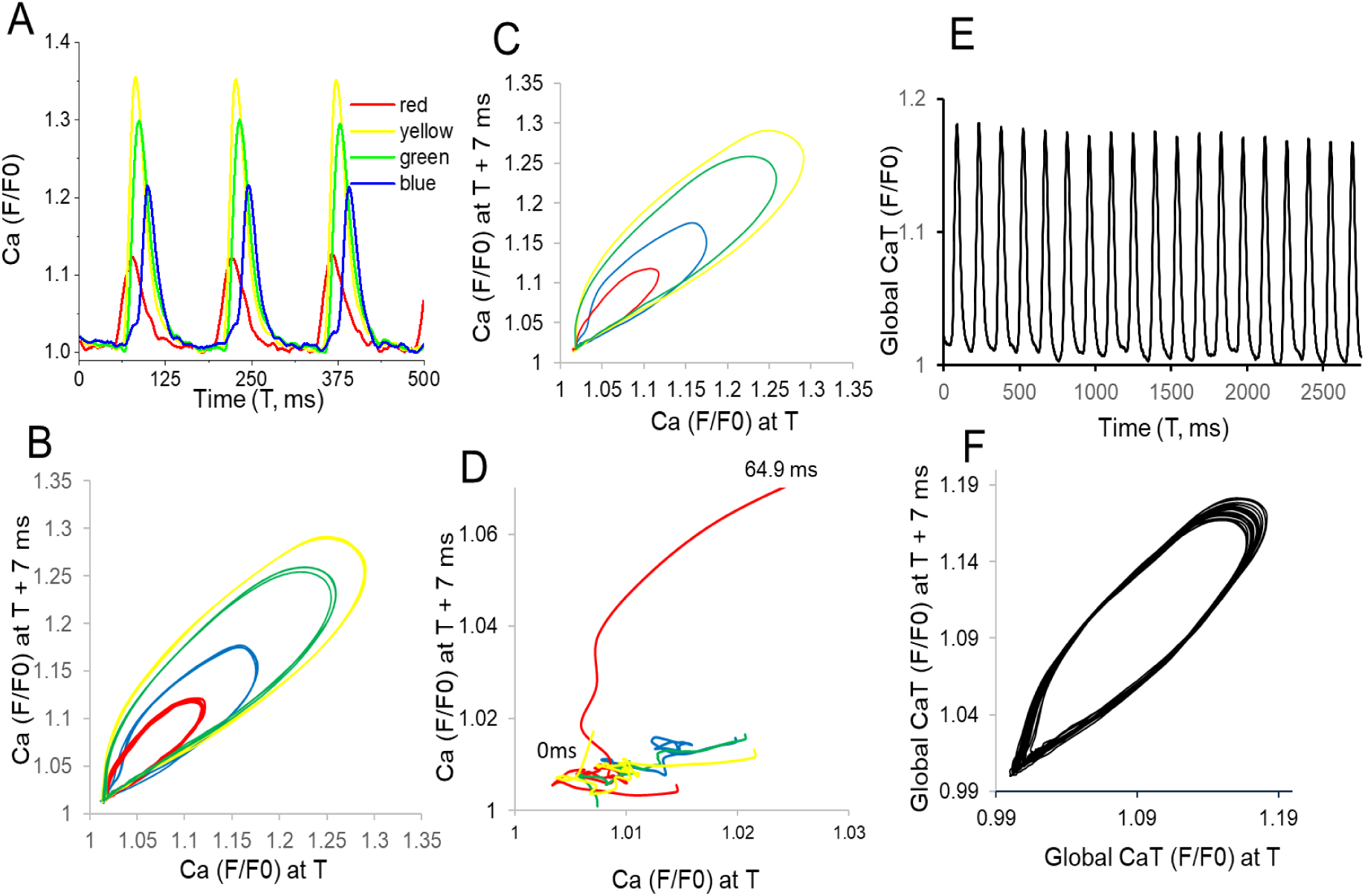
Visualization of the direction of rotation and time-delayed trajectories of reginal and global Ca^2+^ signals. (A) Depicts the spatially averaged amplitudes of the Ca^2+^ dynamics across four color-coded clusters during a period of 500 milliseconds, in which three global CaTs emerge. (B) The orbit plots generated from the Ca^2+^ dynamics in Panel A are displayed in both unlagged and lagged (+7ms) formats. (C) Orbit plots of Ca^2+^ dynamics during a shorter period (180 ms) during the time series in Panel A in which the first global CaT emerged. (D) A magnified view of the orbit plots in Panel C focusing on the initial (∼64.5 ms, pre-foot and foot) global CaT formation. (E) Global CaTs recorded during an entire longer 2.7 sec. time-series recording. (F) Orbit plots derived from the time series of the global CaTs in Panel E, presented in both unlagged and lagged (+7ms) formats.

Each of the color-coded Ca^2+^ dynamic trajectories begins with a small oscillation, as clearly seen in **Panel D,** and that a clockwise elliptical-like trajectory ended near to its starting point, but not exactly at always at the same starting point (**Panels B-C**). This difference in the initial condition creates variability of orbits within a given cluster, suggesting that SAN Ca^2+^ dynamic is not purely deterministic but partially stochastic. Example of orbit plots of individual ROIs from each of the four color-coded cluster during entire time-series showed some degree of impulse-to-impulse variability observed within each cluster (**Fig S10**). However, it appeared that orbit trajectory within the red cluster was greater than that of other clusters.

**Fig. S14.**
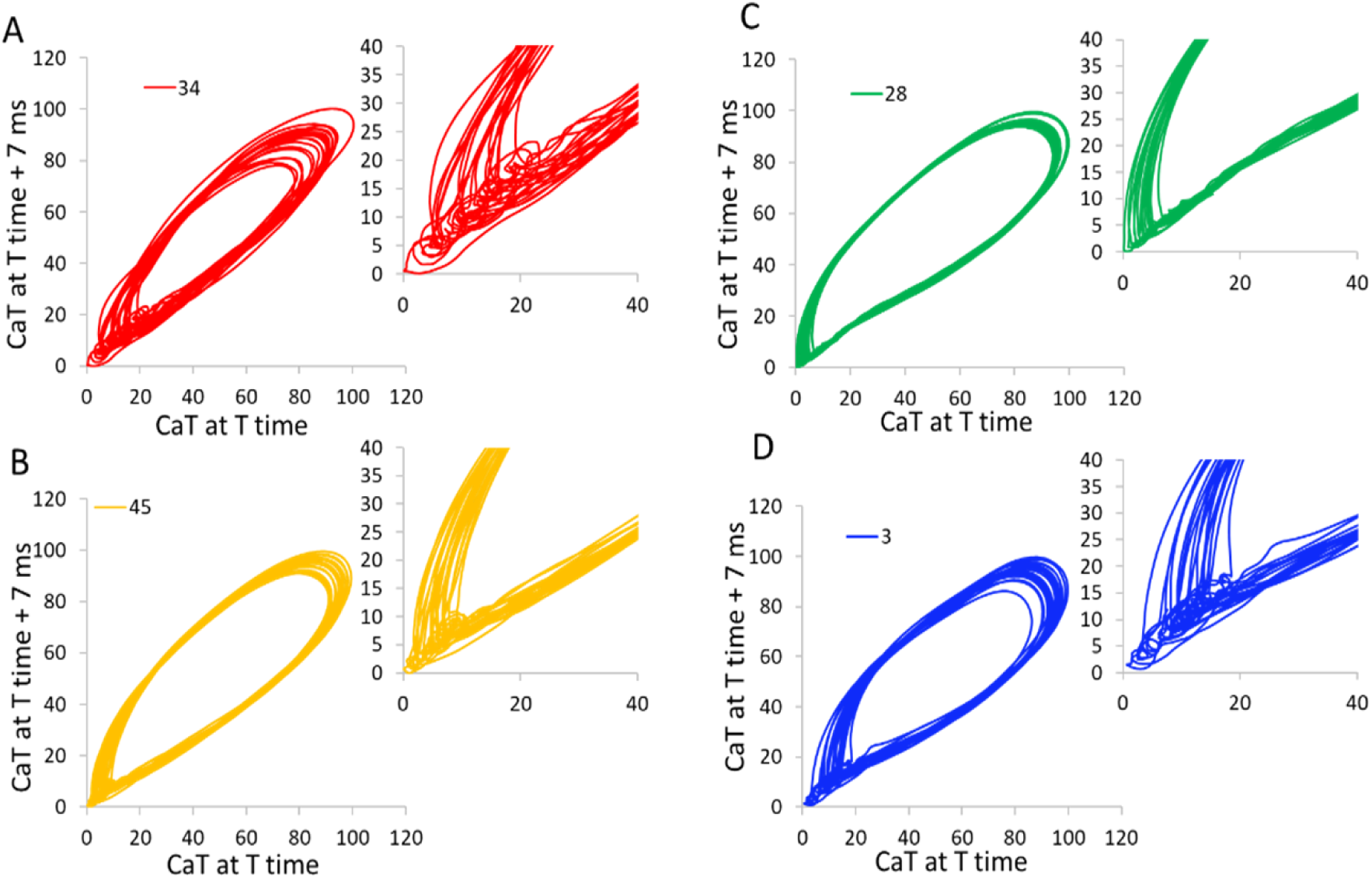
Examples of orbit plots for individual regions of interest (ROIs) from each of the four color-coded clusters.

### Lyapunov exponent (λ) analysis

We calculated the Lyapunov exponent (λ), a measure of how quickly nearby Ca^2+^ dynamic trajectories in the phase-space diverge over time to estimate the degrees of stochastic and deterministic behavior embedded within the 2.7 sec time-series recording. To estimate λ, we employed a two-dimensional phase space reconstruction technique optimized for capturing system dynamics, while maintaining statistical robustness with two-dimensional embedding (which affords visualization of the system’s trajectory in phase space, effectively capturing the essential dynamics of the Ca^2+^ signaling network (see Methods).Fig. **S15, Panel A** illustrates an example of the strongly deterministic (chaotic) behavior of a Lorenz attractor (λ = 0.146), **Panel C** illustrates the behavior of a stochastic sinusoidal wave (λ ≈ 0.0001, Panel C), and **Panel B** illustrates the behavior of the SAN global CaT (λ = 0.082). The positive Lyapunov λ of the SAN Ca^2+^ signaling network confirms its sensitivity to initial conditions, consistent with the nonlinear patterns and memory observed in the orbit plots (**Figs. S13-14**).

**Fig. S15.**
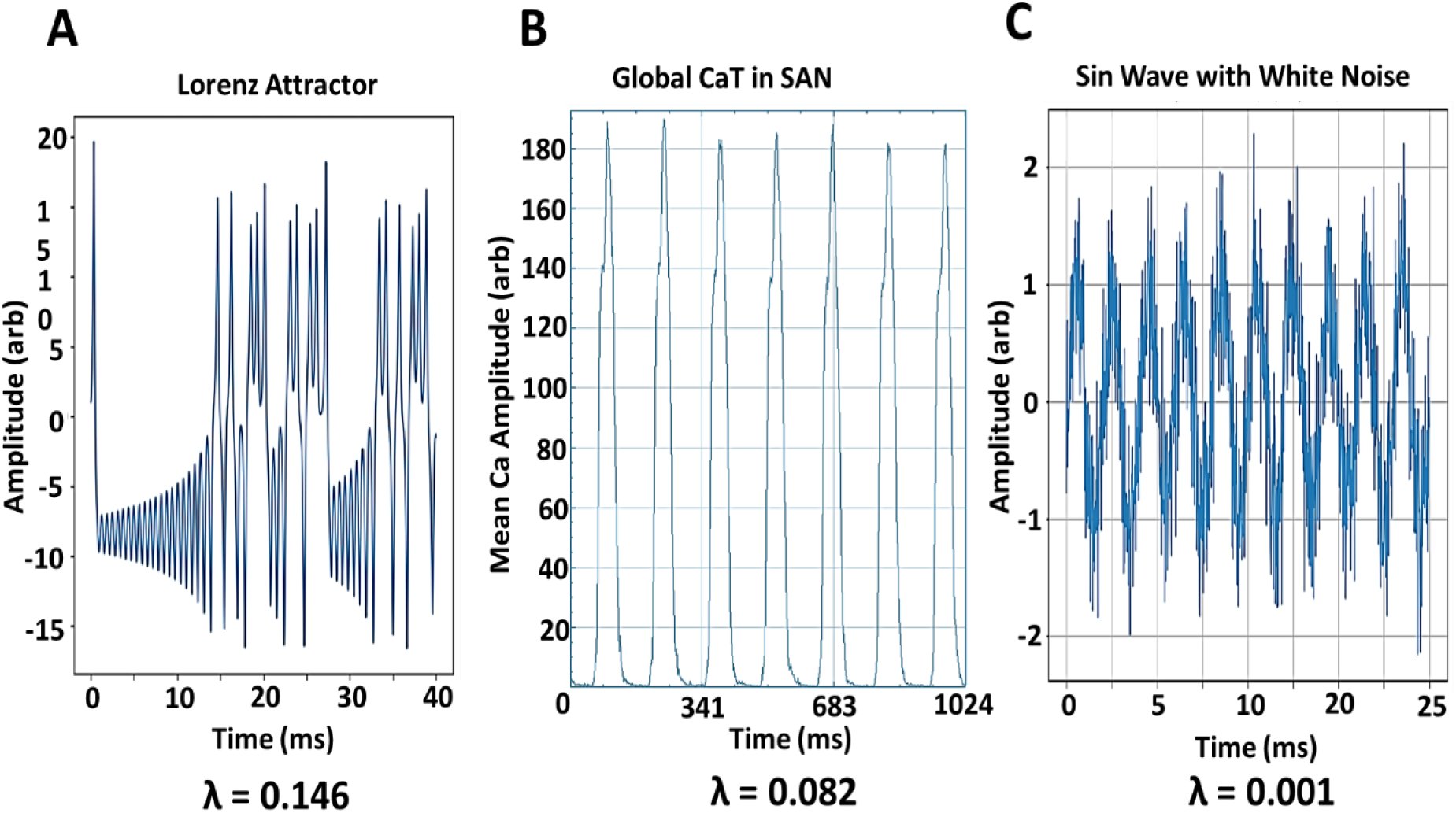
Positive Lyapunov exponent quantifies an intermediate level of chaos within the global SAN CaT. (A) Lorenz attractor, a classic example of deterministic chaos behavior (λ = 0.146); (B) The original recorded time series of global CaTs (λ = 0.082). (C) Noisy sinusoidal waves, representing a stochastic system (λ ≈ 0.0001); and (the reconstruction parameters included: embedding dimension = 2, lag = 1, minimum time separation = 4, tau = 1, and minimum neighbors = 10).

The SAN Ca^2+^ signaling network exhibits a Lyapunov exponent (λ = 0.082) intermediate between the strongly chaotic Lorenz attractor (λ = 0.146) and a noisy sinusoidal wave (λ ≈ 0.0001). This indicates moderate chaotic dynamics with sensitivity to initial conditions, but more structured than the Lorenz attractor and not purely random, complementing the results of the complex dynamics in stability and orbit plots (**Fig. 4, Figs. S13-S14**)

Analyses of the Lyapunov exponent (λ) for each of the four color-coded clusters outputted equal to λ = 0.0570, λ = 0.0531, λ = 0.0566, and λ = 0.040, for red, yellow, green, and blue clusters respectively. This means that small changes in the initial condition of the red cluster can influence the subsequent trajectory of the Ca^2+^ signal in time-series, infusing a variability signature into impulses in the timeseries. It is important to note that the Lyapunov exponent for integrated SAN Ca^2+^ dynamic (λ = 0.082) is greater than that of any given cluster, suggesting that deterministic behavior, in addition to stochastic behavior, is embedded within the SAN Ca^2+^ dynamic.

## References

1. Berridge MJ, Bootman MD, Roderick HL. Calcium signalling: Dynamics, homeostasis and remodelling. Nat Rev Mol Cell Bio. 2003;4(7):517–29. doi: 10.1038/nrm1155. PubMed PMID: WOS:000183892000011.

2. Berridge MJ, Lipp P, Bootman MD. The versatility and universality of calcium signalling. Nat Rev Mol Cell Bio. 2000;1(1):11–21. doi: Doi 10.1038/35036035. PubMed PMID: WOS:000165765100012.

3. Borrus DS, Stettler MK, Grover CJ, Kalajian EJ, Gu J, Smith GDC, Del Negro CA. Inspiratory and sigh breathing rhythms depend on distinct cellular signalling mechanisms in the preBotzinger complex. J Physiol-London. 2024;602(5):809–34. doi: 10.1113/Jp285582. PubMed PMID: WOS:001161691600001.

4. Stozer A, Dolensek J, Rupnik MS. Glucose-stimulated calcium dynamics in islets of Langerhans in acute mouse pancreas tissue slices. PLoS One. 2013;8(1):e54638. Epub 2013/01/30. doi: 10.1371/journal.pone.0054638. PubMed PMID: 23358454; PubMed Central PMCID: PMCPMC3554663.

5. Belote RL, Simon SM. Ca2+ transients in melanocyte dendrites and dendritic spine-like structures evoked by cell-to-cell signaling. J Cell Biol. 2020;219(1). doi: 10.1083/jcb.201902014. PubMed PMID: 31821412; PubMed Central PMCID: PMCPMC7039208.

6. Tyser RCV, Miranda AMA, Chen CM, Davidson SM, Srinivas S, Riley PR. Calcium handling precedes cardiac differentiation to initiate the first heartbeat. Elife. 2016;5. doi: ARTN e17113 10.7554/eLife.17113. PubMed PMID: WOS:000386050500001.

7. Jia BZ, Qi Y, Wong-Campos JD, Megason SG, Cohen AE. A bioelectrical phase transition patterns the first vertebrate heartbeats. Nature. 2023;622(7981):149-55. Epub 20230927. doi: 10.1038/s41586-023-06561-z. PubMed PMID: 37758945.

8. Donald L, Lakatta EG. What makes the sinoatrial node tick? A question not for the faint of heart. Philos Trans R Soc Lond B Biol Sci. 2023;378(1879):20220180. Epub 20230501. doi: 10.1098/rstb.2022.0180. PubMed PMID: 37122227; PubMed Central PMCID: PMCPMC10150214.

9. Lakatta EG, Maltsev VA, Vinogradova TM. A coupled SYSTEM of intracellular Ca2+ clocks and surface membrane voltage clocks controls the timekeeping mechanism of the heart’s pacemaker. Circ Res. 2010;106(4):659–73. doi: 10.1161/CIRCRESAHA.109.206078. PubMed PMID: 20203315; PubMed Central PMCID: PMCPMC2837285.

10. Maltsev VA, Vinogradova TM, Stern MD, Lakatta EG. Letter to the editor: “Validating the requirement for beat-to-beat coupling of the Ca2+ clock and M clock in pacemaker cell normal automaticity“. Am J Physiol Heart Circ Physiol. 2011;300(6):H2323-4; author reply H5-6. doi: 10.1152/ajpheart.00110.2011. PubMed PMID: 21632980; PubMed Central PMCID: PMCPMC3119099.

11. Vinogradova TM, Kobrinsky E, Lakatta EG. Dual Activation of Phosphodiesterases 3 and 4 Regulates Basal Spontaneous Beating Rate of Cardiac Pacemaker Cells: Role of Compartmentalization? Front Physiol. 2018;9:1301. Epub 20181009. doi: 10.3389/fphys.2018.01301. PubMed PMID: 30356755; PubMed Central PMCID: PMCPMC6189467.

12. Vinogradova TM, Lakatta EG. Regulation of basal and reserve cardiac pacemaker function by interactions of cAMP-mediated PKA-dependent Ca2+ cycling with surface membrane channels. J Mol Cell Cardiol. 2009;47(4):456–74. Epub 20090630. doi: 10.1016/j.yjmcc.2009.06.014. PubMed PMID: 19573534; PubMed Central PMCID: PMCPMC2757791.

13. Vinogradova TM, Lakatta EG. Dual Activation of Phosphodiesterase 3 and 4 Regulates Basal Cardiac Pacemaker Function and Beyond. Int J Mol Sci. 2021;22(16). Epub 20210805. doi: 10.3390/ijms22168414. PubMed PMID: 34445119; PubMed Central PMCID: PMCPMC8395138.

14. Vinogradova TM, Maltsev VA, Bogdanov KY, Lyashkov AE, Lakatta EG. Rhythmic Ca oscillations drive sinoatrial nodal cell pacemaker function to make the heart tick. Ann Ny Acad Sci. 2005;1047:138–56. doi: 10.1196/annals.1341.013. PubMed PMID: WOS:000231874400013.

15. Vinogradova TM, Tagirova Sirenko S, Lakatta EG. Unique Ca(2+)-Cycling Protein Abundance and Regulation Sustains Local Ca(2+) Releases and Spontaneous Firing of Rabbit Sinoatrial Node Cells. Int J Mol Sci. 2018;19(8). Epub 20180725. doi: 10.3390/ijms19082173. PubMed PMID: 30044420; PubMed Central PMCID: PMCPMC6121616.

16. Maltsev AV, Maltsev VA, Mikheev M, Maltseva LA, Sirenko SG, Lakatta EG, Stern MD. Synchronization of stochastic Ca(2)(+) release units creates a rhythmic Ca(2)(+) clock in cardiac pacemaker cells. Biophys J. 2011;100(2):271–83. doi: 10.1016/j.bpj.2010.11.081. PubMed PMID: 21244823; PubMed Central PMCID: PMCPMC3021664.

17. Bychkov R, Juhaszova M, Tsutsui K, Coletta C, Stern MD, Maltsev VA, Lakatta EG. Synchronized Cardiac Impulses Emerge From Heterogeneous Local Calcium Signals Within and Among Cells of Pacemaker Tissue. Jacc-Clin Electrophy. 2020;6(8):907–31. doi: 10.1016/j.jacep.2020.06.022. PubMed PMID: WOS:000602739500001.

18. Bychkov R, Juhaszova M, Calvo-Rubio Barrera M, Donald LAH, Coletta C, Shumaker C, Moorman K, Sirenko ST, Maltsev AV, Sollott SJ, Lakatta EG. The Heart’s Pacemaker Mimics Brain Cytoarchitecture and Function: Novel Interstitial Cells Expose Complexity of the SAN. JACC Clin Electrophysiol. 2022;8(10):1191–215. Epub 20220928. doi: 10.1016/j.jacep.2022.07.003. PubMed PMID: 36182566; PubMed Central PMCID: PMCPMC9665104.

19. Efimov IR, Nikolski VP, Salama G. Optical imaging of the heart. Circ Res. 2004;95(1):21–33. doi: 10.1161/01.RES.0000130529.18016.35. PubMed PMID: 15242982.

20. Lang D, Petrov V, Lou Q, Osipov G, Efimov IR. Spatiotemporal control of heart rate in a rabbit heart. J Electrocardiol. 2011;44(6):626–34. Epub 20110919. doi: 10.1016/j.jelectrocard.2011.08.010. PubMed PMID: 21937057; PubMed Central PMCID: PMCPMC3220935.

21. Li N, Hansen BJ, Csepe TA, Zhao J, Ignozzi AJ, Sul LV, Zakharkin SO, Kalyanasundaram A, Davis JP, Biesiadecki BJ, Kilic A, Janssen PML, Mohler PJ, Weiss R, Hummel JD, Fedorov VV. Redundant and diverse intranodal pacemakers and conduction pathways protect the human sinoatrial node from failure. Sci Transl Med. 2017;9(400). doi: 10.1126/scitranslmed.aam5607. PubMed PMID: 28747516; PubMed Central PMCID: PMCPMC5775890.

22. Faisal AA, Selen LP, Wolpert DM. Noise in the nervous system. Nat Rev Neurosci. 2008;9(4):292–303. doi: 10.1038/nrn2258. PubMed PMID: 18319728; PubMed Central PMCID: PMCPMC2631351.

23. Guo DQ, Perc M, Liu TJ, Yao DZ. Functional importance of noise in neuronal information processing. Epl-Europhys Lett. 2018;124(5). doi: Artn 50001 10.1209/0295-5075/124/50001. PubMed PMID: WOS:000454552700001.

24. Liu HX, Lu LL, Zhu Y, Wei ZC, Yi M. Stochastic resonance: The response to envelope modulation signal for neural networks with different topologies. Physica A. 2022;607. doi: ARTN 128177 10.1016/j.physa.2022.128177. PubMed PMID: WOS:000876411900009.

25. Gallos LK, Makse HA, Sigman M. A small world of weak ties provides optimal global integration of self-similar modules in functional brain networks. P Natl Acad Sci USA. 2012;109(8):2825–30. doi: 10.1073/pnas.1106612109. PubMed PMID: WOS:000300495100037.

26. Sporns O, Zwi JD. The small world of the cerebral cortex. Neuroinformatics. 2004;2(2):145–62. doi: Doi 10.1385/Ni:2:2:145. PubMed PMID: WOS:000223582200003.

27. Watts DJ, Strogatz SH. Collective dynamics of ‘small-world’ networks. Nature. 1998;393(6684):440-2. doi: Doi 10.1038/30918. PubMed PMID: WOS:000074020000035.

28. Qu ZL, Garfinkel A, Weiss JN, Nivala M. Multi-scale modeling in biology: How to bridge the gaps between scales? Prog Biophys Mol Bio. 2011;107(1):21–31. doi: 10.1016/j.pbiomolbio.2011.06.004. PubMed PMID: WOS:000296544000004.

29. Guarina L, Moghbel AN, Pourhosseinzadeh MS, Cudmore RH, Sato D, Clancy CE, Santana LF. Biological noise is a key determinant of the reproducibility and adaptability of cardiac pacemaking and EC coupling. J Gen Physiol. 2022;154(9). doi: ARTN e202012613 10.1085/jgp.202012613. PubMed PMID: WOS:000860454700001.

30. Xiong LI, Garfinkel A. Are physiological oscillations physiological? J Physiol. 2023. Epub 20230825. doi: 10.1113/JP285015. PubMed PMID: 37622389.

31. Abreu T, Silva PA, Sancho F, Temperville A. Analytical approximate wave form for asymmetric waves. Coast Eng. 2010;57(7):656–67. doi: 10.1016/j.coastaleng.2010.02.005. PubMed PMID: WOS:000278342200003.

32. Ramsay JO, Silverman BW. Functional data analysis. New York: Springer; 1997. xiv, 310 p. p.

33. Chen Y, Yang H. Multiscale recurrence analysis of long-term nonlinear and nonstationary time series. Chaos Soliton Fract. 2012;45(7):978–87. doi: 10.1016/j.chaos.2012.03.013. PubMed PMID: WOS:000306383000008.

34. Yang H. Multiscale recurrence quantification analysis of spatial cardiac vectorcardiogram signals. IEEE Trans Biomed Eng. 2011;58(2):339–47. Epub 2010/08/10. doi: 10.1109/TBME.2010.2063704. PubMed PMID: 20693104.

35. Meyers A, Buqammaz M, Yang H. Cross-recurrence analysis for pattern matching of multidimensional physiological signals. Chaos. 2020;30(12):123125. Epub 2021/01/01. doi: 10.1063/5.0030838. PubMed PMID: 33380053.

36. Jalife J. Mutual Entrainment and Electrical Coupling as Mechanisms for Synchronous Firing of Rabbit Sino-Atrial Pace-Maker Cells. J Physiol-London. 1984;356(Nov):221–43. doi: DOI 10.1113/jphysiol.1984.sp015461. PubMed PMID: WOS:A1984TT18900014.

37. Michaels DC, Chialvo DR, Matyas EP, Jalife J. Chaotic activity in a mathematical model of the vagally driven sinoatrial node. Circ Res. 1989;65(5):1350–60. doi: 10.1161/01.res.65.5.1350. PubMed PMID: 2805248.

38. Packard NH, Crutchfield JP, Farmer JD, Shaw RS. Geometry from a Time-Series. Phys Rev Lett. 1980;45(9):712–6. doi: DOI 10.1103/PhysRevLett.45.712. PubMed PMID: WOS:A1980KE58800012.

39. Lyashkov AE, Beahr J, Lakatta EG, Yaniv Y, Maltsev VA. Positive Feedback Mechanisms among Local Ca Releases, NCX, and I Ignite Pacemaker Action Potentials. Biophys J. 2018;114(5):1176–89. doi: 10.1016/j.bpj.2017.12.043. PubMed PMID: WOS:000428017500020.

40. Fontes MS, van Veen TA, de Bakker JM, van Rijen HV. Functional consequences of abnormal Cx43 expression in the heart. Biochim Biophys Acta. 2012;1818(8):2020–9. Epub 2011/08/16. doi: 10.1016/j.bbamem.2011.07.039. PubMed PMID: 21839722.

41. Bogdanov KY, Maltsev VA, Vinogradova TM, Lyashkov AE, Spurgeon HA, Stern MD, Lakatta EG. Membrane potential fluctuations resulting from submembrane Ca2+ releases in rabbit sinoatrial nodal cells impart an exponential phase to the late diastolic depolarization that controls their chronotropic state. Circ Res. 2006;99(9):979–87. Epub 20060928. doi: 10.1161/01.RES.0000247933.66532.0b. PubMed PMID: 17008599.

42. Tagirova Sirenko S, Tsutsui K, Tarasov KV, Yang D, Wirth AN, Maltsev VA, Ziman BD, Yaniv Y, Lakatta EG. Self-Similar Synchronization of Calcium and Membrane Potential Transitions During Action Potential Cycles Predict Heart Rate Across Species. JACC Clin Electrophysiol. 2021;7(11):1331–44. Epub 20210428. doi: 10.1016/j.jacep.2021.02.016. PubMed PMID: 33933406; PubMed Central PMCID: PMCPMC10089231.

43. Vinogradova TM, Lyashkov AE, Zhu W, Ruknudin AM, Sirenko S, Yang D, Deo S, Barlow M, Johnson S, Caffrey JL, Zhou YY, Xiao RP, Cheng H, Stern MD, Maltsev VA, Lakatta EG. High basal protein kinase A-dependent phosphorylation drives rhythmic internal Ca2+ store oscillations and spontaneous beating of cardiac pacemaker cells. Circ Res. 2006;98(4):505–14. Epub 20060119. doi: 10.1161/01.RES.0000204575.94040.d1. PubMed PMID: 16424365.

44. Vinogradova TM, Maltsev VA, Bogdanov KY, Lyashkov AE, Lakatta EG. Rhythmic Ca2+ oscillations drive sinoatrial nodal cell pacemaker function to make the heart tick. Ann N Y Acad Sci. 2005;1047:138–56. doi: 10.1196/annals.1341.013. PubMed PMID: 16093492.

45. Yang D, Morrell CH, Lyashkov AE, Tagirova Sirenko S, Zahanich I, Yaniv Y, Vinogradova TM, Ziman BD, Maltsev VA, Lakatta EG. Ca(2+) and Membrane Potential Transitions During Action Potentials Are Self-Similar to Each Other and to Variability of AP Firing Intervals Across the Broad Physiologic Range of AP Intervals During Autonomic Receptor Stimulation. Front Physiol. 2021;12:612770. Epub 20210908. doi: 10.3389/fphys.2021.612770. PubMed PMID: 34566668; PubMed Central PMCID: PMCPMC8456031.

46. Ben David S, Ho KYL, Tanentzapf G, Zaritsky A. Formation of recurring transient Ca2+-based intercellular communities during hematopoiesis. P Natl Acad Sci USA. 2024;121(16). doi: ARTN e231815512110.1073/pnas.2318155121. PubMed PMID: WOS:001209473100001.

47. Del Negro CA, Hayes JA, Rekling JC. Dendritic calcium activity precedes inspiratory bursts in preBotzinger complex neurons. J Neurosci. 2011;31(3):1017–22. Epub 2011/01/21. doi: 10.1523/JNEUROSCI.4731-10.2011. PubMed PMID: 21248126; PubMed Central PMCID: PMCPMC3075810.

48. Pedroni A, Yilmaz E, Del Vecchio L, Bhattarai P, Vidal IT, Dai YE, Koutsogiannis K, Kizil C, Ampatzis K. Decoding the molecular, cellular, and functional heterogeneity of zebrafish intracardiac nervous system. Nat Commun. 2024;15(1):10483. Epub 2024/12/05. doi: 10.1038/s41467-024-54830-w. PubMed PMID: 39632839; PubMed Central PMCID: PMCPMC11618350.

49. McMillen P, Levin M. Collective intelligence: A unifying concept for integrating biology across scales and substrates. Commun Biol. 2024;7(1):378. Epub 2024/03/29. doi: 10.1038/s42003-024-06037-4. PubMed PMID: 38548821; PubMed Central PMCID: PMCPMC10978875 sponsored research agreement with a company, Astonishing Labs, to fund projects relevant to the collective intelligence of cells.

50. Gallos LK, Makse HA, Sigman M. A small world of weak ties provides optimal global integration of self-similar modules in functional brain networks. Proc Natl Acad Sci U S A. 2012;109(8):2825–30. Epub 2012/02/07. doi: 10.1073/pnas.1106612109. PubMed PMID: 22308319; PubMed Central PMCID: PMCPMC3286928.

51. Wilson EO. Consilience : the unity of knowledge. 1st ed. New York: Knopf : Distributed by Random House; 1998. 332 p. p.

52. Facchin F, Canaider S, Tassinari R, Zannini C, Bianconi E, Taglioli V, Olivi E, Cavallini C, Tausel M, Ventura C. Physical energies to the rescue of damaged tissues. World J Stem Cells. 2019;11(6):297–321. doi: 10.4252/wjsc.v11.i6.297. PubMed PMID: WOS:000472964200002.

53. Fedorov VV, Chang R, Glukhov AV, Kostecki G, Janks D, Schuessler RB, Efimov IR. Complex Interactions Between the Sinoatrial Node and Atrium During Reentrant Arrhythmias in the Canine Heart. Circulation. 2010;122(8):782–U79. doi: 10.1161/Circulationaha.109.935288. PubMed PMID: WOS:000281193100005.

54. Glukhov AV, Fedorov VV, Kalish PW, Ravikumar VK, Lou Q, Janks D, Schuessler RB, Moazami N, Efimov IR. Conduction Remodeling in Human End-Stage Nonischemic Left Ventricular Cardiomyopathy. Circulation. 2012;125(15):1835-+. doi: 10.1161/Circulationaha.111.047274. PubMed PMID: WOS:000302957800011.

55. Boyett MR, Honjo H, Kodama I. The sinoatrial node, a heterogeneous pacemaker structure. Cardiovasc Res. 2000;47(4):658–87. doi: 10.1016/s0008-6363(00)00135-8. PubMed PMID: 10974216.

56. Bouman LN, Jongsma HJ. Structure and function of the sino-atrial node: a review. Eur Heart J. 1986;7(2):94–104. doi: 10.1093/oxfordjournals.eurheartj.a062047. PubMed PMID: 2422036.

57. Fedorov VV, Glukhov AV, Chang R. Conduction barriers and pathways of the sinoatrial pacemaker complex: their role in normal rhythm and atrial arrhythmias. Am J Physiol Heart Circ Physiol. 2012;302(9):H1773–83. Epub 20120120. doi: 10.1152/ajpheart.00892.2011. PubMed PMID: 22268110.

58. Verheijck EE, van Kempen MJA, Veereschild M, Lurvink J, Jongsma HJ, Bouman LN. Electrophysiological features of the mouse sinoatrial node in relation to connexin distribution. Cardiovasc Res. 2001;52(1):40–50. doi: Doi 10.1016/S0008-6363(01)00364-9. PubMed PMID: WOS:000171953400006.

59. Boyett MR. ‘And the beat goes on’ The cardiac conduction system: the wiring system of the heart. Exp Physiol. 2009;94(10):1035–49. doi: 10.1113/expphysiol.2009.046920. PubMed PMID: WOS:000269866400001.

60. Chen WT, Chen YC, Lu YY, Kao YH, Huang JH, Lin YK, Chen SA, Chen YJ. Apamin modulates electrophysiological characteristics of the pulmonary vein and the Sinoatrial Node. Eur J Clin Invest. 2013;43(9):957–63. doi: 10.1111/eci.12125. PubMed PMID: WOS:000323152800008.

61. Huang JH. Apamin modulates electrophysiological characteristics of the pulmonary vein and the Sinoatrial Node (vol 43, pg 957, 2013). Eur J Clin Invest. 2016;46(7):675-. doi: 10.1111/eci.12645. PubMed PMID: WOS:000379669500008.

62. Torrente AG, Zhang R, Wang HD, Zaini A, Kim B, Yue X, Philipson KD, Goldhaber JI. Contribution of small conductance K channels to sinoatrial node pacemaker activity: insights from atrial-specific Na/Ca exchange knockout mice. J Physiol-London. 2017;595(12):3847–65. doi: 10.1113/Jp274249. PubMed PMID: WOS:000404654500015.

63. Sirenko ST, Zahanich I, Li Y, Lukyanenko YO, Lyashkov AE, Ziman BD, Tarasov KV, Younes A, Riordon DR, Tarasova YS, Yang D, Vinogradova TM, Maltsev VA, Lakatta EG. Phosphoprotein Phosphatase 1 but Not 2A Activity Modulates Coupled-Clock Mechanisms to Impact on Intrinsic Automaticity of Sinoatrial Nodal Pacemaker Cells. Cells. 2021;10(11). Epub 2021/11/28. doi: 10.3390/cells10113106. PubMed PMID: 34831329; PubMed Central PMCID: PMCPMC8623309.

64. Allessie MA, Bonke FI. Direct demonstration of sinus node reentry in the rabbit heart. Circ Res. 1979;44(4):557–68. doi: 10.1161/01.res.44.4.557. PubMed PMID: 428051.

65. Allessie MA, Bonke FIM, Kirchhof CJHJ. Electrotonic Influence of Atrial Myocardium on Sinus Node Function in the Rabbit. J Physiol-London. 1985;366(Sep):P36-P. PubMed PMID: WOS:A1985ARF5900060.

66. Bachmann G. The Inter-Auricular Time Interval. Am J Physiol. 1916;41(3):309–20. doi: DOI 10.1152/ajplegacy.1916.41.3.309. PubMed PMID: WOS:000207509400004.

67. Mori S, Shivkumar K. Atlas of Cardiac Anatomy Cardiotext; 2022.

68. Chhabra L, Devadoss R, Chaubey VK, Spodick DH. Interatrial Block in the Modern Era. Curr Cardiol Rev. 2014;10(3):181–9. doi: 10.2174/1573403×10666140514101748. PubMed PMID: WOS:000212837200003.

69. Nitta T, Ishii Y, Miyagi Y, Ohmori H, Sakamoto S, Tanaka S. Concurrent multiple left atrial focal activations with fibrillatory conduction and right atrial focal or reentrant activation as the mechanism in atrial fibrillation. J Thorac Cardiov Sur. 2004;127(3):770–8. doi: 10.1016/j.jtcvs.2003.05.001. PubMed PMID: WOS:000220115400025.

70. Swartz MF, Fink GW, Lutz CJ, Taffet SM, Berenfeld O, Vikstrom KL, Kasprowicz K, Bhatta L, Puskas F, Kalifa J, Jalife J. Left versus right atrial difference in dominant frequency, K channel transcripts, and fibrosis in patients developing atrial fibrillation after cardiac surgery. Heart Rhythm. 2009;6(10):1415–22. doi: 10.1016/j.hrthm.2009.06.018. PubMed PMID: WOS:000270502700003.

71. Jalife J, Moe GK. Effect of Electrotonic Potentials on Pacemaker Activity of Canine Purkinje-Fibers in Relation to Parasystole. Circulation Research. 1976;39(6):801–8. doi: Doi 10.1161/01.Res.39.6.801. PubMed PMID: WOS:A1976CP45700008.

## References

1. Bychkov R, Juhaszova M, Tsutsui K, Coletta C, Stern MD, Maltsev VA, Lakatta EG. Synchronized Cardiac Impulses Emerge From Heterogeneous Local Calcium Signals Within and Among Cells of Pacemaker Tissue. Jacc-Clin Electrophy. Aug 2020;6(8):907–931. doi:10.1016/j.jacep.2020.06.022

2. Bergé P, 1934-, Pomeau Y, Vidal CC. Order within chaos : towards a deterministic approach to turbulence. New York : Wiley ; Paris : Herman; 1986.

3. Packard NH, Crutchfield JP, Farmer JD, Shaw RS. Geometry from a Time-Series. Phys Rev Lett. 1980;45(9):712–716. doi:DOI 10.1103/PhysRevLett.45.712

4. Chen Y, Yang H. Multiscale recurrence analysis of long-term nonlinear and nonstationary time series. Chaos Soliton Fract. Jul 2012;45(7):978–987. doi:10.1016/j.chaos.2012.03.013

5. Yang H. Multiscale recurrence quantification analysis of spatial cardiac vectorcardiogram signals. IEEE Trans Biomed Eng. Feb 2011;58(2):339–47. doi:10.1109/TBME.2010.2063704

6. Abreu T, Silva PA, Sancho F, Temperville A. Analytical approximate wave form for asymmetric waves. Coast Eng. Jul 2010;57(7):656–667. doi:10.1016/j.coastaleng.2010.02.005

7. Huang NE. Introduction to the Hilbert-Huang Transform and Its Related Mathematical Problems. Interd Math Sci. 2005;5:1–26. doi:Book_Doi 10.1142/9789812703347

8. Mateo C, Talavera JA. Short-time Fourier transform with the window size fixed in the frequency domain. Digital Signal Processing. 2018;77:13–21.

9. Meyers A, Buqammaz M, Yang H. Cross-recurrence analysis for pattern matching of multidimensional physiological signals. Chaos. Dec 2020;30(12):123125. doi:10.1063/5.0030838

10. Ramsay JO, Silverman BW. Functional data analysis. Springer series in statistics. Springer; 1997:xiv, 310 p.

11. Jalife J. Mutual Entrainment and Electrical Coupling as Mechanisms for Synchronous Firing of Rabbit Sino-Atrial Pace-Maker Cells. J Physiol-London. 1984;356(Nov):221–243. doi:DOI 10.1113/jphysiol.1984.sp015461

